# Scalable generalized linear mixed model for region-based association tests in large biobanks and cohorts

**DOI:** 10.1101/583278

**Authors:** Wei Zhou, Zhangchen Zhao, Jonas B. Nielsen, Lars G. Fritsche, Jonathon LeFaive, Sarah A. Gagliano Taliun, Wenjian Bi, Maiken E. Gabrielsen, Mark J. Daly, Benjamin M. Neale, Kristian Hveem, Goncalo R. Abecasis, Cristen J. Willer, Seunggeun Lee

## Abstract

With very large sample sizes, population-based cohorts and biobanks provide an exciting opportunity to identify genetic components of complex traits. To analyze rare variants, gene or region-based multiple variant aggregate tests are commonly used to increase association test power. However, due to the substantial computation cost, existing region-based rare variant tests cannot analyze hundreds of thousands of samples while accounting for confounders, such as population stratification and sample relatedness. Here we propose a scalable generalized mixed model region-based association test that can handle large sample sizes and accounts for unbalanced case-control ratios for binary traits. This method, SAIGE-GENE, utilizes state-of-the-art optimization strategies to reduce computational and memory cost, and hence is applicable to exome-wide and genome-wide region-based analysis for hundreds of thousands of samples. Through the analysis of the HUNT study of 69,716 Norwegian samples and the UK Biobank data of 408,910 White British samples, we show that SAIGE-GENE can efficiently analyze large sample data (N > 400,000) with type I error rates well controlled.

## Introduction

In recent years, large cohort studies and biobanks, such as Trans-Omics for Precision Medicine (TOPMed) study^1^ and UK Biobank^2^, have sequenced or genotyped hundreds of thousands of samples, which are invaluable resources to identify genetic components of complex traits, including rare variants (minor allele frequency (MAF) <1%). It is well known that single variant tests are underpowered to identify trait-associated rare variants’. Gene- or region-based tests, such as Burden test, SKAT’ and SKAT-O^5^, can be more powerful by grouping rare variants into functional units, i.e. genes. To adjust for both population structure and sample relatedness, gene-based tests have been extended to mixed models^6^. For example, EmmaX^7^ based SKAT’ approaches (EmmaX-SKAT) have been implemented and used for many rare variant association studies including TOPMed^1,8^. The generalized linear mixed model gene-based test, SMMAT, has been recently developed^6^. However, these approaches require O(*N*^3^) computation time and O(*N*^2^) memory usages, where *N* is the sample size, which are not scalable to large datasets.

Here, we propose a novel method called SAIGE-GENE for region-based association analysis that is capable of handling very large samples(>400,000 individuals), while inferring and accounting for sample relatedness. SAIGE-GENE is an extension of the previously developed single variant association method, SAIGE^9^, with a modification suitable to rare variants. Same as SAIGE, it utilizes state-of-the-art optimization strategies to reduce computation cost for fitting null mixed models. To ensure computation efficiency while improving test accuracy for rare variants, SAIGE-GENE approximates the variance of score statistics calculated with the full genetic relationship matrix (GRM) using the variance calculated with a sparse GRM and the ratios of these two variances estimated from a subset of genetic markers. Because the sparse GRM, which is constructed by thresholding small values in the full GRM, preserves close family structures, this approach provides a far more accurate variance estimation for very rare variants (minor allele count (MAC) < 20) than the original approach in SAIGE^9^. By combining single variant score statistics, SAIGE-GENE can perform Burden, SKAT and SKAT-O type gene-based tests. We have also developed conditional analysis to perform association tests conditioning on a single variant or multiple variants to identify independent rare variant association signals. Furthermore, SAIGE-GENE can account for unbalanced case-control ratios of binary traits by adopting a robust adjustment based on saddlepoint approximation^10–12^ (SPA) and efficient resampling^13^ (ER). The robust adjustment was previously developed for independent samples^14^ and we have extended it for related samples in SAIGE-GENE.

We have demonstrated that SAIGE-GENE controls for type I error rates in related samples for both quantitative and binary traits through extensive simulations as well as real data analysis, including the HUNT study for 69,716 Norwegian samples^15,16^ and the UK Biobank for 408,910 White British samples^2^. By evaluating its computation performance of SAIGE-GENE, we have shown its feasibility for large-scale genome-wide analysis. To perform exome-wide gene-based tests on 400,000 samples with on average 50 markers per gene, SAIGE-GENE requires 2,238 CPU hours and less than 36 Gb memory, while current methods will cost more than > 10 Tb in memory. We have further applied SAIGE-GENE to 53 quantitative traits and 10 binary traits in the UK Biobank and identified several significantly associated genes through exome-wide gene-based tests.

## RESULTS

### Overview of Methods

SAIGE-GENE consists of two main steps: 1. Fitting the null generalized linear mixed model (GLMM) to estimate variance components and other model parameters. 2. Testing for association between each genetic variant set, such as a gene or a region, and the phenotype. Three different association tests: Burden, SKAT, and SKAT-O have been implemented in SAIGE-GENE. The workflow is shown in the **Supplementary Figure 1**.

SAIGE-GENE uses similar optimization strategies as utilized in the original SAIGE to achieve the scalability for fitting the null GLMM and estimating the model parameters in Step 1. In particular, the spectral decomposition has been replaced by the preconditioning conjugate gradient (PCG) to solve linear systems without calculating and inverting the *N* × *N* GRM. To reduce the memory usage, raw genotypes are stored in a binary vector and elements of GRM are calculated when needed rather than being stored.

One of the most time-consuming part in association tests is to calculate variance of single variant score statistic, which requires O(*N*^2^) computation. To reduce computation cost, existing approaches, such as SAIGE^9^, BOLT-LMM^17^, and GRAMMA-Gamma^18^, approximate the variance of single variant score statistics with the full GRM using the variance estimate without a GRM and the ratio of these two variances. The ratio, which is assumed to be constant, is estimated using a subset of randomly selected genetic markers. However, for very rare variants with MAC below 20, the constant ratio assumption is not satisfied (**Supplementary Figure 2, left panel**). This is because rare variants are more susceptible to close family structures. Thus, to better approximate the variance, SAIGE-GENE incorporates close family structures through a sparse GRM, in which GRM elements below a user-specified relatedness coefficient are zeroed out and close family structures are preserved. The ratio between the variance with the full GRM and with the sparse GRM is much less variable (**Supplementary Figure 2, right panel**). To construct a sparse GRM, a small subset of randomly selected genetic markers, i.e. 2,000, are firstly used to quickly estimate which sample pairs pass the user-specified coefficient of relatedness cutoff, e.g. ≥ 0.125 for up to 3^rd^ degree relatives. Then the coefficients of relatedness for those related pairs are further estimated using the full set of genetic markers, which equal to values in the full GRM. Given that estimated values for variance ratios vary by MAC for the extremely rare variants (**Supplementary Figure 2, left panel**), such as singletons and doubletons, the variance ratios need to be estimated separately for different MAC categories. By default, MAC categories are set to be MAC equals to 1,2,3,4,5,6 to 10, 11 to 20, and > 20.

In Step 2, gene-based tests are conducted using single variant score statistics and their covariance estimates, which are approximated as the product of the covariance with the sparse GRM and the pre-estimated ratio. SAIGE-GENE can carry out Burden, SKAT, and SKAT-O approaches. Since SKAT-O is a combined test of Burden and SKAT, and hence provides a robust power, SAIGE-GENE performs SKAT-O by default.

If a gene or a region is significantly associated with the phenotype of interest, it is necessary to test if the signal is from rare variants or just a shadow of common variants in the same locus. We have developed conditional analysis using linkage disequilibrium (LD) information between conditioning markers and the tested gene^19^. Details are described in the Online Methods section.

SAIGE-GENE uses the same generalized linear mixed model as in SMMAT, while SMMAT calculates the variances of the score statistics for all tested genes using the full GRM directly and hence can be thought of as the “exact” method. When the trait is continuous, GLMM used by SAIGE-GENE and SMMAT is equivalent to the linear mixed model (LMM) of EmmaX-SKAT. We have further shown that SAIGE-GENE provides consistent association p-values to the two “exact” methods, EmmaX-SKAT and SMMAT (r^2^ of - log_10_ p-values > 0.99), using both simulation studies (**Supplementary Figure 3**) and real data analysis for down-sampled UK Biobank and HUNT (**Supplementary Figure 4**), but with much smaller computation and memory cost (**Figure 1**). We have also shown that SAIGE-GENE with different coefficient of relatedness cutoffs (0.125 and 0.2) produced nearly identical association p-values for automated read pulse rates in UK Biobank (**Supplementary Figure 5**).

**Figure 1.**
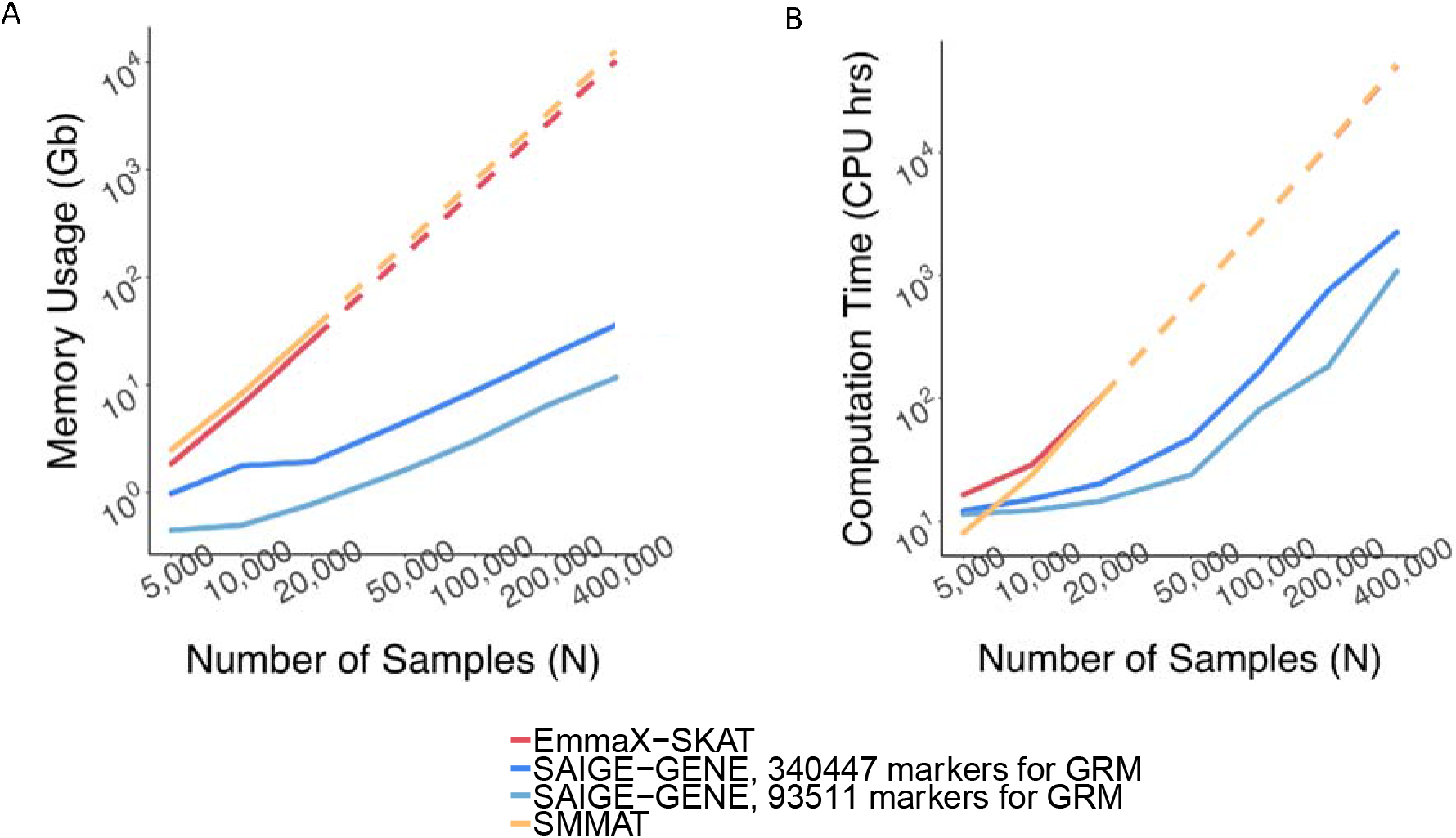
Estimated and projected computation cost by sample sizes (N) for gene-based tests for 15,342 genes, each containing 50 rare variants. Benchmarking was performed on randomly sub-sampled UK Biobank data with 408,144 White British participants for waist-to-hip ratio. The reported run times and memory are medians of five runs with samples randomly selected from the full sample set using different sampling seeds. There ported computation time and memory for EmmaX-SKAT and SMMAT is the projected computation time when N > 20,000. A. Log-log plots of the memory usage as a function of sample size (N) B. Log-log plots of the run time as a function of sample size (N). Numerical data are provided in **Supplementary Table 1**.

For binary phenotypes with unbalanced case-control ratios (<1:9), single variant score statistics do not follow the normal distribution, leading to inflated type I error rates for region-based test:^13^. To address this problem, we have recently developed a scalable robust adjustment for independent samples^14^. The approach uses saddlepoint approximation^10–12^ (SPA) and efficient resampling^13^ (ER) to calibrate the variance of single variant score statistics. We have extended this approach to GLMM for SAIGE-GENE, which provides greatly improved type I error control than the unadjusted approach of assuming normality (**Supplementary Figure 6**). Details can be found in **Supplementary Materials 1.3.3.**

### Computation and Memory Cost

To evaluate the computation performance of SAiGE-GENE, we randomly sampled subsets of the 408,144 UK Biobank participants with the White British ancestry and non-missing measurements for waist hip ratio^2^. We benchmarked SAIGE-GENE, EmmaX-SKAT, and SMMAT for exome-wide gene-based SKAT-O tests, in which 15,342 genes were tested with assuming that each has 50 rare variants.

Memory usage is plotted on a log10 scale against sample sizes in **Figure 1A**. The memory cost of SAIGE-GENE is linear to the number of markers, *M*_*1*_, used for kinship estimation, but using too few markers may not be sufficient to account for subtle sample relatedness in the data, leading to inflated type I error rates in genetic association tests^9,20^. SAIGE-GENE uses 11.74 Gb with *M*_*1*_ = 93,511 and 35.59 Gb when *M*_*1*_ = 340,447 when the sample size *N* is 400,000, making it feasible for large sample data. In contrast, with *N* = 400,000 the memory usages in EmmaX-SKAT and SMMAT are projected to be nearly 10Tb, which makes them impossible to be used for large sample data.

Total computation time for exome-wide gene-based tests is plotted on a log10 scale against the sample size as shown in **Figure 1B**. Computation time for Step 1 and Step 2 are plotted separately in **Supplementary Figure 7** with numbers presented in **Supplementary Table 1**. The computation time for Step 1 in SAIGE-GENE is approximately *O*(*M*_*1*_*N*^1.5^) and in SMMAT and EmmaX-SKAT is O(N^3^), where *M*_*1*_ is the number of markers used for estimating the full GRM and *N* is the sample size. In Step 2, the association test for each gene costs *O*(*qK*) in SAIGE-GENE, where *q* is the number of markers in the gene and *K* is the number of non-zero elements in the sparse GRM. Compared to *O*(*qN*^2^) in Step 2 of SMMAT and EmmaX-SKAT, SAIGE-GENE decreases the computation time dramatically. For example, in the UK Biobank (N =408,910) with the relatedness coefficient ≥ 0.125 (corresponding to preserving samples with 3^rd^ degree or closer relatives in the GRM), *K* = 493,536, which is the same order of magnitude of *N*, and hence *O*(*qK*) is greatly smaller than *O*(*qN*^2^). As the computation time in Step 2 is approximately linear to *q*, the number of markers in each variant set, the total computation time for exome-wide gene-based tests was projected by different *q* and plotted in **Supplementary Figure 8**. In addition, we plotted the projected computation time for genome-wide region-based tests against the sample size as shown in **Supplementary Figure 9**, in which 286,000 chunks with 50 markers per chunk were assumed to be tested, corresponding to 14.3 million markers in HRC-imputed UK Biobank data with MAF ≤ 1% and imputation info score ≥ 0.8.

With *M*_*1*_ = 340,447, it takes SAIGE-GENE 2,238 CPU hours for exome-wide gene-based tests and 3,919 CPU hours for genome-wide region-based tests for waist hip ratio with *N* = 400,000 and each test contains 50 markers on average. Compared to EmmaX-SKAT and SMMAT, SAIGE-GENE is 25 times faster for exome-wide gene-based tests and 161 times faster for genome-wide region-based tests. More details about the computation cost are presented in **Supplementary Table 1**.

To evaluate whether the additional steps in the robust adjustment for binary traits increases computation cost, we have obtained computation time of SAIGE-GENE with and without the adjustment when analyzing the UK Biobank data for glaucoma (PheCode:365). Samples were randomly selected from 4,462 glaucoma cases and 397,701 controls respectively, so the case-control ratio remained the same in sub-sampled data sets. The results are presented in **Supplementary Table 2** and plotted in **Supplementary Figure 10**, showing that the robust adjustment only slightly increases the computation cost (1,269 vs 1,232 CPU hours for exome-wide analysis with *M*_*1*_ = 93,511) compared to the unadjusted approach.

The computation time for constructing the sparse GRM is 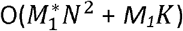. 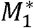 is the number of a small set of markers used for initial determination of related sample pairs based on a relationship coefficient cutoff, which by default is set to be 2,000. This step is only needed for each data set for one time to create a sparse GRM and the constructed sparse GRM will be re-used for all phenotypes in the same cohort or biobank. For example, for the UK Biobank with *N* = 408,910, *M*_*1*_= 340,447, 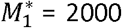, *K* = 493,536 with the relationship coefficient ≥ 0.125, corresponding to up to 3^rd^ degree relatives, it took 312 CPU hours to create the sparse GRM. Parallel computation is allowed for this step.

### Gene-based association analysis of quantitative traits in HUNT and UK Biobank

We applied SAIGE-GENE to analyze 13,416 genes, with at least two rare (MAF ≤ 1%) missense and stop-gain variants that were directly genotyped or imputed from HRC for high-density lipoprotein (HDL) in 69,716 Norwegian samples from a population-based Nord Trøndelag Health Study (HUNT)^9^. The HUNT study has substantial sample relatedness, in which ~55,000 samples have at least one up to 3rd degree relatives. The quantile-quantile (QQ) plot for the p-values of SKAT-O tests from SAIGE-GENE for HDL in HUNT is shown **Figure 2A**. As **Table 1** shows, eight genes reached the exome-wide significant threshold (p-value ≤ 2.5×10^−6^) and all of them are located in the previously reported GWAS loci for HDL^21,22^. By extending 500kb up and down stream, a top significant hit from single-variant association tests has been identified around each gene. For genes *UPC*, *LIPG*, *NR1H3*, and *CKAP5*, the top hits are common variants with MAF > 5% and the top hits in *FSD1L*, *ABCA1* and *RNF111* are less frequent non-coding variants that are not included in the gene-based tests. After conditioning on top hits, all genes, except for *FSD1L*, remained exome-wide significant, suggesting that SAIGE-GENE has identified associations of rare coding variants of those genes that are independent from the nearby association signals, pointing to candidate causal genes at those loci.

**Figure 2.**
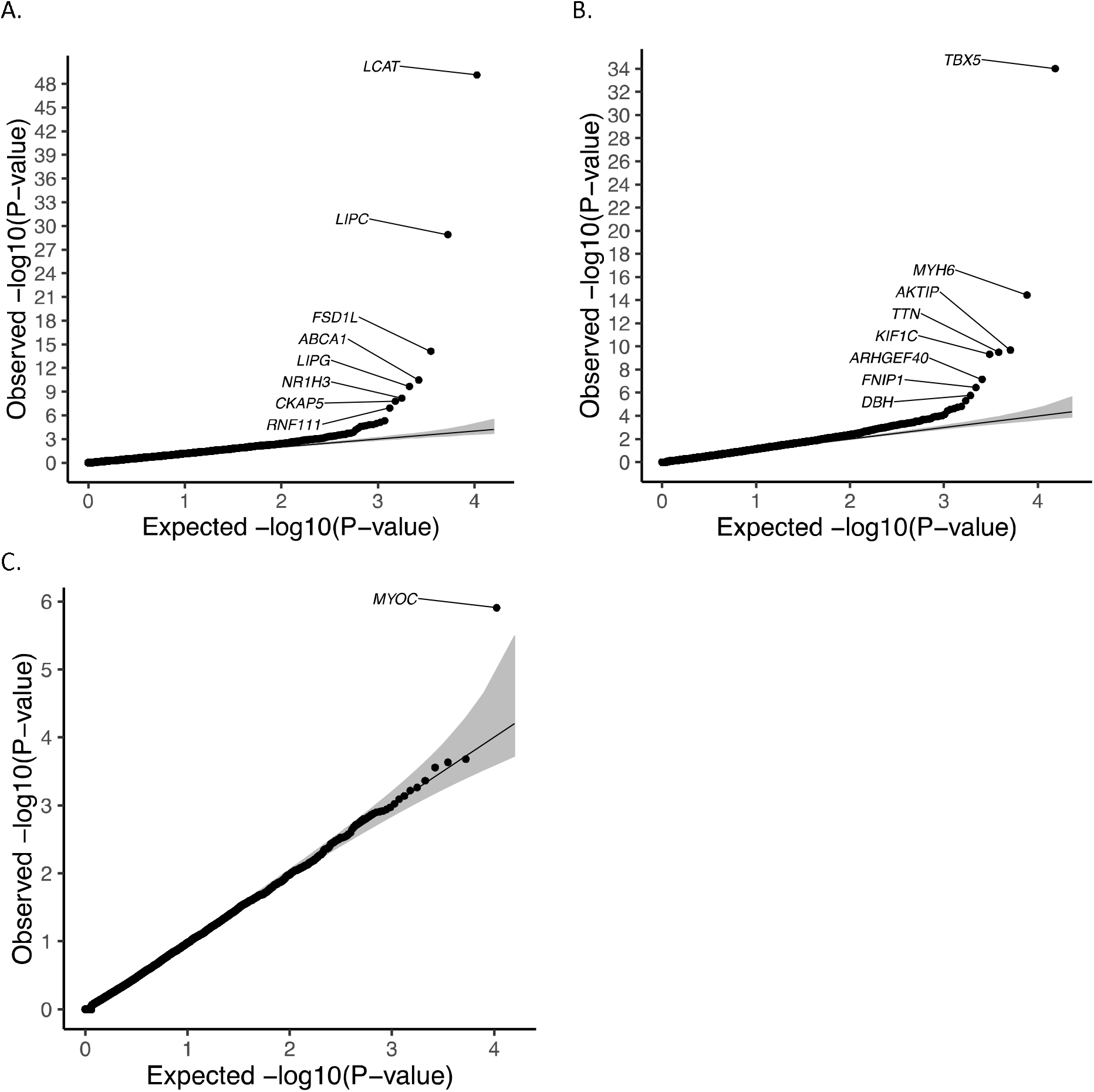
Quantile-quantile plots of exome-wide gene-based association results for A. high-density lipoprotein (HDL) in the HUNT study (N = 69,214). SKAT-O approach in SAIGE-GENE was performed for 13,416 genes with stop-gain and missense variants with MAF ≤ 1%, of which 10,600 having at least two variants are plotted. B. automated read pulse rate in the UK Biobank N = 385,365). C. glaucoma in the UK Biobank (N cases = 4,462; N controls = 397,761). SKAT-O approach in SAIGE-GENE was performed for 15,338 genes with stop-gain and missense variants with MAF ≤ 1%, of which 12,638 having at least two variants are plotting.

**Table 1.**
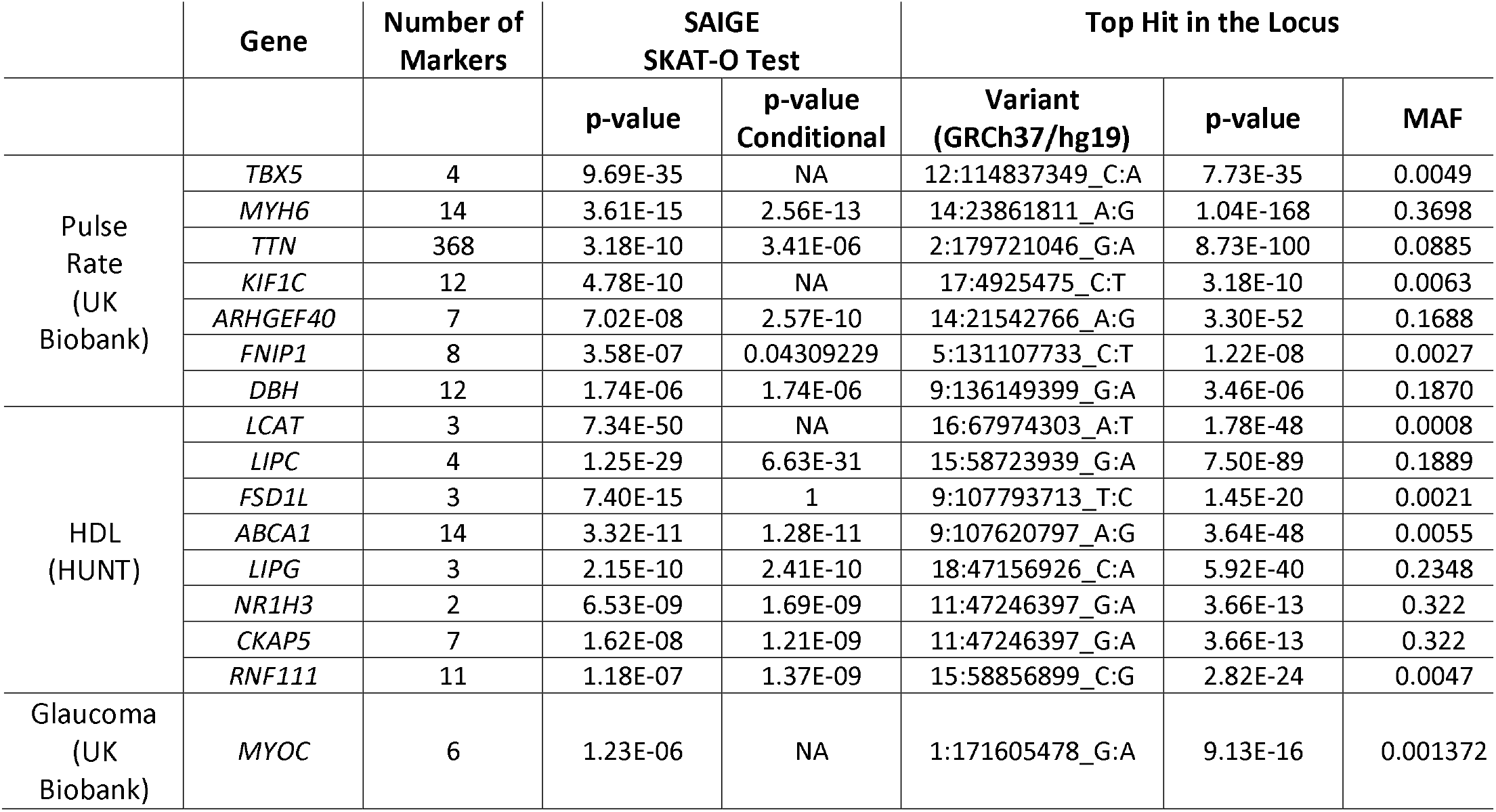
Genes that are significantly associated with automated read pulse rate and glaucoma in the UK Biobank and high-density lipoprotein (HDL) in the HUNT study with SKAT-O p-values < 2.5×10^−6^ from SAIGE-GENE. Conditional analysis was performed when the top hit in the locus(+/− 500kb of the start and end positions of the gene) is not included in the gene-based test. The p-value of conditional analysis is NA when the top hit is a rare missense or stop gain variant included in the gene-based test.

We also applied SAIGE-GENE to analyze 15,342 genes for 53 quantitative traits using 408,910 UK Biobank participants with White British ancestry2. Heritability estimates based on the full GRM are presented in **Supplementary Table 3A**. **Supplementary Table 4A** presents all genes with p-values reaching the exome-wide significant threshold (p ≤ 2.5×10^−6^). The same MAF cutoff ≤ 1%, for missense and stop-gain variants were applied. **Figure 2B** shows the QQ plot for automated read pulse rate as an exemplary quantitative phenotype in the UK Biobank. After conditioning on the most significant nearby variants, *MYH6*, *ARHGEF40* and *DBH* remain significant (**Table 1**). Gene *TBX5*, *MYH6*, *TTN*, and *ARHGEF40* are known genes for heart rates by previous GWAS studies^23–26^. To our knowledge, *KIF1C* and *DBH* have not been reported by association studies for heart rates, but both homozygous and heterozygous *DBH* mutant mice have decreased heart rates^27^. For the gene *DBH*, no single variant reaches the genome-wide significant threshold (the most significant variant is 9:136149399 (GRCh37) with MAF = 18.7% and p-value =3.46×10^−6^).

In the analysis of all 53 quantitative traits in the UK Biobank,199 gene-phenotype pairs were significant at exome-wide significant threshold (p ≤ 2.5×10^−6^). Among them fifteen genes for fourteen phenotypes were not significant by the single variant test, as the most significant single-variant association p-value in each of these loci (500kb up and down stream around each gene) did not reach the genome-wide threshold (p-value < 5×10^−8^)(**Supplementary Table 5**). For example, *TBX5*, which has been previously reported to be associated with heart rates^23^, was significant by SAIGE-GENE for the automated read pulse rate (p-value_SKAT-O_ = 2.87×10^−7^). However, the top variant in the locus was not genome-wide significant (p-value = 2.91×10^−7^). *ARID1B* has been previously reported to be associated with blood pressure in individuals with African ancestry^28^ and identified by SAIGE-GENE for automated read mean of diastolic blood pressure (p-value_SKAT-O_ = 1.08×10^−6^), while the most significant single variant association p-value was 9.01×10^−7^. In addition, SAIGE-GENE has identified several potentially novel gene-phenotype associations, including *DBH* for automated read pulse rate (p-value_SKAT-O_ =1.74×10^−6^),*C10orf35* for body fat percentage (p-value_SKAT-O_ = 3.64×10^−7^) a gene have been reported to be associated with type 2 diabetes^29^ and blood lipids by previous GWAS^30^. After conditioning on the most significant nearby variants, total 64 genes for 12 traits remained exome-wide significant (**Supplementary Table 6A**). Our results have successfully replicated several previous findings, such as the association between the rare coding variants of *ADAMTS3* and height^31^, *ZFAT* and height^31^, and *RRAS* and blood pressure^32^. These results have demonstrated the value of gene-based tests for identifying genetic factors for complex traits.

### Gene-based association analysis of binary traits in UK Biobank

We also applied SAiGE-GENE to ten binary phenotypes with various case-control ratios in the UK Biobank. The heritability estimates in a liability scale are presented in **Supplementary Table 3B**. Nine genes for six binary phenotypes reached the exome-wide significant threshold (p-value < 2.5×10^−6^) (**Supplementary Table 4B**), all of which have been identified by both SAIGE-GENE and single variant tests, including the gene *MYOC*, known for glaucoma^33^(**Figure 2C**). Six genes for six binary phenotypes remained exome-wide significant after conditioning on top variants (**Supplementary Table 6B**). Gene *GORASP1*, encoding Golgi Reassembly Stacking Protein 1 involved in the vesicle-mediated transport pathway, remained significant after conditioning on the top hit for diseases of hair and hair follicles.

### Simulation Studies

We investigated the empirical type I error rates and power of SAIGE-GENE through simulation. We followed the steps described in the Online Methods section to simulate genotypes and phenotypes for 10,000 samples in two settings. One has 500 families and 5,000 unrelated samples and the other one has 1,000 families, each with 10 family members based on the pedigree shown in **Supplementary Figure 11**.

### Type I error rates

The type I error rates of SAIGE-GENE, EmmaX-SKAT, and SMMAT have been evaluated based on gene-based association tests performed on 10^7^ simulated gene-phenotype combinations, each with 20 genetic variants with MAF ≤ 1% on average. A sparse GRM with a cutoff 0.2 for the coefficient of relatedness was used in SAIGE-GENE. Two different values of variance component parameter corresponding to the heritability *h*^*2*^ = 0.2 and 0.4 were considered for continuous traits, respectively (see **ONLINE METHODS**). The empirical type I error rates at the α = 0.05, 10^−4^ and 2.5×10^−6^ are shown in the **Supplementary Table 7**. Our simulation results suggest that SAIGE-GENE has relatively well controlled type I error rates, while the type I error rates are slightly inflated when heritability is relatively high (*h*^*2*^= 0. 4). Similar results have been observed on a larger sample size with 1,000 families and 10,000 unrelated samples (**Supplementary Materials 2.1** and **Supplementary Table 8**). Adjusting the test statistics using the genomic control (GC) inflation factor lambda has addressed the inflation (**Supplementary Materials 1.3.4**).

Further simulations have been conducted to evaluate type I error rates of SAIGE-GENE, EmmaX-SKAT, and SMMAT for skewed distributed phenotypes, which are common in real data (**Supplementary Figure 12A**). All three methods had inflated type I error rates for phenotypes having skewed distribution (**Supplementary Table 9**). With inverse normal transformation on phenotypes (**Supplementary Figure 12B**), the inflation has been dramatically reduced but slight inflation was still observed (**Supplementary Table 9**). A potential reason is that inverse normal transformation disrupts sample relatedness in raw phenotypes, leading to poor fitting for the null GLMM. We then conducted a three-step phenotype transformation procedure as described in **Supplementary Materials 2.2**, which maintains sample relatedness in raw phenotypes, and all three methods then have well controlled type I error rates (**Supplementary Table 10**). From simulation studies using real genotype data from the UK Biobank, we show that SAIGE-GENE well controlled type I error rates in the presence of subtle population structure or non-negligible cryptic relatedness between families (**Supplementary Table 11 and 12**). Details have been described in **Supplementary Materials 2.3** and **2.4**.

We have also evaluated the empirical type I error rates of SAIGE-GENE for binary traits with various case-control ratios. Similar with continuous traits, a sparse GRM with a cutoff 0.2 for the coefficient of relatedness was used. The variance component parameter *τ* =⍰l was assumed, corresponding to liability-scale heritability 0.23. As expected, when case-control ratios were balanced or moderately unbalanced (e.g. 1:1 and 1:9), type I error rates were well controlled even without the robust adjustment, while when the ratios were extremely unbalanced (e.g. 1:19 and 1:99), inflation was observed (**Supplementary Table 13A and Supplementary Figure 6**). With the robust adjustment combining SPA and ER, type I error rates were relatively well controlled in the presence of unbalanced case-control ratios (**Supplementary Table 13B and Supplementary Figure 6**). However, for phenotypes with case-control ratio=1:99, slight inflation was still observed, although the inflation has been dramatically alleviated compared to the unadjusted method. Then the genomic control adjustment can be used to further control the type I error rates (**Supplementary Table 13B**). We have also evaluated empirical type I error rates of SAIGE-GENE for binary traits under case-control sampling with case-control ratios 1:1 and 1:9 based on a disease prevalence 1% in the population (**Supplementary Materials 2.5**) and observed well-controlled type I error rates (**Supplementary Table 14**).

### Power

Next, we evaluated empirical power of SAIGE-GENE and EmmaX-SKAT for quantitative traits. Two different settings of proportions of causal variants were used: 10% and 40%. In each setting, among causal variants, 80% and 100% have negative effect sizes. The absolute effect sizes for causal variants are set to be |0.3log_10_(MAF)| and |log_10_(MAF)|, respectively, when the proportions of causal variants are 0.4 and 0.1. **Supplementary Table 15** shows that the power of both methods is nearly identical for all simulation settings for Burden, SKAT and SKAT-O tests.

We have also evaluated empirical power of SAIGE-GENE for binary traits using two different study designs: cohort study with various disease prevalence (0.01-0.5); and case-control sampling with different case-control ratios (1:1-1:19) based on a disease prevalence 1% in the population. In each setting, 40% variants are simulated as causal variants. Among them, 80% are risk-increasing variants and 20% are risk-decreasing. The absolute effect sizes of causal variants are set to be |0.55log_10_(MAF)| and |0.35log_10_(MAF)| for cohort study and case-control sampling, respectively. **Supplementary Table 16** shows the empirical power of SKAT-O in both simulation studies. SAIGE-GENE had similar empirical power as unadjusted SAIGE-GENE in balanced case-control ratios and higher power in unbalanced scenarios. The power is small when case: control ratio is 1:99 due to the limited number of cases (100 cases), which can be alleviated with larger sample size.

### Code and data availability

SAIGE-GENE is implemented as an open-source R package available at https://github.com/weizhouUMICHS/AIGE/master.

The summary statistics and QQ plots for 53 quantitative phenotypes and 10 binary phenotypes in UK Biobank by SAIGE-GENE are currently available for public download at https://www.leelabsg.org/resources.

## DISCUSSION

In summary, we have presented a method, SAIGE-GENE, to perform gene- or region-based association tests in large cohorts or biobanks in the presence of sample relatedness. Similar to SAIGE^9^, which was previously developed by our group for single-variant association tests, SAIGE-GENE uses generalized linear mixed models to account for sample relatedness, scalable computational approaches for large sample sizes, and the robust adjustment^14^ to account for unbalanced case-control ratios of binary traits. SAIGE-GENE uses several optimization strategies that are similar to those used in SAIGE to make fitting the null GLMM feasible for large sample sizes. For example, instead of storing the genetic relationship matrix (GRM) in the memory, SAiGE-GENE stores genotypes that are used for constructing the matrix in a binary vector and computes the elements of the matrix as needed. Preconditioned conjugate gradient algorithm is also used to solve linear systems instead of the Cholesky decomposition. However, some optimization approaches are specifically applied in the gene-based tests in regard of rare variants. As estimating the variances of score statistics for rare variants are more sensible to family structures, we use a sparse GRM to preserve close family structures rather than ignoring all sample relatedness. In addition, the variance ratios are estimated for different minor allele count (MAC) categories, especially for those extremely rare variants with MAC lower than or equal to 20.

For binary phenotypes, SAIGE-GENE applies the robust adjustment combining SPA and ER, thereby also relatively well controls the type I error rates for both balanced and unbalanced case-control phenotypes. However, slight inflation is still observed in extremely unbalanced phenotypes (≤:99). To address this possible issue, we suggest using the genomic control to further control type I error.

In numerical optimization, using good initial values can improve the model convergence. In the analysis of 24 quantitative traits in the UK Biobank with sample size (N) ≥ 100,000, we note that the models with the full GRM and the sparse GRM produced different variance component estimates, but they are relatively concordant (Pearson’s correlation R^2^ = 0.66, **Supplementary Figure 13**). This indicates that the parameter estimates from the sparse GRM can be used as initial values to facilitate the model fitting. We implemented this approach in SAIGE-GENE.

SAIGE-GENE has some limitations. First, similar to SAIGE and other mixed-model methods, the time for algorithm convergence to fit the generalize linear mixed models may vary among phenotypes and study samples given different heritability levels and sample relatedness. Second, similar to SAIGE^9^ and SMMAT^6^, SAIGE-GENE uses penalized quasi-likelihood (PQL)^34^ for binary traits to estimate the variance component in binary phenotypes which is known to be biased. However, as shown in simulation studies in SAIGE^9^ and SMMAT^6^, PQL-based approaches works well to adjust for sample relatedness.

Overall, we have shown that SAIGE-GENE can account for sample relatedness while maintaining test power through extensive simulation studies. By applying SAIGE-GENE to the HUNT study^9^ and the UK Biobank^2^ followed by conditioning on most significant variants in the testing loci, we have demonstrated that SAIGE-GENE can identify potentially novel association signals that are independent from the nearby association signals from the single-variant tests. Currently, our method is the only available mixed effect model approach for gene- or region-based rare variant tests for large sample data, while accounting for unbalanced case-control ratios for binary traits. By providing a scalable solution to the current largest and future even larger datasets, our method will contribute to identifying trait-susceptibility rare variants and genetic architecture of complex traits.

### URLs

SAIGE (version 0.35.8.8), https://github.com/weizhouUMICH/SAIGE/.

SMMAT (version 1.0.2), https://github.com/hanchenphd/GMMAT.

EmmaX-SKAT (SKAT version_l. 3.2.1), https://cran.r-project.org/web/packages/SKAT/index.html.

UK-Biobank analysis results (Gene-based summary statistics for 53 quantitative phenotypes in the UK Biobank by SAIGE-GENE), https://www.leelabsg.org/resources.

## Supporting information

Algorithm details, Supplementary Figures, Supplementary Tables

## ACKNOWLEDGMENTS

This research has been conducted using the UK Biobank Resource under application number 45227. SL and WB were supported by NIH R01HG008773. WZ was supported by NIH T32-Smoller 234736.

## AUTHOR CONTRIBUTIONS

W.Z., Z.Z., and S.L. designed experiments. W.Z., Z.Z., and S.L. performed experiments. W.Z. implemented the software with input from W.B. and J.L‥ J.B., L.G.F and S.A.G.T. constructed phenotypes for UK Biobank data. M.E.G. and K.H. provided data for the HUNT study. W.Z., Z.Z., C.W., S.L. and G.R. A. analyzed UK Biobank data. Helpful advice was provided by B.M.N and M.J.D‥W.Z., Z.Z., and S.L. wrote the manuscript with input from S.A.G.T. and M.E.G‥

## ONLINE METHODS

### Generalized linear mixed model

In a study with sample size *N*, we denote the phenotype of the *ith* individual using *y*_*i*_ for both continuous and binary traits. Let the 1 × *(p* + 1) vector *X*_*i*_ represent *p* covariates including the intercept, the *N* × *q* matrix *G*_*i*_ represent the allele counts (0,1 or 2) for *q* variants in the gene to test.

The generalized linear mixed model can be written as

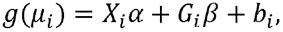

where *μ*_*i*_ is the mean of phenotype, *b*_*i*_ is the random effect, which is assumed to be distributed as *N*(O,τ *ψ*), where *ψ* is an *N* × *N* genetic relationship matrix (GRM) and τ is the additive genetic variance parameter. The link function *g* is the identity function for continuous traits with an error term *ε~*N*(O, ϕl)* and logistic function for binary traits. The parameter *α* is a *(p* + 1) × 1 coefficient vector of fixed effects and *β* is a *q* × 1 coefficient vector of the genetic effect.

### Estimate variance component and other model parameters (Step 1)

Same as in the original SAIGE^9^ and GMMAT^35^, to fit the null GLMM in SAIGE-GENE, penalized quasi-likelihood (PQL) method^34,36^ and the computational efficient average information restricted maximum likelihood (AI-REML) algorithm^35,37^ are used to iteratively estimate 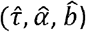 under the null hypothesis of *β* = 0. At iteration *k*,let 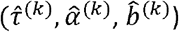 be estimated 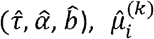 be the estimated mean of *y*_*i*_ and 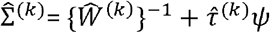 be an *N* × *N* matrix of the variance of working vector 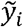, in which *ψ* is the *N* × *N* GRM. For continuous traits, 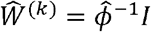 and 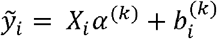. For binary traits, 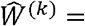 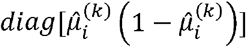 and 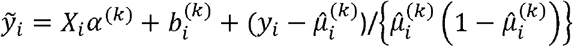. To obtain the log quasi-likelihood and average information at each iteration, SAIGE and SAIGE-GENE use the preconditioned conjugate gradient approach (PCG)^31,32^ to obtain the product of inverse of 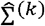 and any other vector by iteratively solving a linear system with 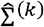. This approach is more computationally efficient than using Cholesky decomposition to obtain 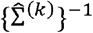. The numerical accuracy of PCG has been evaluated in the SAIGE paper^9^.

### Gene-based association tests (Step 2)

Test statistics of the Burden, SKAT and SKAT-O tests for a gene can be constructed based on score statistics from the marginal model for individual variants in the gene. Suppose there are *q* variants in the region or gene to test. The score statistic for variant *j*(j=l,‥, *q*) under *H*_*0*_: *β*_*j*_ = 0 is 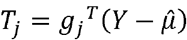 where *g*_*j*_ and *Y* are *N* × 1 genotype and phenotype vectors, respectively, and 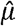 is the estimated mean of *Y* under the null hypothesis.

Let *u*_*j*_ denote a threshold indicator or weight for variant j and U = diag(*u*_*i*_,…, *u*_*q*_) be a diagonal matrix With *u*_*j*_ as the *j*th element. Similar to the original SKAT and SKAT-O papers^4,5^ to upweight rare variants, the default setting in SAIGE-GENE is *u*_*j*_ = *Beta*(*MAF*_*j*_,1, 25), which upweight rarer variants. The Burden test statistics can be written as 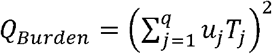. Suppose 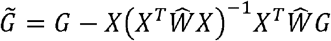 is the covariate adjusted genotype matrix, where *G* = (*g*_1_,…, *g_q_)* is the *N* × *q* genotype matrix of the *q* genetic variants, and 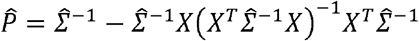 with 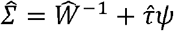. Under the null hypothesis of no genetic effects, *Q*_*Burden*_ followed 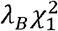, where 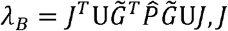 is a *q* × 1 vector with all elements being unity and 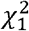 is a chi-squared distribution with 1 degree of freedom^3^. The SKAT test^4^ can be written as 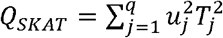, which follows a mixture of chi-square distribution 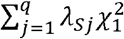, where *λ*_*Sj*_ are the eigenvalues of 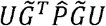. The SKAT-0 test^5^ uses a linear combination of the Burden and SKAT tests statistics *Q*_*sKATO*_ = (1 − *ρ)Q_sKAT_* + *ρQ_u8rden_*,0 ≤ *ρ* ≤ 1. To conduct the test, the minimum p-value from grid of *ρ* is calculated and the p-value of the minimum p-value is estimated through numerical integration. Following the suggestion in Lee *et al*^38^, we use a grid of eight values of *ρ* = (0, 0.1^2^, 0.2^2^, 0.3^2^, 0.4^2^, 0.5^2^, 0.5, 1) to find the minimum p-value.

### Approximate 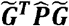

For each gene, given 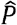, the calculation of 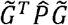 requires applying PCG for each variant in the gene, which can be computationally very expensive. Suppose *g̃* represents a covariate adjusted single variant genotype vector. To reduce computation cost, an approximation approach has been used in SAIGE, BOLT-LMM^17^ and GRAMMAR-GAMMAR^18^, in which the ratio between 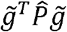 and 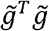 is estimated by a small subset of randomly selected genetic markers. The ratio has been shown to be approximately constant for all variants. Given the estimated ratio 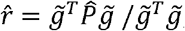,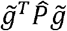 for all other variants can be obtained as 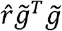. However, the variations of the estimated *r̂* for extremely rare variants are large and including some closely related samples in the denominator helps reduce the variation of *r̂* as shown in **Supplementary Figure 2**. Let *ψ*_*s*_ denote a sparse GRM that preserves close family structure and *ψ*_*f*_ denote a full GRM. We estimate the ratio 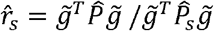, where 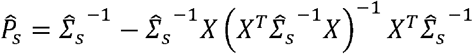 and 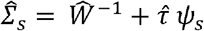.

In *ψ*_*s*_, elements below a user-specified relatedness coefficient cutoff, i.e. >3^rd^ degree relatedness, are zeroed out with only close family structures being preserved. To construct *ψ*_*s*_, a subset of randomly selected genetic markers, i.e. 2,000, is firstly used to quickly estimate which related samples pass the user-specified cutoff. Then the relatedness coefficients for those samples are further estimated using the full set of genetic markers, which equal to corresponding values in the *ψ*_*f*_. In the model fitting using 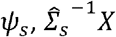 and 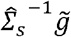 need to be calculated. For this we use a sparse-LU based solve method^39^ implemented in R. The constructed *ψ*_*s*_ is also used for approximating the variance of score statistics with *ψ*_*f*_. For a biobank or a data set, *ψ*_*s*_ only needs to be constructed once and can be re-used for any phenotypes in the same date set.

SAIGE-GENE estimates variance ratios for different MAC categories. By default, MAC categories are set to be MAC equals to 1, 2, 3, 4, 5, 6 to 10, 11 to 20, and is greater than 20. Once the MAC categorical variance ratios are estimated, for each genetic marker in tested genes or regions, *r̂*_*s*_ can be obtained according to its MAC. Let 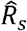 be a *q* × *q* diagonal matrix whose *j^th^* diagonal element is the ratio *r̂*_*s*_ for the jth marker in the gene (i.e.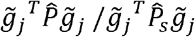). For the tested gene with *q* markers, 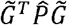 can be approximated as 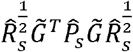 (See **Supplementary Materials** for more details).

### Robust adjustment for 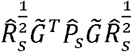 to account for unbalanced case-control ratios

To account for unbalanced case-control ratios of binary traits in region- or gene-based tests, we recently developed a robust adjustment for independent samples^14^. The approach first obtains well-calibrated p-values of single variant score statistics using SPA^10–12^ and ER^13^. SPA is a method to calculate p-values by inverting the cumulant generating function (CGF). Since CGF completely specifies the distribution, SPA can be far more accurate than using the normal distribution. However, since SPA is still an asymptotic based approach, it does not work well when variants are very rare (ex. MAC ≤10). For those variants, we use ER, which resamples the case-control status of only individuals carrying a minor allele and is extremely fast for very rare variants. To account for the fact that individuals can have different non-genetic risk of diseases (due to covariates), the resampling was done with the estimated disease riskμ. Next, variances of single variant score statistics are obtained by inverting those p-values, which are then used to calibrate the variances of region- or gene-based test statistics. We have extended the approach for related samples in SAIGE-GENE. For variants with MAC > 10, single-variant p-values are obtained by SAIGE, which basically applies SPA to GLMM. For variants with MAC ≤10, we use ER with GLMM estimated 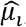, which includes the random effect to maintain the correlation structure among samples. After calculating p-values of *T*_*j*_ for j=l,…,q, the variance of *T*_*j*_ is calibrated by inverting the corresponding p-value. Then the calibrated variance is applied to 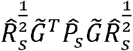 to compute robust p-value for the region-or gene-based test. The details can be found in **Supplementary Materials.**

### Conditional analysis

In SAIGE-GENE, we have implemented the conditional analysis to perform gene-based tests conditioning on a given markers using the summary statistics from the unconditional gene-based tests and the linkage disequilibrium *r*^2^ between testing and conditioning markers^19^. Let *G* be the genotypes for a gene to be tested for association, which contains *q* markers, and *G*_2_ be the genotypes for the conditioning markers, which contains *q*_2_ markers. Let *β* denote a *q* × 1 coefficient vector of the genetic effect for the gene to be tested and *β*_2_ be a *q*_2_ × 1 coefficient vector of the genetic effect for the conditioning markers. The genotype matrix with the non-genetic covariates projected out 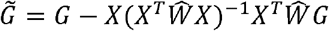 and 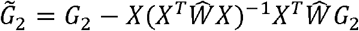. In the unconditioned association tests, the test statistics 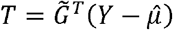 and 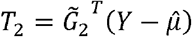. In conditional analysis, under the null hypothesis, 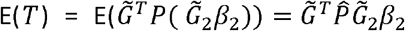 and 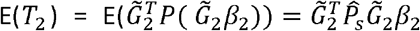. *T* and *T*_2_ jointly follow the multivariate normal with mean(E(*T*),E(*T*_2_)) and variance 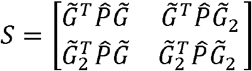

Thus under the null hypothesis of no association of T, i.e. H_0_: *β* = 0, the *T*|*T*_2_ follows the conditional normal distribution with 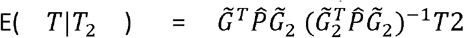 and 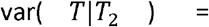 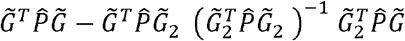, and p-values can be calculated from the conditional distribution.

### Data simulation

We carried out a series of simulations to evaluate and compare the performance of SAIGE-GENE, EmmaX-SKAT^5,7^ and SMMAT^6^. We used the sequence data from 10,000 European ancestry chromosomes over 1Mb regions that was generated using the calibrated coalescent model in the SKAT R package^5^. We randomly selected 10,000 regions with 3Kb from the sequence data, followed by the gene-dropping simulation^44^ using these sequences as founder haplotypes that were propagated through the pedigree of 10 family members shown in **Supplementary Figure 11**. Only variants with MAF ≤ 1% were used for simulation studies. Quantitative phenotypes were generated from the following linear mixed model *y*_*i*_ = *X*_1_ + *X*_2_ + *G*_*i*_*β* + *b*_*i*_ + *ɛ*_*i*_, where *G*_*i*_ is the genotype value, *β* is the genetic effect sizes, *b*_*i*_ is the random effect simulated from *N*(O, τ *ψ*), and *ɛ*_*i*_ is the error term simulated from *N*(O, (1 - τ)*I*).

Two covariates, X_1_ and X_2_, were simulated from Bernoulli(0.5) and N(0,1), respectively. Binary phenotypes were generated from the logistic mixed model *logit*(*π*_*io*_) = *α*_*0*_ + *b*_*i*_ + *X*_1_+ *X*_2_ + *G*_*i*_*β*, where *β* is the genetic log odds ratio, *b*_*i*_ is the random effect simulated from *N* (0, τ *ψ*) with τ = 1. The intercept *α*_0_ was determined by the disease prevalence (i.e. case-control ratios). Given τ = 1, the liability scale heritability is 0.23^45^.

To evaluate the type I error rates at exome-wide α=2.5×10^−6^, we first simulated 10,000 regions, and then simulated 1000 sets of quantitative phenotypes for each simulated region with different random seeds under the null hypothesis with *β* = 0. Gene-based association tests were performed using SAIGE-GENE, EmmaX-SKAT, and SMMAT therefore in total 10^7^ tests for each of Burden, SKAT, and SKAT-O tests were carried out. Two different settings for τ were evaluated: 0.2 and 0.4 and two different sample relatedness settings were used: one has 500 families and 5,000 independent samples and other one has 1,000 families, each with 10 family members. We also simulated 1,000 sets of binary phenotypes for case-control ratios 1:99, 1:19, 1:9, 1:4, and 1:1 for 500 families and 5,000 independent samples. Burden, SKAT, and SKAT-O tests were performed on the 10,000 genome regions using SAIGE-GENE, in total 10^7^ tests for each method for each case-control ratio.

For the power simulation, phenotypes were generated under the alternative hypothesis *β* ≠ 0. Two different settings for proportions of causal variants are used: 10% and 40%, corresponding to |*β*| = |*log10(MAF)*| and |*β*| = |*0.3log10(MAF)*|, respectively. In each setting, 80% and 100% had negative effect sizes. We simulated 1,000 datasets in each simulation, and power was evaluated attest-specific empirical α, which yields nominal α=2.5×10^−6^. The empirical α was estimated from the type I error simulations. Similarly, 1,000 sets of binary traits were generated for 10,000 samples (500 families and 5,000 independent samples) under the alternative hypothesis *β*≠ 0 using two different settings: cohort study with various disease prevalence (0.01, 0.05, 0.1, and 0.5); and case-control sampling with three different case-control ratios (1:19, 1:9, and 1:1) based on a disease prevalence 1% in the population (**Supplementary Materials 2.5**). 40% variants are simulated as causal variants, among which 80% are risk-increasing variants and 20% are risk-decreasing. The absolute effect sizes of causal variants are set to be |0.551og_10_(MAF)| and |0.351og_10_(MAF)| for cohort study and case-control sampling, respectively.

### HUNT and UK Biobank data analysis

We applied SAIGE-GENE to the high-density lipoprotein(HDL) levels in 69,500 Norwegian samples from a population-based Nord Trøndelag Health Study (HUNT) ^9^. About 70,000 HUNT participants were genotyped using Illumina HumanCoreExome v1.0 and 1.1 and imputed using Minimac3^40^ with a merged reference panel of HRC and whole genome sequencing data (WGS) for 2,201 HUNT samples. Variants with imputation r^2^ < 0.8 were excluded from further analysis. Total 13,416 genes with at least two rare (MAF ≤ 1%) missense and/or stop-gain variants with imputation r^2^ ≥ 0.8 were tested. Variants were annotated using Seattle Seq Annotations (http://snp.gs.washington.edu/SeattleSeqAnnotation138/). Age, Sex, genotyping batch, and first four PCs were included as covariates in the model. We used 249,749 pruned genotyped markers to estimate relatedness coefficients in the full GRM for Step 1 and used the relative coefficient cutoff ≥ 0.125 for the sparse GRM.

We have also analyzed 53 quantitative traits and 10 binary traits using SAIGE-GENE in the UK Biobank for 408,910 participants with White British ancestry^2^. Markers that were imputed by the Haplotype Reference Consortium (HRC)^20^ panel with imputation info score ≥ 0.8 were used in the analysis. Total 15,342 genes with at least two rare (MAF ≤ 1%) missense and stop-gain variants that were directly genotyped or successfully imputed from HRC (imputation score 0.8) were tested. Sex, age when attended assessment center, and first four PCs that were estimated using all samples with White British ancestry were adjusted in all tests. We used 340,447 pruned markers, which were pruned from the directly genotyped markers using the following parameters, were used to construct GRM: window size of 500 base pairs(bp), step-size of 50 bp, and pairwise r^2^ < 0.2.We used the relative coefficient cutoff ≥ 0.125 for the sparse GRM.

### Genome build

All genomic coordinates are given in NCBI Build 37/UCSC hg19.

### Reporting Summary

Further information on study design is available in the Nature Research Reporting Summary linked to this article.

