## Supplementary material for "Scalable generalized linear mixed model for region-based association tests in large biobanks and cohorts": Algorithm details, Supplementary Figures, Supplementary Tables

### Supplementary Materials

### 1. Algorithm details

#### 1.1 Step 0. Constructing the sparse GRM

In the sparse GRM, denoted by $\psi_s$, GRM elements below a user-specified relative coefficient cutoff are zeroed out with close family structures preserved. To improve the test accuracy for rare variants, SAIGE-GENE approximates the variance of score statistics calculated with the full GRM $\psi_f$ using the variance calculated with the sparse GRM $\psi_s$ and the ratios of these two variance estimates estimated using a subset of genetic markers.

To construct the sparse GRM $\psi_s$, a small subset of randomly select markers were used to identify related sample pairs whose relative coefficient pass the use-specified cutoff, which is to find out the indices of non-zero elements in $\psi_s$. Next, the values of the nonzero elements in $\psi_s$ are then estimated using the full set of genetic markers that are used in Step 1 for $\psi_f$. This step is only needed for once for each data set or biobank and parallel computation is allowed. Once the sparse GRM is constructed for a data set, it can be re-used in SAIGE-GENE for all phenotypes.

#### 1.2 Step 1. Fitting the null generalized linear mixed model

The same model fitting framework and computation approaches used in the original SAIGE[1] are used in SAIGE-GENE to fit the null GLMM for large sample sizes. These include estimating model parameters using the AI-REML approach[2,3], solving linear systems by the preconditioned conjugate gradient method[4], using Hutchinson's randomized trace estimator[5,6] to obtain traces of matrices, and allowing for parallel computation for the vector multiplication. For AI-REML, we particularly used the approach in GMMAT[2,7] to calculate the average information without performing the *n* by *n* matrix inversion by using PCG. For details of the likelihood, parameter estimates and information matrices, please refer to the Supplementary Note in the SAIGE paper[1]. In addition, SAIGE-GENE can estimate variance component parameters, thus heritability, by fitting a null GLMM using the sparse GRM. The estimated variance component parameters can then be used as initial values for the model fitting with the full GRM, which can be a better approach than using a randomly chosen initial value. By plotting the heritability estimates using the sparse GRM versus using the full GRM for 24 quantitative traits with sample size larger than or equal to 10,000 from the UK Biobank (**Supplementary Figure 13**), we have shown that variance component estimates from the full and sparse GRMs are relatively concordant (Pearson's correlation $R^2$ = 0.66). For real-data analysis, robust performance of convergence has also been observed for some phenotypes, such as waist hip ratio (N = 408,144) in the UK Biobank. Using initial values 0.5 for heritability, step 1 did not even converge after 6,300 CPU hours, while using initial values estimated with the sparse GRM, it took 1836 CPU hours to finish the step 1.

#### 1.3 Step 2. Gene-based association tests

Test statistics of the Burden, SKAT and SKAT-O tests for a gene can be constructed based on the score statistics from the marginal model for individual variants in the gene. Suppose there are $q$ variants in the region or gene to test. The score test statistics for variant $j$ (j=1,.., $q$) under $H_o$: $\beta_j = 0$ is $T_j = g_j^T(Y - \hat{\mu})$

where $g_j$ and $Y$ are $N \times 1$ genotype and phenotype vectors, respectively, and $\hat{\mu}$ is the estimated mean of $Y$ under the null hypothesis.

Let $u_j$ denote a threshold indicator or weight for variant $j$ and U be a diagonal matrix with $u_j$ as the $j$th element. The Burden test statistics can be written as $Q_{Burden} = \left(\sum_{j=1}^{q} u_j T_j\right)^2$. Suppose $\tilde{G} = G - X(X^T \widehat{W} X)^{-1} X^T \widehat{W} G$, where $G = (g_1, \ldots, g_q)$ is the $N \times q$ genotype matrix of the $q$ genetic variants, and $\hat{P} = \hat{\Sigma}^{-1} - \hat{\Sigma}^{-1} X \left(X^T \hat{\Sigma}^{-1} X\right)^{-1} X^T \hat{\Sigma}^{-1}$ with $\hat{\Sigma} = \widehat{W}^{-1} + \hat{\tau}\psi$. Under the null hypothesis of no genetic effects, $Q_{Burden}$ followed $\lambda_B \chi_1^2$, where $\lambda_B = J^T \mathrm{U} \tilde{G}^T \hat{P} \tilde{G} \mathrm{U} J$ and $J$ is a $q \times 1$ vector with all elements being unity and $\chi_1^2$is a chi-squared distribution with 1 degree of freedom[8]. The SKAT test[9] can be written as $Q_{SKAT} = \sum_{j=1}^{q} u_j^2 T_j^2$, which follows a mixture of chi-square distribution $\sum_{j=1}^{q} \lambda_{Sj} \chi_1^2$, where $\lambda_{Sj}$ are the eigenvalues of $U \tilde{G}^T \hat{P} \tilde{G} U$. The SKAT-O test developed by Lee et al in 2012 [10] uses a linear combination of the Burden and SKAT tests statistics $Q_{SKATO} = (1-\rho) Q_{SKAT} + \rho Q_{Burden}, 0 \leq \rho \leq 1$. To conduct the test, the minimum p-value from grid of $\rho$ is calculated and the p-value of the minimum p-value is estimated through numerical integration. Following the suggestion in Lee *et al*[11], we use a grid of eight values of $\rho = (0, 0.1^2, 0.2^2, 0.3^2, 0.4^2, 0.5^2, 0.5, 1)$ to find the minimum p-value.

##### 1.3.1 Estimating $\tilde{G}^T \hat{P} \tilde{G}$

For each gene, given $\hat{P}$, calculation of $\tilde{G}^T \hat{P} \tilde{G}$ can be computationally expensive Suppose $\tilde{g} = g - X\left(X^T \widehat{W} X\right)^{-1} X^T \widehat{W} g$, which represents a covariate adjusted single variant genotype $N \times 1$ vector vector. To reduce computation cost, an approximation approach has been used in SAIGE[1], BOLT-LMM[12] and GRAMMAR-GAMMAR[13], in which the ratio between $\tilde{g}^T \hat{P} \tilde{g}$ and $\tilde{g}^T \tilde{g}$ is estimated by a small subset of randomly selected genetic markers that has been shown to be approximately constant for all variants[1]. Given the ratio $\hat{r} = \tilde{g}^T \hat{P} \tilde{g} / \tilde{g}^T \tilde{g}$, $\tilde{g}^T \hat{P} \tilde{g}$ for all other variants can be easily obtained as $\hat{r} \tilde{g}^T \tilde{g}$. However, the variations of estimated $\hat{r}$ for extremely rare variants are large and including some closely related samples in the denominator helps reduce the variation of $\hat{r}$ as shown in **Supplementary Figure 2**. It can also be observed from the plots in **Supplementary Figure 2** that the variance ratio for those extremely rare variants could be quite different from the ratio for more frequent variants, so SAIGE-GENE estimates variance ratios for different MAC categories. By default, MAC categories are set to be MAC equals to 1, 2, 3, 4, 5, 6 to 10, 11 to 20, and is greater than 20. For each MAC category, a ratio $\hat{r}_s$ is estimated as the average of the ratios computed from 30 randomly selected markers, among which every marker has a ratio $\tilde{g}^T \hat{P} \tilde{g} / \tilde{g}^T \hat{P}_s \tilde{g}$, where $\hat{P}_s = \hat{\Sigma}_s^{-1} - \hat{\Sigma}_s^{-1} X \left(X^T \hat{\Sigma}_s^{-1} X\right)^{-1} X^T \hat{\Sigma}_s^{-1}$ and $\hat{\Sigma}_s = \widehat{W}^{-1} + \tau \psi_s$. $\psi_S$ is a sparse GRM that preserves closely related samples. The coefficient of variance (CV) of $\hat{r}$ is used to evaluate the numerical stability of the $\hat{r}$ estimation. As in SAIGE, the default value of CV threshold is 0.001. If CV of $\hat{r}$ is larger than the threshold, SAIGE-GENE will increase the number of markers by 10 to estimate $\hat{r}$ until the estimation is stable with CV below or equal to the threshold. Once the variance ratios have been estimated for different MAC categories. For each genetic marker in genes or regions that are to be tested in Step 2, a $\hat{r}_s$ can be obtained according to its MAC. Let $\hat{R}_s$ be a $q \times 1$ vector whose jth element is the ratio $\hat{r}_s$ for the jth marker in the tested gene. For the tested gene with $q$ markers, $\tilde{G}^T \hat{P} \tilde{G}$ can be approximated as $\hat{R}_S^{\frac{1}{2}} \tilde{G}^T \hat{P}_S \tilde{G} \hat{R}_S^{\frac{1}{2}}$.

Products of $\hat{\Sigma}_s^{-1}$and other vectors or matrices are obtained using the sparse LU decomposition through the solve function in R[14]. Note that the $N \times N$ matrix $\hat{P}_s$ is not a sparse matrix because $\hat{\Sigma}_s^{-1}$is not sparse and $\tilde{G}$ is also a dense matrix, which is converted from the sparse matrix $G$, as $\tilde{G} = G - X\left(X^T\widehat{W}X\right)^{-1}X^T\widehat{W}G$. It can be shown that

$$\tilde{G}^T\hat{P}_s\tilde{G} = \left(G^T - G^T\widehat{W}X\left(X^T\widehat{W}X\right)^{-1}X^T\right)\left(\hat{\Sigma}_s^{-1} - \hat{\Sigma}_s^{-1}X\left(X^T\hat{\Sigma}_s^{-1}X\right)^{-1}X^T\hat{\Sigma}_s^{-1}\right)\left(G - X\left(X^T\widehat{W}X\right)^{-1}X^T\widehat{W}G\right) = G^T\hat{P}_sG$$

Computation of $G^T\hat{P}_sG$ is more computational efficient than that of $\tilde{G}^T\hat{P}_s\tilde{G}$ as $G$ can be stored as a sparse matrix.

##### 1.3.2 Conditional analysis

To test whether the association signals from a tested gene or region are independent from a given marker or multiple markers, the conditional analysis based on summary statistics from unconditional association tests with the linkage disequilibrium $r^2$ among testing and conditioning markers[15] have been implemented in SAIGE-GENE.

Let $G$ be the genotypes for a gene to be tested for association, which contains $q$ markers, and $G_2$ be the genotypes for the conditioning markers, which contains $q_2$ markers. Let $\beta$ denote a $q \times 1$ coefficient vector of the genetic effect for the gene to be tested and $\beta_2$ be a $q_2 \times 1$ coefficient vector of the genetic effect for the conditioning markers. The genotype matrix with the non-genetic covariates projected out $\tilde{G} = G - X\left(X^T\widehat{W}X\right)^{-1}X^T\widehat{W}G$ and $\tilde{G}_2 = G_2 - X\left(X^T\widehat{W}X\right)^{-1}X^T\widehat{W}G_2$. In the unconditioned association tests, the test statistics $T = \tilde{G}^T(Y - \hat{\mu})$ and $T_2 = \tilde{G}_2^{\,T}(Y - \hat{\mu})$. In conditional analysis, under the null hypothesis, E($T$) = E($\tilde{G}^TP(\tilde{G}_2\beta_2)) = \tilde{G}^T\hat{P}\tilde{G}_2\beta_2$ and E($T_2$) = E($\tilde{G}_2^TP(\tilde{G}_2\beta_2)) = \tilde{G}_2^T\hat{P}_s\tilde{G}_2\beta_2$. $T$ and $T_2$ jointly follow the multivariate normal with mean (E($T$), E($T_2$)) and variance $S = \begin{bmatrix}\tilde{G}^T\hat{P}\tilde{G} & \tilde{G}^T\hat{P}\tilde{G}_2 \\ \tilde{G}_2^T\hat{P}\tilde{G} & \tilde{G}_2^T\hat{P}\tilde{G}_2\end{bmatrix}$.

Thus under the null hypothesis of $\beta$ =0, the $T|T_2$ follows the conditional distribution E( $T|T_2$ ) = $\tilde{G}^T\hat{P}\tilde{G}_2\,(\tilde{G}_2^T\hat{P}\tilde{G}_2)^{-1}T_2$ and var( $T|T_2$ ) = $\tilde{G}^T\hat{P}\tilde{G} - \tilde{G}^T\hat{P}\tilde{G}_2\left(\tilde{G}_2^T\hat{P}\tilde{G}_2\right)^{-1}\tilde{G}_2^T\hat{P}\tilde{G}$ = $G^T\hat{P}G - G^T\hat{P}G_2(G_2^T\hat{P}G_2)^{-1}G_2^T\hat{P}G$. The test statistic of the conditional analysis can be written as (T - E($T|T_2$))$^2$/ var($T|T_2$), which follows the $\chi^2$ distribtuion with one degree of freedom. Similar to the unconditional analysis, we approximate these terms by $\tilde{G}^T\hat{P}_s\tilde{G}, \tilde{G}^T\hat{P}_s\tilde{G}_2, \tilde{G}_2^T\hat{P}_s\tilde{G}_2$, and $\tilde{G}_2^T\hat{P}_s\tilde{G}$ and the corresponding variance ratio matrices.

##### 1.3.3 Robust adjustment to account for unbalanced case-control ratios in binary traits

To account for unbalanced case-control ratios in binary traits, we use a recently developed robust adjustment approach with a simple modification[16]. Note that the robust adjustment was developed for independent samples. It uses saddlepoint approximation (SPA)[17-19] and efficient resampling (ER)[20] to obtain accurate single variant association P-value for the $jth$ variant and then calibrates the variance of score statistics $T_j$. SPA is a statistical method to calculate the distribution function using the cumulant generating function (CGF). Suppose $K_j(t)$ is the CGF of the score statistic $T_j$, which can be derived based on the fact that $Y_i \sim \text{Bernoulli}(\mu_i)$ under the null. For independent samples, the estimation of $K_j(t)$ is

$$\widehat{K_j}(t; \hat{\mu}, c) = \sum_{i=1}^{N}\log\left(1 - \widehat{\mu_i} + \widehat{\mu_i}e^{t\widetilde{G_i}}\right) - t\sum_{i=1}^{N}\widetilde{G_i}\widehat{\mu_i},$$

where $\widehat{\mu_i}$ is the estimation of $\mu_i$ from the null model, and $\widetilde{G_i}$ is the covariate adjusted genotype vector. To account for sample relatedness, we use the original SAIGE, which adapts the SPA to GLMM. Under the GLMM, we use the following CGF

$$\widehat{K_j}(\mathrm{t}; \hat{\mu}, \mathrm{c}) = \sum_{i=1}^{N} \log\left(1 - \hat{\mu}_i + \hat{\mu}_i e^{ct\widetilde{G_i}}\right) - ct\sum_{i=1}^{N} \widetilde{G_i}\widehat{\mu_i},$$

where $c = \mathrm{Var}^*(T_j)^{-1/2}$ and $\mathrm{Var}^*(T_j) = \widetilde{G_j}^T W \widetilde{G_j}$ is a variance estimator without accounting the fact that the random effect $b$ is estimated from data. Then, the distribution function of the score statistic $T_j$ can be approximated by

$$\Pr(T_j < q) = \tilde{F}(q) = \Phi\left\{w + \frac{1}{w} log\left(\frac{v}{w}\right)\right\},$$

where $w = sgn(\hat{t})\sqrt{2\left(\hat{t}s - K_j(\hat{t})\right)}$, $v = \hat{t}\sqrt{K_j''(\hat{t})}$, $\hat{t}$ is the solution to the equation $K_j'(\hat{t}) = q$, and $\Phi$ is the distribution function of the standard normal distribution. For details, please refer to the SAIGE paper[1].

Since SPA is an asymptotic based approach, it can provide incorrect p-values when MAC is very low (ex. MAC < 10). To address this issue, we use ER[20] for variants with MAC $\leq$ 10. ER is a resampling method that resamples the case–control status of individuals with a minor allele at a given variant given the disease risk $\mu_i$. ER was developed under the assumption that samples are independent. Here we apply ER with the disease risk $\widehat{\mu_i}$ estimated by GLMM, using the fact that given random effects $b_i$, samples are independent. The dependency among samples are incorporated through the random effect estimates.

The remaining part is nearly identical as the robust method in independent samples[16]. Let $\hat{V} = diag(\hat{v}_1, \dots, \hat{v}_q)^T$ is a $q$x$q$ diagonal matrix of the diagonal element of $\hat{R}_S^{\frac{1}{2}}\tilde{G}^T \hat{P}_S \tilde{G} \hat{R}_S^{\frac{1}{2}}$. We note that $j$-th diagonal element of $\hat{V}$ is the estimated variance of $T_j$. For each variant $j$, when the score statistic $T_j$ lies outside of two standard deviations of the mean (i.e. zero), we apply SPA (when MAC > 10) or ER (when MAC $\leq$ 10) to calculate the p-value $\tilde{p}_j$, and calculate $\tilde{v}_j = T_j^2/\chi^2_{quantile}(1 - \tilde{p}_j)$, where $\chi^2_{quantile}$ is the quantile function of the chi-square distribution with one degree of freedom. If $T_j$ lies within the two standard deviations of the mean, we use $\tilde{v}_j = \hat{v}_j$. Suppose $\tilde{R} = diag(\tilde{v}_1/\hat{v}_1, \tilde{v}_2/\hat{v}_2, \dots, \tilde{v}_q/\hat{v}_q)$ is a $q$x$q$ diagonal matrix. Now we use $\tilde{R}^{\frac{1}{2}} \hat{R}_S^{\frac{1}{2}}\tilde{G}^T \hat{P}_S \tilde{G} \hat{R}_S^{\frac{1}{2}} \tilde{R}^{\frac{1}{2}}$ as the estimate of $\tilde{G}^T \hat{P} \tilde{G}$ to calculate p-values and carry out conditional analysis.

#### 1.3.4. Genomic Control (GC) for the further adjustment of p-values

To further control for type I error rates, SAIGE-GENE allows for using the genome control inflation factor. Let $\lambda_{GC}$ be the genome control inflation factor from the gene-based test, which is obtained by converting p-values of gene-based test to $\chi^2_1$ statistics. To better capture the inflation in tail areas we obtain $\lambda_{GC}$ at p-value=0.05. And then we divide the $\lambda_{GC}$ in the $\chi^2_1$ statistics and then obtain the p-values using $\chi^2_1$ distribution. As shown in **Supplementary Table 7** and **Supplementary Table 13**, this simple approach has successfully attenuated the type I error inflation.

#### 1.4. Novel features in SAIGE-GENE compared to SMMAT

Same as SMMAT, SAIGE-GENE uses the logistic mixed model to conduct region- or gene-based association tests (Burden, SKAT and SKAT-O). Compared to SMMAT, SAIGE-GENE mainly has two improvements, which make it the only method so far that is feasible for large samples sizes, while accounting for case-control imbalance for binary phenotypes. It utilizes optimization strategies as used in the original SAIGE for the scalability, including storing raw genotypes in a binary vector and elements of GRM are calculated when needed rather than being stored to reduce the memory usage, replacing the cholesky decomposition by the preconditioning conjugate gradient (PCG) to solve linear systems without calculating and inverting the $N \times N$ GRM, and approximating the variance of score statistics with the full GRM using the variance with a sparse GRM and the ratio of the two variances. To account for unbalanced case-control ratios for binary phenotypes, SAIGE-GENE uses a robust adjustment approach combing SPA[17-19] and ER[20] as described in 1.3.3.

### 2. Additional simulation and real-data analysis results

#### 2.1 Simulation studies with a larger sample size

Given that SAIGE-GENE has relatively well controlled type I error rates based on simulation studies (**Supplementary Table 7**), it is noted that type I error rates are slightly higher for SAIGE-GENE than the other two methods. One potential reason for this could be that SAIGE-GENE uses several approximation approaches, as described in **1.4**. to achieve feasibility for large data sets and a tradeoff exists between accuracy and computational efficiency for these approaches. To evaluate type I error rates with larger sample sizes, we conducted a simulation study containing 1,000 families and 10,000 independent samples, which doubles the sample sizes as in the original simulation with 500 families and 5,000 independent samples. With the heritability $\mathrm{h}^2 = 0.2$, the empricial type I error rates for the Burden test, SKAT, and SKAT-O at three different α were estimated based on $10^7$ tests, for which 1,000 randomly simulated phenotype sets and each was tested on 10,000 variant sets (**Supplementary Table 8**). The type I error rates with the larger sample size (**Supplementary Table 8**) are similar to those with a smaller sample size with 500 families and 5,000 independent samples (**Supplementary Table 7**).

#### 2.2 Simulation studies with skewed distributed phenotypes

In simulation studies, the phenotype $y_i$ was generated to follow a normal distribution, but it can be skewed in real data. Hence, we have conducted additional simulation studies to evaluate the type I error rates of SAIGE-GENE in presence of skewed phenotypic distributions and compared to the other two methods. As described in the Data Simulation subsection of ONLINE METHODS**,** phenotypes were simulated from the following linear mixed model $y_i = X_1 + X_2 + G_i\beta + b_i + \varepsilon_i$ , where $G_i$ is the genotype value, $\beta$ is the genetic effect sizes. Two covariates, $X_1$ and $X_2$ , were simulated from Bernoulli(0.5) and N(0,1), respectively. $b_i$ is the random effect simulated from $N(0, \tau\psi)$ and $\varepsilon_i$ is the error term simulated from chi-square distribution with degree of freedom 1 and thereby the distribution of $y$ is skewed (**Supplementary Figure 12A**). 500 families and 5,000 independent samples were simulated and the empirical type I error rates for Burden test, SKAT, and SKAT-O at the three different α were estimated based on $10^7$ tests, for which 1,000 randomly simulated phenotype sets and each was tested on 10,000 variant sets (**Supplementary Table 9**). All three methods have inflated type I error rates for phenotypes having skewed distribution, especially in SKAT and SKAT-O tests. After the phenotypes were inverse normal transformed (**Supplementary Figure 12B**), type I error rate inflation has been substantially reduced (**Supplementary Table 9**).

Note that there is still slight inflation in all three methods after the inverse normal transformation on phenotypes, which can be because the inverse normal transformation may disrupt sample relatedness in the original phenotypes and thus impact the null model fitting. We then conducted a three-step phenotype transformation procedure, which is presumably able to avoid the issue above. Firstly, we fitted the null mixed model using raw skewed distributed phenotypes. Next, we conducted the inverse normal transformation on the residuals from step 1. Finally, transformed residuals were then used to fit another null mixed model, followed by gene- or region-based association tests. As expected, using this three-step phenotype transformation procedure, the type I error rates are well controlled in both SAIGE-GENE and SMMAT for all Burden, SKAT, and SKAT-O tests (**Supplementary Table 10**).

#### 2.3 Simulation studies in the presence of population structure

To evaluate whether SAIGE-GENE can control type I error rates in the presence of subtle population stratification, we randomly selected 5,000 white British-UK samples and 5,000 non-UK European samples from the UK-Biobank after removing up to 3rd relatives. The phenotypes were simulated based on real genotypes of randomly selected $L$= 30,000 LD-pruned (r2 < 0.2) markers with MAF $\geq$ 1%. In particular, phenotypes were simulated following the model $y_i = X_{i1} + X_{i2} + \sum_{j=1}^{L} \hat{G}_{ij}\, \beta + \varepsilon_i$, where $\hat{G}_{ij}$ is the standardized genotype value for the $jth$ marker of $ith$ individual, $\beta$ is the genetic effect size following $N(0, \tau/L)$, where $\tau = 0.2$, and $\varepsilon_i$ is the error term simulated from $N(0, (1-\tau)I)$. Two covariates, $X_{i1}$ and $X_{i2}$, were simulated from Bernoulli(0.5) and N(0,1). We conducted ~10,000 gene-based tests for each simulated phenotype set, replicated the simulation for 1,000 times. Note that we include the first 4 principal components, which were estimated for all European participants in the UK Biobank, as well as $X_1$ and $X_2$ as covariates in the linear mixed model. We evaluated the empirical type I error rates at the α = 0.05, $10^{-4}$ and $2.5 \times 10^{-6}$ as shown in **Supplementary Table 11**, which suggest that SAIGE can produce well calibrated p-values in the presence of subtle population stratification.

#### 2.4 Simulation studies in the presence of non-negligible cryptic relatedness between families

To evaluate whether SAIGE-GENE can control type I error rates in the presence of non-negligible cryptic relatedness between families, we have randomly selected 10,000 samples with white British ancestry from UK Biobank. In particular, 5,000 samples were selected among those who are up to $3^{rd}$ degree relatives and 5,000 sample were selected from the rest of the unrelated pool. Phenotypes were simulated using the same approach in Section 2.3. We conducted ~10,000 gene-based tests for each simulated phenotype set, replicated the simulation for 1,000 times. For each phenotype set, a null linear mixed model was fitted in Step 1 with covariates including the first 4 principal components, which were estimated for all White-British participants in the UK Biobank, and $X_1$ and $X_2$. We evaluated the empirical type I error rates at the α = 0.05, $10^{-4}$ and $2.5 \times 10^{-6}$ as shown in **Supplementary Table 12**. These results have indicated that SAIGE can produce well calibrated type I error rates in the presence of non-negligible cryptic relatedness between families.

#### 2.5 Simulation studies under case-control sampling

We have conducted additional simulation studies to evaluate the performance of SAIGE-GENE under case-control sampling. We simulated genotypes and phenotypes with prevalence 1% for 250,000 independent samples and 250,000 families, each with 10 family members (**Supplementary Figure 11**) as an underlying large cohort. We then randomly selected 5,000 controls, together with the 5,000 cases, for a phenotype with case-control ratio 1:1. In addition, for a phenotype with case-control ratio 1:9, we randomly selected 500 cases and 9,500 controls from the large cohort. Empirical type I error rates of SAIGE-GENE have been evaluated based on the 10 million tests (**Supplementary Table 14**), indicating that the type I error rates of SAIGE-GENE were well controlled under the case-control sampling. We have also evaluated the empirical power of SAIGE-GENE under case-control sampling through simulation studies as described in the RESULTS section of main texts. Results are presented in **Supplementary Table 16,** showing that under case-control sampling, the empirical power of SAIGE-GENE with and without robust adjustment is similar when the case-control ratio is relatively balanced (1:1). As expected, SAIGE-GENE with robust adjustment has higher power in unbalanced scenarios ($\leq 1{:}\,9$) than SAIGE-GENE without robust adjustment. Similar patterns have been observed for empirical power of SAIGE-GENE in simulations of a cohort study (**Supplementary Table 16**). Note that the power in cohort study and case-control sampling is not directly comparable due to the difference effect sizes.

#### 2.6 Exome-wide gene-based tests for automated read pulse rate in UK Biobank with different relatedness cutoffs in the sparse GRM

As a sensitivity analysis, we used another sample relatedness cutoff 0.2 for the sparse GRM to analyze automated read pulse rates in UK Biobank, which means all elements in the full GRM below 0.2 are zero'd in the sparse GRM. Scatter plots comparing p-values of the 15,342 genes with two different sample relatedness cutoff (0.125 and 0.2) are presented in **Supplementary Figure 5**, showing highly concordant association p-values for all three gene-based tests: Burden, SKAT, and SKAT-O tests.

### 3. Supplementary figures

**Supplementary Figure 1.** Workflow of SAIGE-GENE.

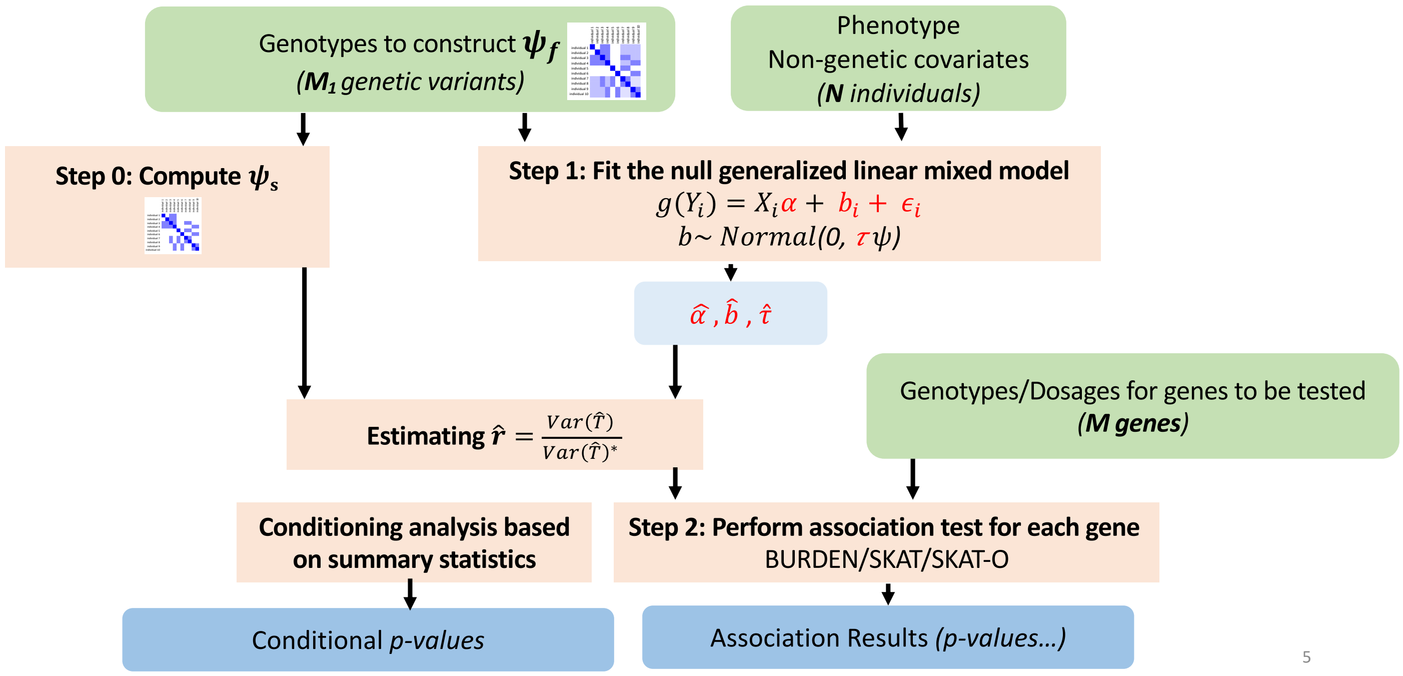

**Supplementary Figure 2.** Plots of the variance ratio of the score statistics by MAC for rare variants with and without the full GRM for sample relatedness (left) and with the full GRM and a sparse GRM for closely related samples(right). A. 500 families and 5,000 independent individuals were simulated with $h^2$= 0.2 based on the pedigree structure shown in **Supplementary Figure 11.** The sparse GRM was constructed using a coefficient of relatedness cutoff 0.2. B. 20,000 samples with White British ancestry were randomly selected from the UK Biobank and the null model was fitted for the automated read pulse rate. The sparse GRM was constructed using a coefficient of relatedness cutoff 0.125. C. 20,000 samples were randomly selected form the HUNT study and the null model was fitted for HDL. The sparse GRM was constructed using a coefficient of relatedness cutoff 0.125.

A. Simulation: 500 families and 5,000 independent individuals

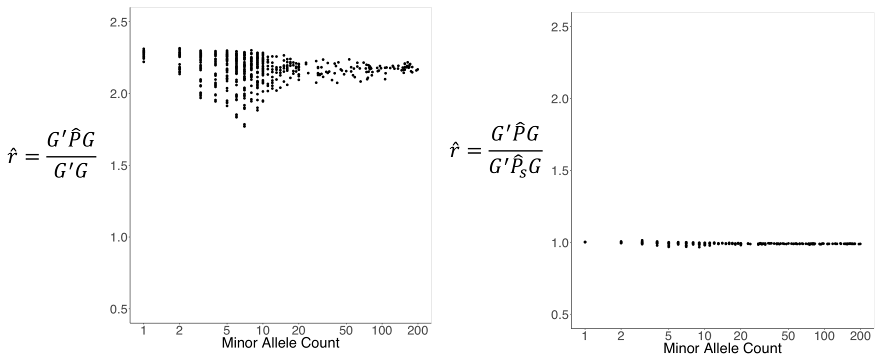


B. UK Biobank: Pulse rate automated read (mean), randomly selected 20,000 individuals

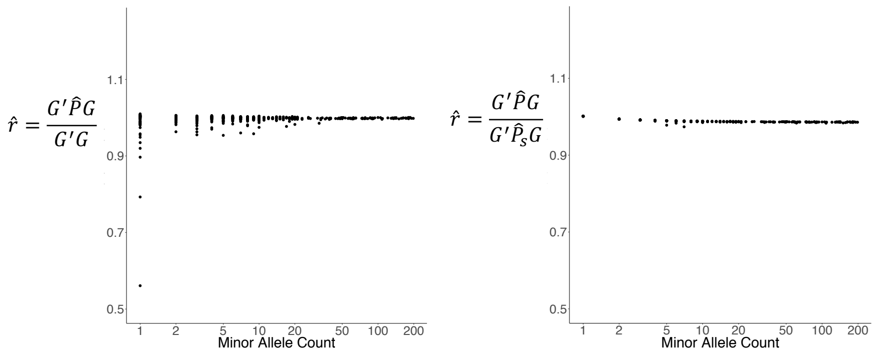


C. HUNT: HDL, randomly selected 20,000 individuals

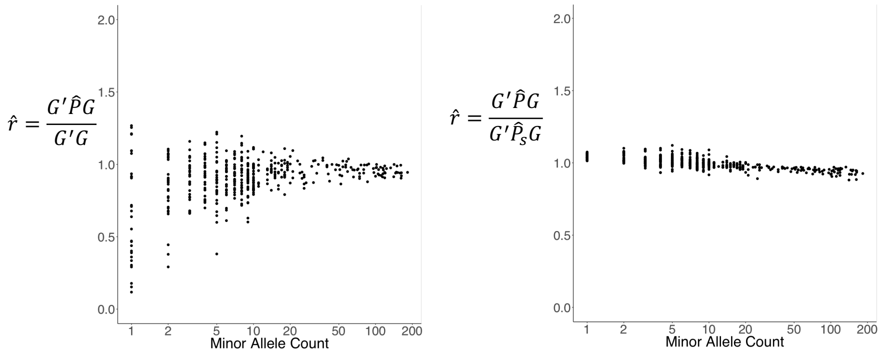

**Supplementary Figure 3.** Scatter plots of association p-values from SAIGE-GENE versus SMMAT[21] and EmmaX-SKAT for the Burden, SKAT, and SKAT-O tests based on simulation data on the -log10 scale. 1,000,000 genes were tested with 1000 families, each having 10 members, as shown in the **Supplementary Figure 11.** The Pearson's correlation coefficients $r^2 > 0.99$ for -log10(P-values) between SAIGE and SMMAT and between SAIGE and EmmaX-SKAT. A. $h^2$= 0.2, B. $h^2$= 0.4

A. $h^2$= 0.2

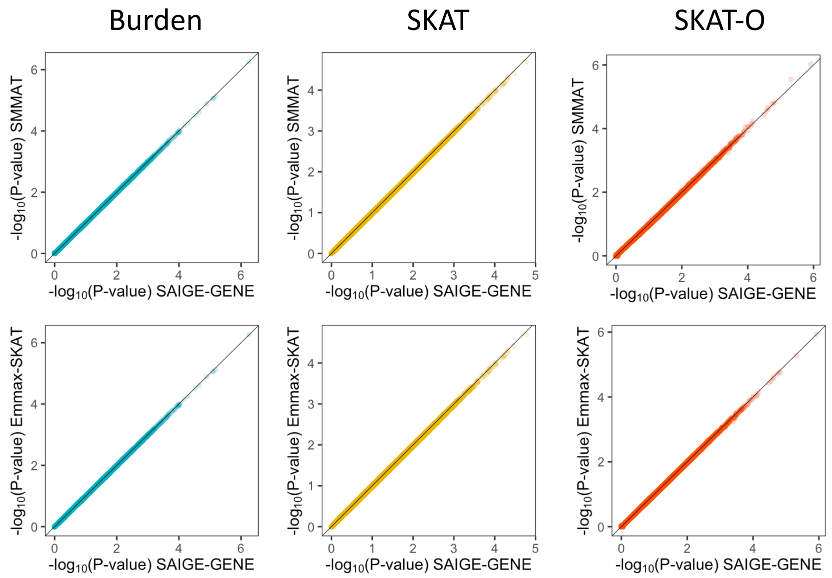


B. $h^2$= 0.4

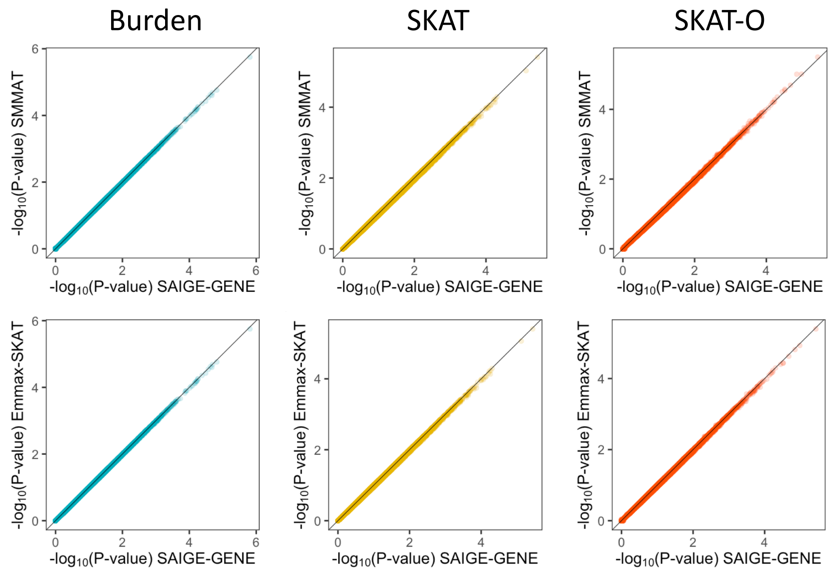

**Supplementary Figure 4.** Scatter plots of association p-values from SAIGE-GENE versus SMMAT and EmmaX-SKAT for the Burden, SKAT, and SKAT-O tests based on real data analysis on the -log10 scale. 12,000 genes were tested for A. automated read pulse rate using 20,000 randomly selected white British samples in the HRC-imputed UK Biobank; B. HDL using 20,000 randomly selected samples in HUNT. Missense and stop-gain variants with MAF ≤ 1% were included. The Pearson's correlation coefficients $r^2 > 0.99$ for -log10(P-values) between SAIGE and SMMAT and between SAIGE and EmmaX-SKAT.

A. automated read pulse rate in the UK Biobank

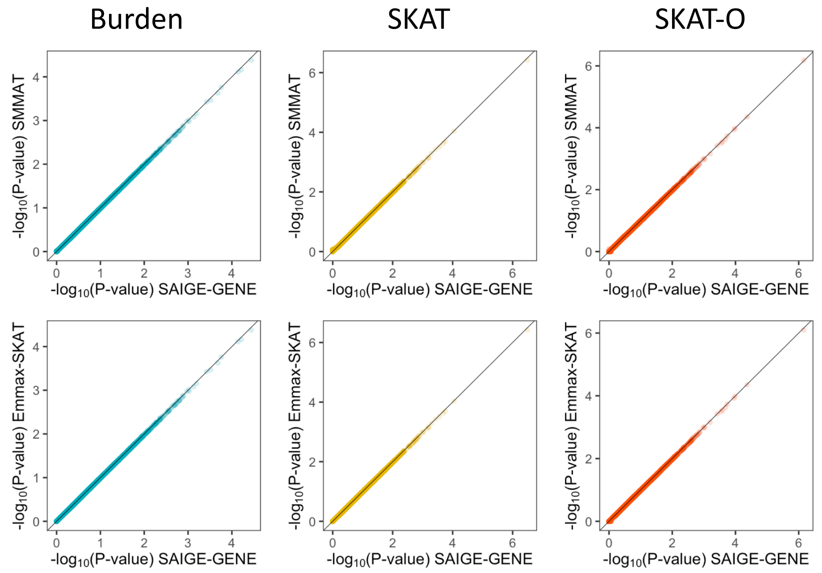


B. HDL in the HUNT study

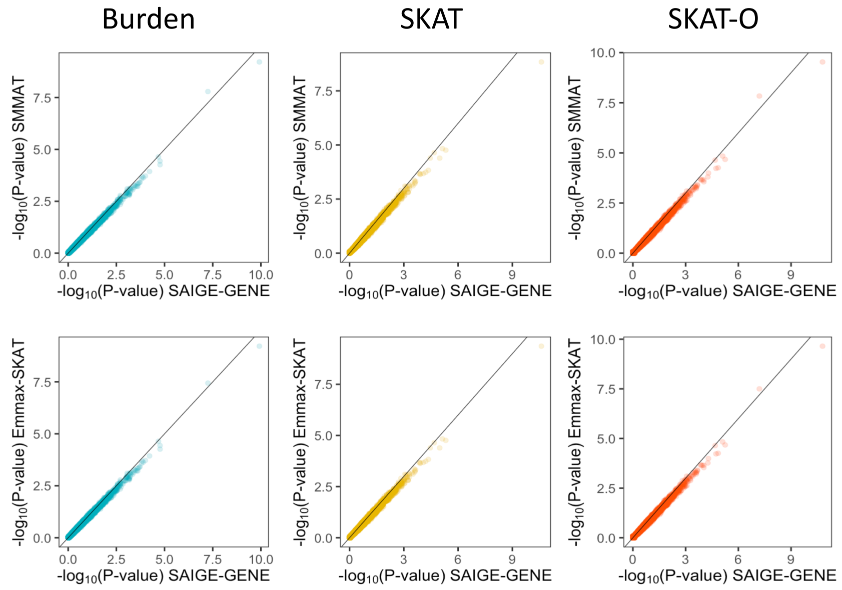

**Supplementary Figure 5.** Scatter plots of association p-values on the -log10 scale from SAIGE-GENE with two sample relatedness cutoffs for the sparse GRM, 0.125 and 0.2. 15,338 genes were tested for automated read pulse rate in white British samples in the HRC-imputed UK Biobank. Missense and stop-gain variants with MAF ≤ 1% were included. A. Burden. B. SKAT, C. SKAT-O

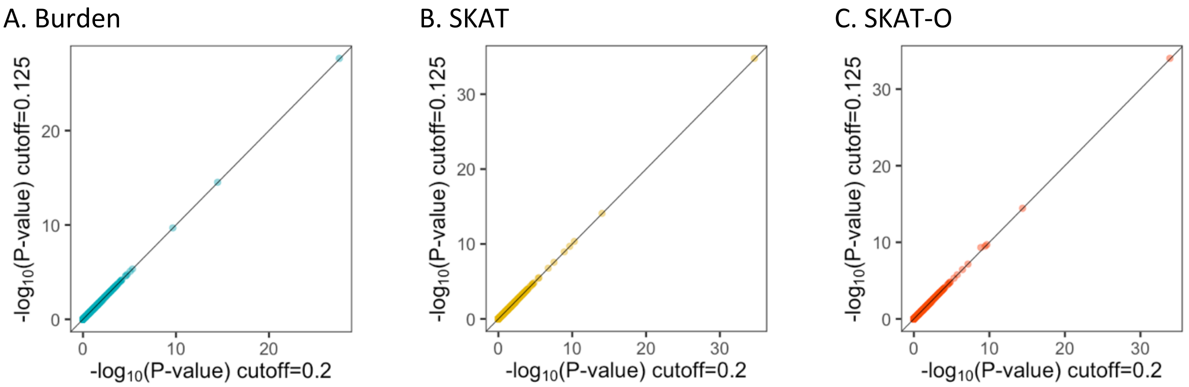


**Supplementary Figure 6.** Quantile-quantile plots of association p-values for 10 million variant sets from the simulation study for phenotypes with various case-control ratios.

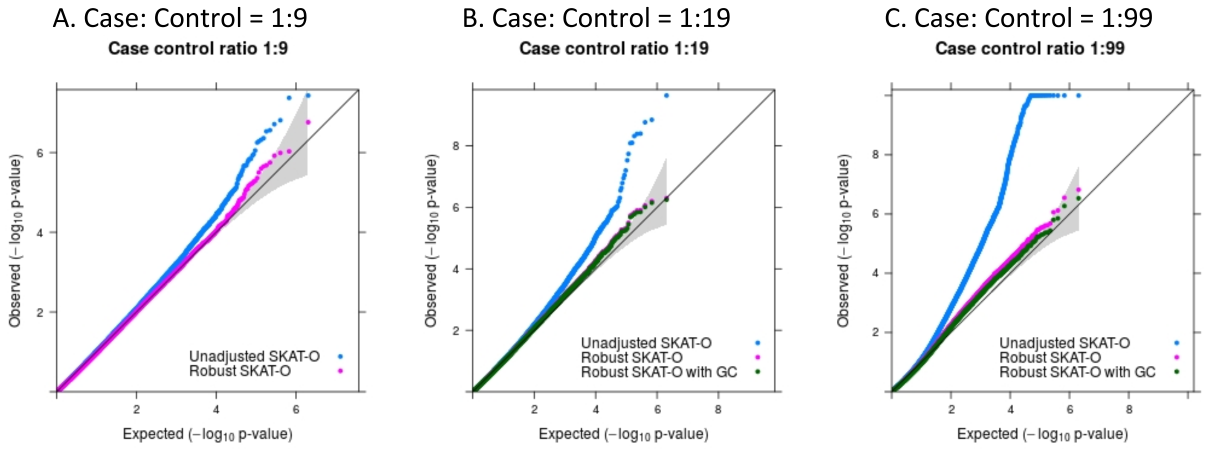

**Supplementary Figure 7.** Empirical computation time for A. step 1 for fitting a null mixed model and B. step 2 for association tests, respectively by sample sizes (N) for gene-based tests for 15,342 genes, each containing 50 rare variants. Benchmarking was performed on randomly sub-sampled UK Biobank data with 408,144 White British participants for waist-to-hip ratio. The reported run times and memory are medians of five runs with samples randomly selected from the full sample set using different sampling seeds. The reported computation time and memory for EmmaX-SKAT and SMMAT is the projected computation time when N > 20,000. As the number of tested markers varies by sample sizes, the computation time is projected for 50 markers per gene for plotting. Numerical data are provided in **Supplementary Table 1**.

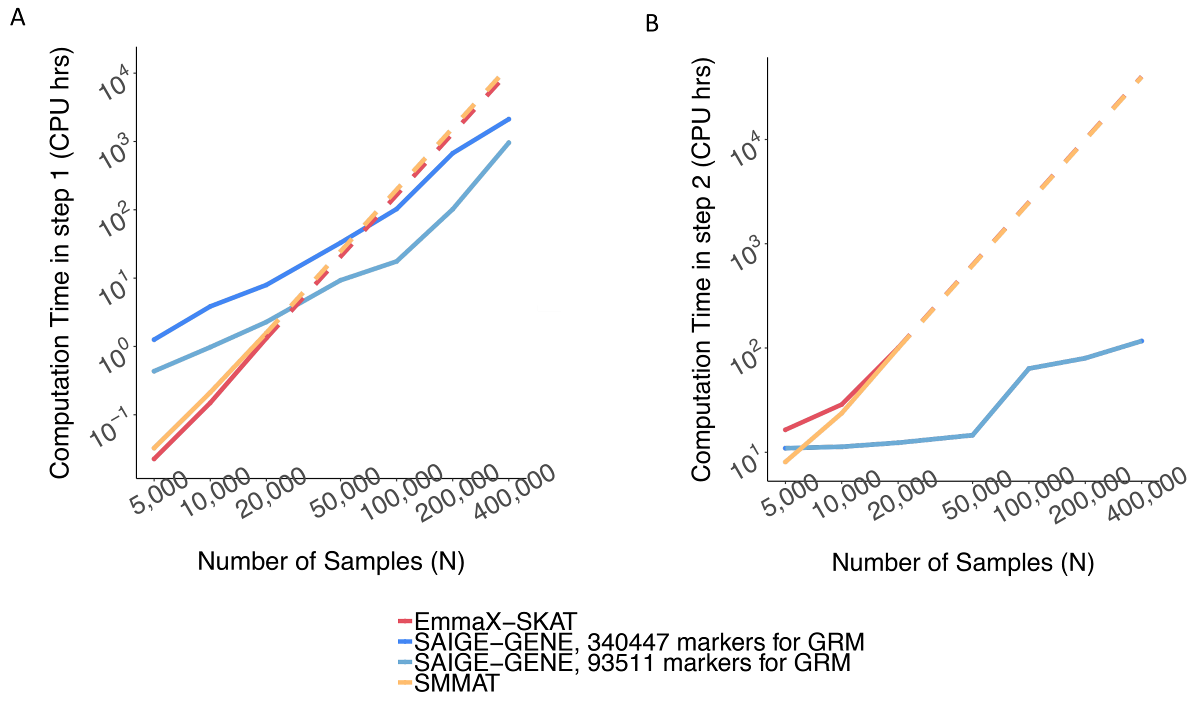

**Supplementary Figure 8.** Log-log plot of the estimated run time as a function of number of markers per gene. Benchmarking was performed on randomly sub-sampled 400,000 UK Biobank data with 408,144 white British participants for waist-to-hip ratio on 15,342 genes. Run times are medians of five runs with samples randomly selected from the full sample set using different sampling seeds. The computation time for other different number of markers per gene is projected based on the benchmarked time.

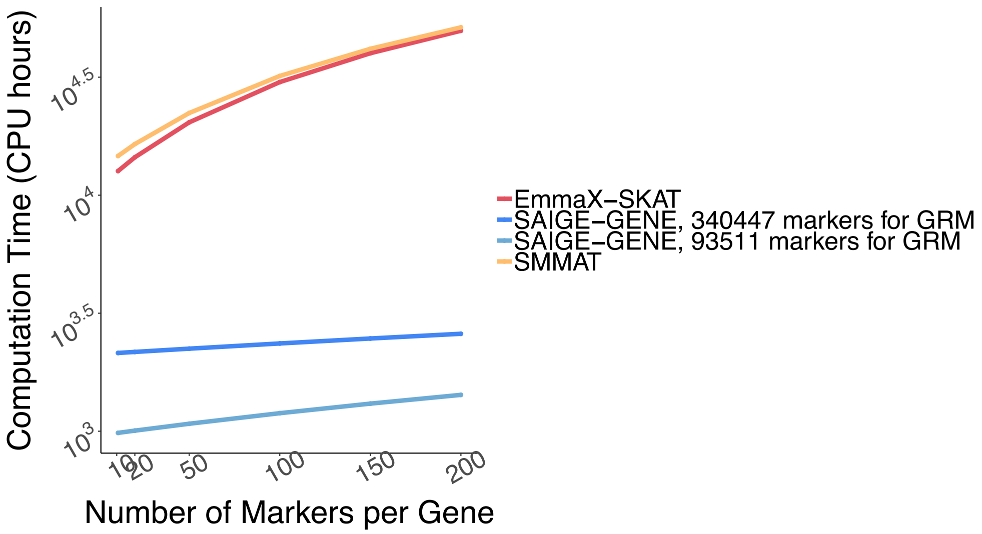

**Supplementary Figure 9.** Log-log plots of the estimated A. run time and B. memory usage as a function of sample size (N) for genome-wide tests for 286,000 chunks, each containing 50 variants on average, given that there are 14.3 million markers in the HRC-imputed UK Biobank with MAF ≤ 1% and imputation info score ≥ 0.8. Numerical data are provided in **Supplementary Table 1**. Benchmarking was performed on randomly sub-sampled UK Biobank data with 408,144 white British participants for waist-to-hip ratio. run times are medians of five runs with samples randomly selected from the full sample set using different sampling seeds.

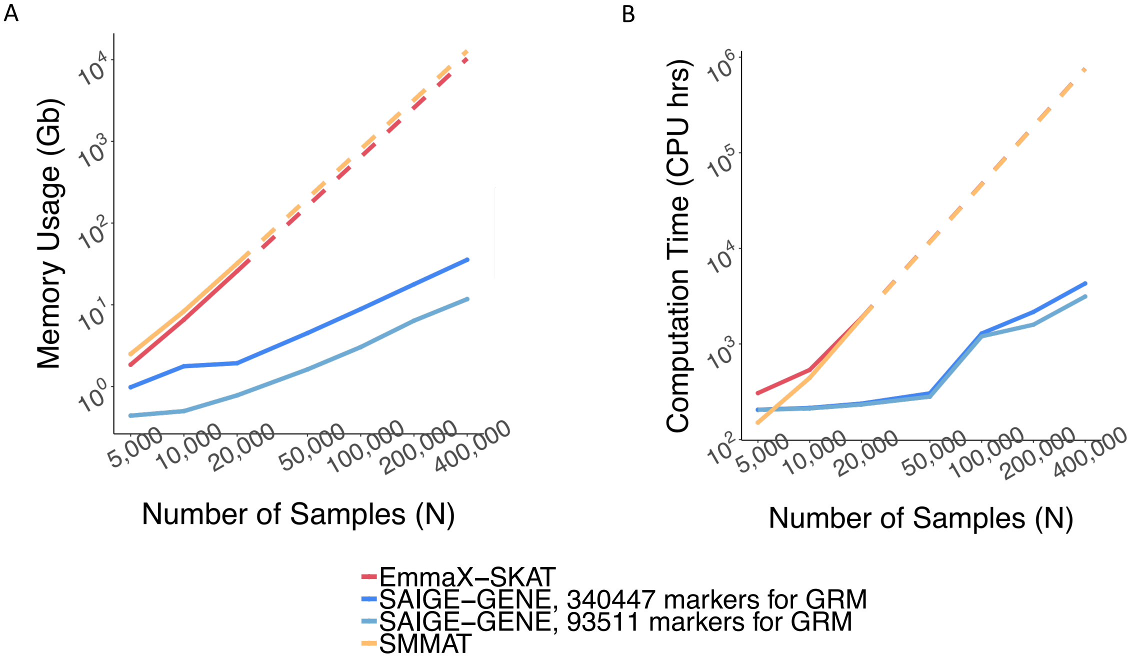

**Supplementary Figure 10.** Log-log plots of the estimated run time for as a function of sample size (N) for SAIGE-GENE with and without using the robust adjustment. A. Exome-wide gene-based tests for 15,871 genes and B. Genome-wide tests for 286,000 chunks. Each gene or chunk contains 50 variants on average. Benchmarking was performed on randomly sub-sampled UK Biobank data with 402,163 white British participants tested for glaucoma (PheCode: 365, 4,462 cases and 397,701 controls). The case-control ratio remained the same in subsampled data sets. The reported run times and memory are medians of five runs with samples randomly selected from the full sample set using different sampling seeds. As the number of tested markers varies by sample sizes, the computation time is projected for 50 markers per gene for plotting. Numerical data are provided in **Supplementary Table 2**.

**A.**

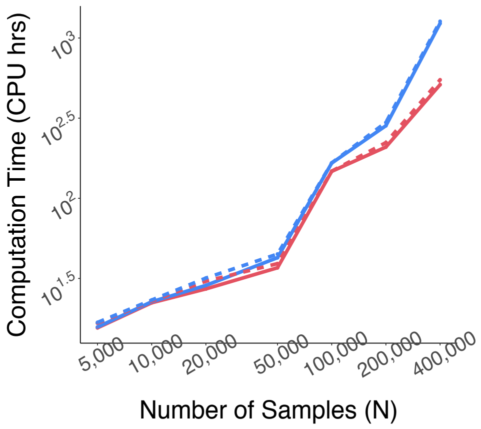


**B.**

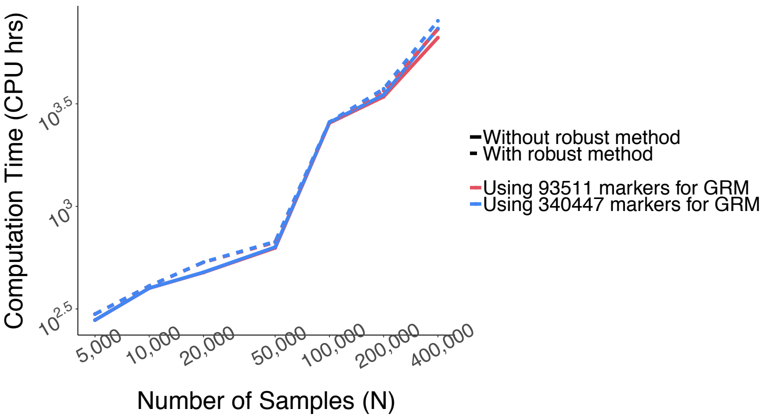

**Supplementary Figure 11.** Pedigree of families, each with 10 members, in the simulation study.

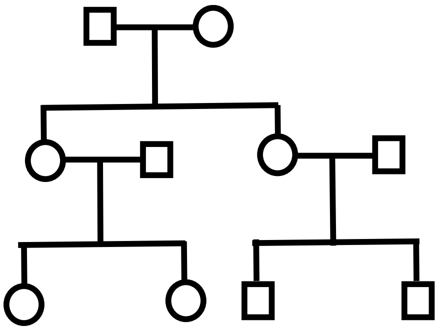

**Supplementary Figure 12.** Histogram of simulated phenotypes A. with skewed distribution. B. after inverse normal transformation as described in the subsection 2.2 of the section "Additional simulation and real-data analysis results"

A.

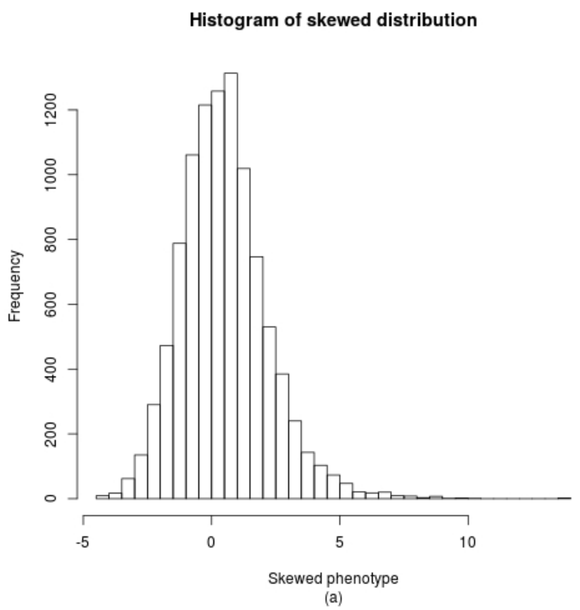


B.

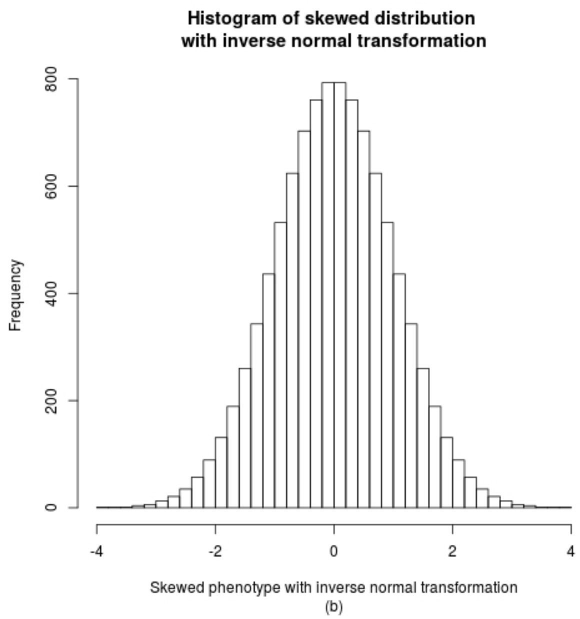

**Supplementary Figure 13.** Comparing heritability estimates using the sparse GRM to heritability estimates using the full GRM for 24 quantitative traits in the UK Biobank with sample size (N) ≥ 100,000. The sparse GRM was constructed using a coefficient of relatedness cutoff 0.125, corresponding to up to 3rd degree relative pairs.

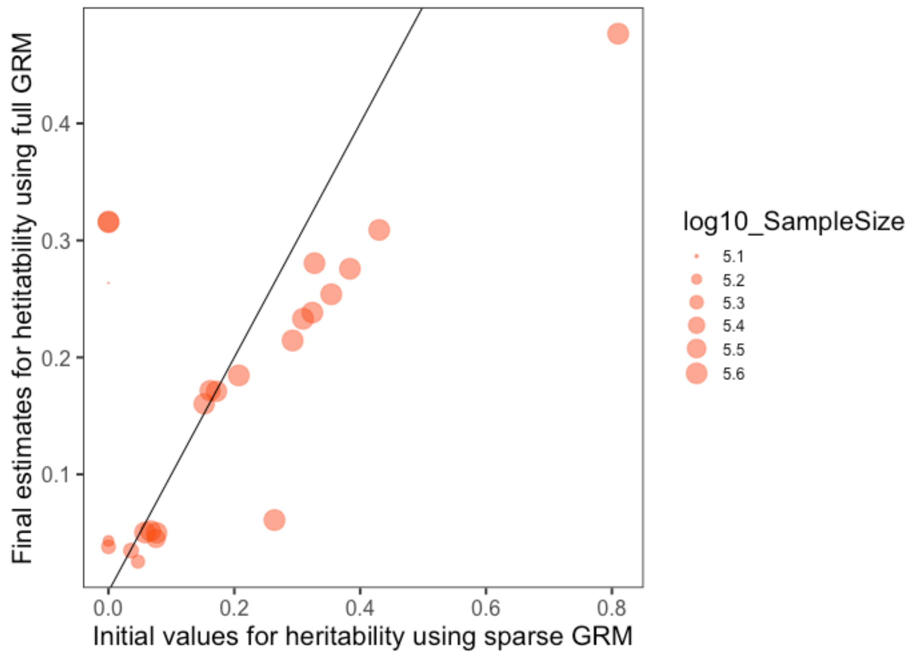

### 4. Supplementary tables

**Supplementary Table 1.** The estimated run time (A) and memory use (B) across different sample sizes. Benchmarking was performed on randomly sub-sampled UK Biobank data with 408,144 white British participants and 15,342 genes were tested for waist hip ratio. For simplicity, the number of markers in the gene was 50 regardless of sample sizes. The run times and memory are medians of five runs with samples randomly selected from the full sample set using different sampling seeds. The computation cost for genome-wide region-based tests were projected from the exome-wide gene-based tests results given that there are 14.3 million markers in the HRC-imputed UK Biobank with MAF $\leq$ 1% and imputation info score $\geq$ 0.8. Total 286,000 chunks are tested with 50 markers in each chunk.

| | sampleSize | step1(CPU hrs) | step2(CPU hrs) | Total Time(CPU hrs) | Memory (Gb) | Program |
|---|---|---|---|---|---|---|
| Exome-wide gene-based tests | 5,000 | 0.43 | 10.94 | 11.37 | 0.44 | SAIGE-GENE; 93,511 markers for GRM |
| | 10,000 | 0.98 | 11.29 | 12.26 | 0.50 | SAIGE-GENE; 93,511 markers for GRM |
| | 20,000 | 2.28 | 12.34 | 14.61 | 0.78 | SAIGE-GENE; 93,511 markers for GRM |
| | 50,000 | 9.29 | 14.57 | 23.87 | 1.62 | SAIGE-GENE; 93,511 markers for GRM |
| | 100,000 | 17.58 | 63.51 | 81.09 | 3.04 | SAIGE-GENE; 93,511 markers for GRM |
| | 200,000 | 101.85 | 79.87 | 181.71 | 6.39 | SAIGE-GENE; 93,511 markers for GRM |
| | 400,000 | 958.98 | 116.65 | 1075.63 | 11.74 | SAIGE-GENE; 93,511 markers for GRM |
| | 5,000 | 1.26 | 10.94 | 12.20 | 0.98 | SAIGE-GENE; 340,447 markers for GRM |
| | 10,000 | 3.85 | 11.29 | 15.14 | 1.77 | SAIGE-GENE; 340,447 markers for GRM |
| | 20,000 | 7.95 | 12.34 | 20.28 | 1.93 | SAIGE-GENE; 340,447 markers for GRM |
| | 50,000 | 32.88 | 14.57 | 47.45 | 4.51 | SAIGE-GENE; 340,447 markers for GRM |
| | 100,000 | 102.71 | 63.51 | 166.22 | 8.87 | SAIGE-GENE; 340,447 markers for GRM |
| | 200,000 | 672.29 | 79.87 | 752.16 | 17.82 | SAIGE-GENE; 340,447 markers for GRM |
| | 400,000 | 2120.89 | 116.65 | 2237.54 | 35.59 | SAIGE-GENE; 340,447 markers for GRM |
| | 5,000 | 0.02 | 16.42 | 16.45 | 1.85 | EmmaX-SKAT |
| | 10,000 | 0.15 | 28.72 | 28.87 | 6.57 | EmmaX-SKAT |
| | 20,000 | 1.32 | 100.32 | 101.64 | 25.82 | EmmaX-SKAT |
| | 50,000 | 20.62 | 626.98 | 647.60 | 161.37 | EmmaX-SKAT |
| | 100,000 | 164.93 | 2507.93 | 2672.86 | 645.50 | EmmaX-SKAT |
| | 200,000 | 1319.44 | 10031.73 | 11351.17 | 2581.99 | EmmaX-SKAT |
| | 400,000 | 10555.55 | 40126.91 | 50682.46 | 10327.96 | EmmaX-SKAT |
| | 5,000 | 0.03 | 8.07 | 8.11 | 2.50 | SMMAT |
| | 10,000 | 0.21 | 23.68 | 23.90 | 8.34 | SMMAT |
| | 20,000 | 1.57 | 99.44 | 101.01 | 32.03 | SMMAT |
| | 50,000 | 24.57 | 621.47 | 646.04 | 200.22 | SMMAT |
| | 100,000 | 196.56 | 2485.89 | 2682.46 | 800.87 | SMMAT |
| | 200,000 | 1572.50 | 9943.57 | 11516.07 | 3203.49 | SMMAT |
| | 400,000 | 12580.00 | 39774.28 | 52354.28 | 12813.95 | SMMAT |
| Genome-wide region-based tests | 5,000 | 0.43 | 203.89 | 204.33 | 0.44 | SAIGE-GENE; 93,511 markers for GRM |
| | 10,000 | 0.98 | 210.42 | 211.40 | 0.50 | SAIGE-GENE; 93,511 markers for GRM |
| | 20,000 | 2.28 | 229.96 | 232.24 | 0.78 | SAIGE-GENE; 93,511 markers for GRM |
| | 50,000 | 9.29 | 271.67 | 280.97 | 1.62 | SAIGE-GENE; 93,511 markers for GRM |
| | 100,000 | 17.58 | 1183.98 | 1201.56 | 3.04 | SAIGE-GENE; 93,511 markers for GRM |
| | 200,000 | 101.85 | 1488.86 | 1590.70 | 6.39 | SAIGE-GENE; 93,511 markers for GRM |
| | 400,000 | 958.98 | 2174.55 | 3133.53 | 11.74 | SAIGE-GENE; 93,511 markers for GRM |
| | 5,000 | 1.26 | 203.89 | 205.15 | 0.98 | SAIGE-GENE; 340,447 markers for GRM |
| | 10,000 | 3.85 | 210.42 | 214.27 | 1.77 | SAIGE-GENE; 340,447 markers for GRM |
| | 20,000 | 7.95 | 229.96 | 237.90 | 1.93 | SAIGE-GENE; 340,447 markers for GRM |
| | 50,000 | 32.88 | 271.67 | 304.55 | 4.51 | SAIGE-GENE; 340,447 markers for GRM |
| | 100,000 | 102.71 | 1183.98 | 1286.69 | 8.87 | SAIGE-GENE; 340,447 markers for GRM |
| | 200,000 | 672.29 | 1488.86 | 2161.15 | 17.82 | SAIGE-GENE; 340,447 markers for GRM |

| | | | | | | |
|---|---|---|---|---|---|---|
| | 400,000 | 2120.89 | 2174.55 | 4295.44 | 35.59 | SAIGE-GENE; 340,447 markers for GRM |
| | 5,000 | 0.02 | 306.15 | 306.17 | 1.85 | EmmaX-SKAT |
| | 10,000 | 0.15 | 535.40 | 535.55 | 6.57 | EmmaX-SKAT |
| | 20,000 | 1.32 | 1870.08 | 1871.40 | 25.82 | EmmaX-SKAT |
| | 50,000 | 20.62 | 11687.99 | 11708.61 | 161.37 | EmmaX-SKAT |
| | 100,000 | 164.93 | 46751.96 | 46916.89 | 645.50 | EmmaX-SKAT |
| | 200,000 | 1319.44 | 187007.85 | 188327.29 | 2581.99 | EmmaX-SKAT |
| | 400,000 | 10555.55 | 748031.39 | 758586.94 | 10327.96 | EmmaX-SKAT |
| | 5,000 | 0.03 | 150.51 | 150.55 | 2.50 | SMMAT |
| | 10,000 | 0.21 | 441.50 | 441.71 | 8.34 | SMMAT |
| | 20,000 | 1.57 | 1853.64 | 1855.22 | 32.03 | SMMAT |
| | 50,000 | 24.57 | 11585.28 | 11609.85 | 200.22 | SMMAT |
| | 100,000 | 196.56 | 46341.12 | 46537.68 | 800.87 | SMMAT |
| | 200,000 | 1572.50 | 185364.46 | 186936.96 | 3203.49 | SMMAT |
| | 400,000 | 12580.00 | 741457.85 | 754037.85 | 12813.95 | SMMAT |

**Supplementary Table 2.** The estimated run time (A) and memory use (B) across different sample sizes for binary traits with and without the robust adjustment. Benchmarking was performed on randomly sub-sampled UK Biobank data with 402,163 white British participants and 15,871 genes were tested for glaucoma (PheCode: 365). Samples were randomly selected from 4,462 glaucoma cases and 397,701 controls respectively, so the case-control ratio remained the same in sub-sampled data sets. For simplicity, the number of markers in the gene was 50 regardless of sample sizes. The run times and memory are medians of five runs with samples randomly selected from the full sample set using different sampling seeds. The computation cost for genome-wide region-based tests were projected from the exome-wide gene-based tests results given that there are 14.3 million markers in the HRC-imputed UK Biobank with MAF $\leq$ 1% and imputation info score $\geq$ 0.8. Total 286,000 chunks are tested with 50 markers in each chunk.

| | sampleSize | step1(CPU hrs) | step2(CPU hrs) | Total Time(CPU hrs) | Memory (Gb) | Program |
|---|---|---|---|---|---|---|
| Exome-wide gene-based tests | 5,000 | 0.09 | 15.47 | 15.56 | 0.35 | SAIGE-GENE without robust adjustment; 93,511 markers for GRM |
| | 10,000 | 0.16 | 22.14 | 22.30 | 0.47 | SAIGE-GENE without robust adjustment; 93,511 markers for GRM |
| | 20,000 | 0.84 | 26.37 | 27.21 | 0.73 | SAIGE-GENE without robust adjustment; 93,511 markers for GRM |
| | 50,000 | 2.18 | 34.69 | 36.87 | 2.68 | SAIGE-GENE without robust adjustment; 93,511 markers for GRM |
| | 100,000 | 6.11 | 141.10 | 147.21 | 2.97 | SAIGE-GENE without robust adjustment; 93,511 markers for GRM |
| | 200,000 | 20.05 | 188.79 | 208.84 | 6.39 | SAIGE-GENE without robust adjustment; 93,511 markers for GRM |
| | 400,000 | 151.22 | 360.24 | 511.46 | 12.76 | SAIGE-GENE without robust adjustment; 93,511 markers for GRM |
| | 5,000 | 0.09 | 16.55 | 16.64 | 0.35 | SAIGE-GENE with robust adjustment; 93,511 markers for GRM |
| | 10,000 | 0.16 | 22.68 | 22.84 | 0.47 | SAIGE-GENE with robust adjustment; 93,511 markers for GRM |
| | 20,000 | 0.84 | 29.55 | 30.39 | 0.73 | SAIGE-GENE with robust adjustment; 93,511 markers for GRM |
| | 50,000 | 2.18 | 36.94 | 39.12 | 2.68 | SAIGE-GENE with robust adjustment; 93,511 markers for GRM |
| | 100,000 | 6.11 | 141.58 | 147.69 | 2.97 | SAIGE-GENE with robust adjustment; 93,511 markers for GRM |
| | 200,000 | 20.05 | 202.43 | 222.48 | 6.39 | SAIGE-GENE with robust adjustment; 93,511 markers for GRM |
| | 400,000 | 151.22 | 396.94 | 548.16 | 12.76 | SAIGE-GENE with robust adjustment; 93,511 markers for GRM |
| | 5,000 | 0.26 | 15.47 | 15.73 | 0.98 | SAIGE-GENE without robust adjustment; 340,447 markers for GRM |
| | 10,000 | 0.49 | 22.14 | 22.63 | 1.06 | SAIGE-GENE without robust adjustment; 340,447 markers for GRM |
| | 20,000 | 2.24 | 26.37 | 28.61 | 3.39 | SAIGE-GENE without robust adjustment; 340,447 markers for GRM |
| | 50,000 | 7.93 | 34.69 | 42.62 | 4.44 | SAIGE-GENE without robust adjustment; 340,447 markers for GRM |
| | 100,000 | 24.68 | 141.10 | 165.79 | 8.94 | SAIGE-GENE without robust adjustment; 340,447 markers for GRM |
| | 200,000 | 94.84 | 188.79 | 283.63 | 17.51 | SAIGE-GENE without robust adjustment; 340,447 markers for GRM |
| | 400,000 | 872.30 | 360.24 | 1232.54 | 35.38 | SAIGE-GENE without robust adjustment; 340,447 markers for GRM |
| | 5,000 | 0.26 | 16.55 | 16.81 | 0.98 | SAIGE-GENE with robust adjustment; 340,447 markers for GRM |
| | 10,000 | 0.49 | 22.68 | 23.18 | 1.06 | SAIGE-GENE with robust adjustment; 340,447 markers for GRM |
| | 20,000 | 2.24 | 29.55 | 31.79 | 3.39 | SAIGE-GENE with robust adjustment; 340,447 markers for GRM |
| | 50,000 | 7.93 | 36.94 | 44.87 | 4.44 | SAIGE-GENE with robust adjustment; 340,447 markers for GRM |
| | 100,000 | 24.68 | 141.58 | 166.27 | 8.94 | SAIGE-GENE with robust adjustment; 340,447 markers for GRM |
| | 200,000 | 94.84 | 202.43 | 297.26 | 17.51 | SAIGE-GENE with robust adjustment; 340,447 markers for GRM |
| | 400,000 | 872.30 | 396.94 | 1269.25 | 35.38 | SAIGE-GENE with robust adjustment; 340,447 markers for GRM |
| Genome-wide region-based tests | 5,000 | 0.09 | 278.79 | 278.88 | 0.35 | SAIGE-GENE without robust adjustment; 93,511 markers for GRM |
| | 10,000 | 0.16 | 398.98 | 399.14 | 0.47 | SAIGE-GENE without robust adjustment; 93,511 markers for GRM |
| | 20,000 | 0.84 | 475.16 | 476.00 | 0.73 | SAIGE-GENE without robust adjustment; 93,511 markers for GRM |
| | 50,000 | 2.18 | 625.10 | 627.28 | 2.68 | SAIGE-GENE without robust adjustment; 93,511 markers for GRM |
| | 100,000 | 6.11 | 2542.81 | 2548.91 | 2.97 | SAIGE-GENE without robust adjustment; 93,511 markers for GRM |
| | 200,000 | 20.05 | 3402.16 | 3422.22 | 6.39 | SAIGE-GENE without robust adjustment; 93,511 markers for GRM |
| | 400,000 | 151.22 | 6491.80 | 6643.02 | 12.76 | SAIGE-GENE without robust adjustment; 93,511 markers for GRM |
| | 5,000 | 0.09 | 298.30 | 298.39 | 0.35 | SAIGE-GENE with robust adjustment; 93,511 markers for GRM |
| | 10,000 | 0.16 | 408.80 | 408.96 | 0.47 | SAIGE-GENE with robust adjustment; 93,511 markers for GRM |
| | 20,000 | 0.84 | 532.50 | 533.34 | 0.73 | SAIGE-GENE with robust adjustment; 93,511 markers for GRM |
| | 50,000 | 2.18 | 665.67 | 667.85 | 2.68 | SAIGE-GENE with robust adjustment; 93,511 markers for GRM |

| | | | | | | |
|---|---|---|---|---|---|---|
| | 100,000 | 6.11 | 2551.42 | 2557.53 | 2.97 | SAIGE-GENE with robust adjustment; 93,511 markers for GRM |
| | 200,000 | 20.05 | 3647.87 | 3667.92 | 6.39 | SAIGE-GENE with robust adjustment; 93,511 markers for GRM |
| | 400,000 | 151.22 | 7153.23 | 7304.45 | 12.76 | SAIGE-GENE with robust adjustment; 93,511 markers for GRM |
| | 5,000 | 0.26 | 278.79 | 279.05 | 0.98 | SAIGE-GENE without robust adjustment; 340,447 markers for GRM |
| | 10,000 | 0.49 | 398.98 | 399.47 | 1.06 | SAIGE-GENE without robust adjustment; 340,447 markers for GRM |
| | 20,000 | 2.24 | 475.16 | 477.40 | 3.39 | SAIGE-GENE without robust adjustment; 340,447 markers for GRM |
| | 50,000 | 7.93 | 625.10 | 633.03 | 4.44 | SAIGE-GENE without robust adjustment; 340,447 markers for GRM |
| | 100,000 | 24.68 | 2542.81 | 2567.49 | 8.94 | SAIGE-GENE without robust adjustment; 340,447 markers for GRM |
| | 200,000 | 94.84 | 3402.16 | 3497.00 | 17.51 | SAIGE-GENE without robust adjustment; 340,447 markers for GRM |
| | 400,000 | 872.30 | 6491.80 | 7364.11 | 35.38 | SAIGE-GENE without robust adjustment; 340,447 markers for GRM |
| | 5,000 | 0.26 | 298.30 | 298.56 | 0.98 | SAIGE-GENE with robust adjustment; 340,447 markers for GRM |
| | 10,000 | 0.49 | 408.80 | 409.29 | 1.06 | SAIGE-GENE with robust adjustment; 340,447 markers for GRM |
| | 20,000 | 2.24 | 532.50 | 534.75 | 3.39 | SAIGE-GENE with robust adjustment; 340,447 markers for GRM |
| | 50,000 | 7.93 | 665.67 | 673.60 | 4.44 | SAIGE-GENE with robust adjustment; 340,447 markers for GRM |
| | 100,000 | 24.68 | 2551.42 | 2576.10 | 8.94 | SAIGE-GENE with robust adjustment; 340,447 markers for GRM |
| | 200,000 | 94.84 | 3647.87 | 3742.70 | 17.51 | SAIGE-GENE with robust adjustment; 340,447 markers for GRM |
| | 400,000 | 872.30 | 7153.23 | 8025.54 | 35.38 | SAIGE-GENE with robust adjustment; 340,447 markers for GRM |

**Supplementary Table 3.** Heritability estimated based on the full GRM by step 1 in SAIGE-GENE A. for 53 quantitative traits and B. for 10 binary traits in the UK Biobank. For quantitative traits, $\hat{h}^2 = \hat{\tau}/(\hat{\phi} + \hat{\tau})$, where $\hat{\tau}$ is the additive genetic variance parameter estimate and $\hat{\phi}$ is the variance parameter estimate for the error term (see **ONLINE METHODS**) from step 1 in SAIGE-GENE. For binary traits, heritability estimates in a liability scale, $\hat{h}^2_{latent}$, is reported here. The $\hat{h}^2_{latent}$ was obtained using the fact that the logistic regression can be described as a liability threshold model with standard logistic distribution, which has variance=$\pi^2/3$ = 3.23. $\hat{h}^2_{latent} = \frac{\hat{\tau}}{\pi^2/3 + \hat{\tau}}$, where $\hat{\tau}$ is the additive genetic variance parameter estimate from step 1 in SAIGE-GENE. Since GRM was constructed using variants with MAF ≥ 0.01, the estimated heritability is the narrow sense of the heritability due to common variants.

**A.**

| Phenotype | Sample Size | $\hat{h}^2$ |
|---|---|---|
| Townsend_deprivation_index | 408422 | 0.061 |
| Waist_circumference | 408227 | 0.214 |
| Hip_circumference | 408182 | 0.233 |
| Waist_hip_ratio | 408144 | 0.185 |
| Height_standing | 408034 | 0.477 |
| Body_mass_index | 407605 | 0.254 |
| Days_per_week_walked_10min | 402016 | 0.050 |
| Weight | 401786 | 0.276 |
| Whole_body_water_mass | 401782 | 0.315 |
| Basal_metabolic_rate | 401771 | 0.309 |
| Whole_body_fat_free_mass | 401747 | 0.316 |
| Body_fat_percentage | 401556 | 0.238 |
| Days_week_vigorous_phys_activity_10min | 389393 | 0.050 |
| Days_per_week_moderate_phys_activity_10min | 389204 | 0.052 |
| Blood_pressure_diastolic_automated_mean | 385365 | 0.160 |
| Pulse_rate_automated_mean | 385365 | 0.171 |
| Blood_pressure_systolic_automated_mean | 385362 | 0.172 |
| Duration_moderate_activity | 303663 | 0.045 |
| Duration_of_vigorous_activity | 221867 | 0.035 |
| Duration_of_light_DIY | 203191 | 0.038 |
| Duration_of_other_exercises | 194610 | 0.025 |
| Duration_of_heavy_DIY | 165319 | 0.043 |
| Smoking_packyear | 124011 | 0.264 |
| Age_high_blood_pressure | 99296 | 0.106 |
| Duration_of_strenuous_sports | 41133 | 0.032 |
| Blood_pressure_diastolic_manual_mean | 35283 | 0.149 |
| Blood_pressure_systolic_manual_mean | 35283 | 0.156 |
| Age_diabetes | 18552 | 0.187 |
| Age_at_death | 11928 | 0.000 |
| Heart_rate | 8421 | 0.208 |
| P_wave_duration | 8421 | 0.296 |
| QRS_duration | 8421 | 0.337 |
| Average_heart_rate_MRI | 3875 | 0.231 |
| Body_surface_area_MRI | 3875 | 0.521 |
| Cardiac_index | 3875 | 0.083 |
| Cardiac_output | 3875 | 0.154 |
| LV_ejection_fraction | 3875 | 0.504 |

| | | |
|---|---|---|
| LV_end_diastolic_volume | 3875 | 0.650 |
| LV_end_systolic_volume | 3875 | 0.522 |
| LV_stroke_volume | 3875 | 0.681 |
| Maximum_carotid_IMT_150 | 2221 | 0.135 |
| Mean_carotid_IMT_150 | 2221 | 0.379 |
| Minimum_carotid_IMT_150 | 2221 | 0.590 |
| Maximum_carotid_IMT_210 | 2209 | 0.830 |
| Mean_carotid_IMT_210 | 2209 | 0.790 |
| Minimum_carotid_IMT_210 | 2209 | 0.678 |
| Maximum_carotid_IMT_120 | 2204 | 0.000 |
| Mean_carotid_IMT_120 | 2204 | 0.000 |
| Minimum_carotid_IMT_120 | 2204 | 0.000 |
| Maximum_carotid_IMT_240 | 2183 | 0.178 |
| Mean_carotid_IMT_240 | 2183 | 0.300 |
| Minimum_carotid_IMT_240 | 2183 | 0.000 |

**B.**

| **Phenotype** | **Phecode** | **Number of cases** | **Number of controls** | $\widehat{h}^2_{latent}$ |
|---|---|---|---|---|
| Hematuria | 593 | 16409 | 379936 | 0.074 |
| Cholelithiasis and cholecystitis | 574 | 16225 | 391307 | 0.121 |
| Prostate cancer | 185 | 6743 | 169185 | 0.082 |
| Diseases of hair and hair follicles | 704 | 5344 | 402357 | 0.156 |
| Colorectal cancer | 153 | 4562 | 382756 | 0.135 |
| Glaucoma | 365 | 4462 | 397761 | 0.133 |
| Pulmonary heart disease | 415 | 4257 | 402375 | 0.111 |
| Melanomas of skin, dx or hx | 172.1 | 2691 | 395071 | 0.161 |
| Ankylosing spondylitis | 715.2 | 620 | 365085 | 0.000 |
| Thyroid cancer | 193 | 358 | 407399 | 0.000 |

**Supplementary Table 4.** Exome-wide significant genes with p-values $\leq 2.5\text{x}10^{-6}$ identified by SAIGE-GENE in the UK Biobank a. for 53 quantitative traits, b for 10 binary traits with various case-control ratios

**A.**

| Phenotype | Gene | P-value | Number of variants | Sample Size |
|---|---|---|---|---|
| Waist circumference | *GPR151* | 2.15E-10 | 6 | 408227 |
| Waist circumference | *C16orf70* | 1.94E-06 | 2 | 408227 |
| Hip circumference | *SYPL2* | 2.91E-08 | 4 | 408182 |
| Hip circumference | *TRAPPC4* | 5.83E-07 | 2 | 408182 |
| Hip circumference | *ANO1* | 5.98E-07 | 4 | 408182 |
| Hip circumference | *GPR151* | 6.08E-07 | 6 | 408182 |
| Hip circumference | *C16orf70* | 1.14E-06 | 2 | 408182 |
| Waist hip ratio | *TAS2R46* | 1.36E-08 | 2 | 408144 |
| Waist hip ratio | *GPR151* | 3.00E-08 | 6 | 408144 |
| Waist hip ratio | *SLC5A3* | 1.33E-07 | 4 | 408144 |
| Height standing | *SCMH1* | 3.41E-39 | 8 | 408034 |
| Height standing | *ACAN* | 1.39E-36 | 33 | 408034 |
| Height standing | *FBN2* | 3.15E-32 | 16 | 408034 |
| Height standing | *ZFAT* | 3.61E-32 | 13 | 408034 |
| Height standing | *HTRA1* | 1.66E-29 | 4 | 408034 |
| Height standing | *NPR3* | 3.60E-25 | 3 | 408034 |
| Height standing | *STC2* | 1.10E-24 | 4 | 408034 |
| Height standing | *SPSB3* | 1.10E-22 | 4 | 408034 |
| Height standing | *NUBP2* | 2.01E-22 | 9 | 408034 |
| Height standing | *ATAD2* | 8.07E-21 | 6 | 408034 |
| Height standing | *ADAMTS6* | 1.83E-17 | 6 | 408034 |
| Height standing | *GRAMD2A* | 3.41E-16 | 4 | 408034 |
| Height standing | *GRM4* | 7.23E-16 | 2 | 408034 |
| Height standing | *MTMR11* | 1.94E-15 | 3 | 408034 |
| Height standing | *CRISPLD2* | 2.04E-14 | 11 | 408034 |
| Height standing | *CERCAM* | 7.05E-14 | 10 | 408034 |
| Height standing | *PDE3B* | 1.88E-13 | 8 | 408034 |
| Height standing | *FNDC3B* | 1.39E-12 | 8 | 408034 |
| Height standing | *PKD1* | 6.40E-12 | 43 | 408034 |
| Height standing | *FER* | 6.68E-12 | 5 | 408034 |
| Height standing | *C16orf70* | 7.45E-12 | 2 | 408034 |
| Height standing | *HAPLN3* | 1.25E-11 | 6 | 408034 |
| Height standing | *ST3GAL4* | 5.37E-11 | 5 | 408034 |
| Height standing | *SHANK1* | 8.09E-11 | 5 | 408034 |
| Height standing | *TRAPPC13* | 1.04E-10 | 2 | 408034 |
| Height standing | *S1PR5* | 1.44E-10 | 5 | 408034 |
| Height standing | *PTCH1* | 2.65E-10 | 15 | 408034 |
| Height standing | *COL8A1* | 3.25E-10 | 2 | 408034 |
| Height standing | *EXD1* | 3.84E-10 | 3 | 408034 |
| Height standing | *ATAD5* | 4.74E-10 | 8 | 408034 |
| Height standing | *ESR1* | 7.28E-10 | 9 | 408034 |
| Height standing | *CLEC3A* | 9.08E-10 | 3 | 408034 |
| Height standing | *PTH1R* | 1.40E-09 | 3 | 408034 |
| Height standing | *FGFR3* | 3.50E-09 | 4 | 408034 |
| Height standing | *NOX4* | 4.09E-09 | 4 | 408034 |
| Height standing | *CYR61* | 4.40E-09 | 2 | 408034 |
| Height standing | *TBX3* | 4.59E-09 | 2 | 408034 |
| Height standing | *SAMD4A* | 4.77E-09 | 7 | 408034 |
| Height standing | *ZCCHC6* | 1.42E-08 | 7 | 408034 |

| | | | | |
|---|---|---|---|---|
| Height standing | *CTU2* | 2.09E-08 | 12 | 408034 |
| Height standing | *LPP* | 2.11E-08 | 8 | 408034 |
| Height standing | *LRRC8A* | 2.87E-08 | 2 | 408034 |
| Height standing | *DGKH* | 3.08E-08 | 9 | 408034 |
| Height standing | *ABCB6* | 4.74E-08 | 11 | 408034 |
| Height standing | *PIEZO1* | 6.63E-08 | 55 | 408034 |
| Height standing | *ELN* | 6.67E-08 | 8 | 408034 |
| Height standing | *ATG10* | 1.35E-07 | 4 | 408034 |
| Height standing | *CADM1* | 1.35E-07 | 2 | 408034 |
| Height standing | *CPPED1* | 1.40E-07 | 7 | 408034 |
| Height standing | *PFDN2* | 1.41E-07 | 2 | 408034 |
| Height standing | *PROP1* | 1.63E-07 | 4 | 408034 |
| Height standing | *PRSS56* | 1.68E-07 | 3 | 408034 |
| Height standing | *GYG1* | 1.78E-07 | 5 | 408034 |
| Height standing | *CHGA* | 2.20E-07 | 9 | 408034 |
| Height standing | *SERPINE2* | 2.80E-07 | 3 | 408034 |
| Height standing | *SPATA5* | 2.83E-07 | 5 | 408034 |
| Height standing | *AMOTL1* | 2.85E-07 | 6 | 408034 |
| Height standing | *TMEM150B* | 2.88E-07 | 3 | 408034 |
| Height standing | *COL11A1* | 2.91E-07 | 8 | 408034 |
| Height standing | *PTK7* | 2.95E-07 | 6 | 408034 |
| Height standing | *PHC3* | 3.24E-07 | 7 | 408034 |
| Height standing | *TARS2* | 4.22E-07 | 3 | 408034 |
| Height standing | *VASN* | 5.25E-07 | 7 | 408034 |
| Height standing | *TXLNA* | 5.66E-07 | 3 | 408034 |
| Height standing | *FAM76A* | 5.70E-07 | 3 | 408034 |
| Height standing | *C11orf57* | 5.71E-07 | 2 | 408034 |
| Height standing | *FBLN2* | 6.61E-07 | 13 | 408034 |
| Height standing | *MC3R* | 6.92E-07 | 2 | 408034 |
| Height standing | *GTF2E2* | 7.05E-07 | 3 | 408034 |
| Height standing | *FIBIN* | 1.09E-06 | 2 | 408034 |
| Height standing | *PREB* | 1.12E-06 | 5 | 408034 |
| Height standing | *SIX6* | 1.16E-06 | 3 | 408034 |
| Height standing | *DOT1L* | 1.20E-06 | 5 | 408034 |
| Height standing | *SETD2* | 1.30E-06 | 19 | 408034 |
| Height standing | *USP37* | 1.40E-06 | 8 | 408034 |
| Height standing | *BCKDHA* | 1.91E-06 | 2 | 408034 |
| Height standing | *HR* | 1.92E-06 | 21 | 408034 |
| Height standing | *PAM* | 1.98E-06 | 5 | 408034 |
| Height standing | *PLG* | 2.04E-06 | 11 | 408034 |
| Height standing | *LLGL1* | 2.30E-06 | 9 | 408034 |
| Height standing | *ZNF205* | 2.42E-06 | 4 | 408034 |
| Body mass index | *GPR151* | 1.74E-09 | 6 | 407605 |
| Body mass index | *SYPL2* | 6.43E-09 | 4 | 407605 |
| Body mass index | *TRAPPC4* | 2.87E-07 | 2 | 407605 |
| Body mass index | *IQSEC1* | 7.02E-07 | 12 | 407605 |
| Body mass index | *HCRTR2* | 1.50E-06 | 4 | 407605 |
| Body mass index | *FRMD5* | 1.92E-06 | 4 | 407605 |
| Weight | *ZFAT* | 1.80E-11 | 13 | 401786 |
| Weight | *C16orf70* | 7.28E-11 | 2 | 401786 |
| Weight | *STC2* | 1.69E-09 | 4 | 401786 |
| Weight | *GPR151* | 1.73E-09 | 6 | 401786 |
| Weight | *ANO1* | 3.10E-08 | 4 | 401786 |
| Weight | *SYPL2* | 6.22E-08 | 4 | 401786 |
| Weight | *FRMD5* | 6.24E-08 | 4 | 401786 |
| Weight | *NUBP2* | 1.70E-07 | 9 | 401786 |
| Weight | *SCMH1* | 1.87E-07 | 8 | 401786 |
| Weight | *TRAPPC4* | 1.16E-06 | 2 | 401786 |

| | | | | |
|---|---|---|---|---|
| Weight | *HTRA1* | 2.00E-06 | 4 | 401786 |
| Weight | *SPSB3* | 2.32E-06 | 4 | 401786 |
| Whole body water mass | *STC2* | 8.55E-24 | 4 | 401782 |
| Whole body water mass | *NUBP2* | 1.52E-19 | 9 | 401782 |
| Whole body water mass | *ZFAT* | 1.83E-18 | 13 | 401782 |
| Whole body water mass | *SCMH1* | 8.44E-18 | 8 | 401782 |
| Whole body water mass | *SPSB3* | 2.40E-17 | 4 | 401782 |
| Whole body water mass | *HTRA1* | 2.11E-12 | 4 | 401782 |
| Whole body water mass | *ANO1* | 4.31E-12 | 4 | 401782 |
| Whole body water mass | *PHC3* | 1.32E-11 | 7 | 401782 |
| Whole body water mass | *C16orf70* | 3.33E-11 | 2 | 401782 |
| Whole body water mass | *ATAD2* | 2.82E-09 | 6 | 401782 |
| Whole body water mass | *GRM4* | 6.47E-09 | 2 | 401782 |
| Whole body water mass | *ESR1* | 1.15E-07 | 9 | 401782 |
| Whole body water mass | *ZMYM6* | 1.75E-07 | 11 | 401782 |
| Whole body water mass | *TMEM150B* | 2.96E-07 | 3 | 401782 |
| Whole body water mass | *FRMD5* | 2.98E-07 | 4 | 401782 |
| Whole body water mass | *LCOR* | 3.88E-07 | 11 | 401782 |
| Whole body water mass | *PLEKHJ1* | 9.18E-07 | 5 | 401782 |
| Whole body water mass | *GLI3* | 1.09E-06 | 8 | 401782 |
| Whole body water mass | *ACAN* | 2.12E-06 | 33 | 401782 |
| Basal metabolic rate | *STC2* | 1.50E-20 | 4 | 401771 |
| Basal metabolic rate | *ZFAT* | 2.39E-17 | 13 | 401771 |
| Basal metabolic rate | *NUBP2* | 4.62E-17 | 9 | 401771 |
| Basal metabolic rate | *SCMH1* | 3.15E-15 | 8 | 401771 |
| Basal metabolic rate | *SPSB3* | 5.37E-15 | 4 | 401771 |
| Basal metabolic rate | *C16orf70* | 9.08E-12 | 2 | 401771 |
| Basal metabolic rate | *HTRA1* | 4.87E-11 | 4 | 401771 |
| Basal metabolic rate | *ANO1* | 5.53E-11 | 4 | 401771 |
| Basal metabolic rate | *PHC3* | 2.11E-10 | 7 | 401771 |
| Basal metabolic rate | *GRM4* | 9.53E-10 | 2 | 401771 |
| Basal metabolic rate | *FRMD5* | 1.35E-07 | 4 | 401771 |
| Basal metabolic rate | *ATAD2* | 2.27E-07 | 6 | 401771 |
| Basal metabolic rate | *LCOR* | 7.41E-07 | 11 | 401771 |
| Basal metabolic rate | *ZMYM6* | 8.04E-07 | 11 | 401771 |
| Basal metabolic rate | *TMEM150B* | 1.10E-06 | 3 | 401771 |
| Basal metabolic rate | *ESR1* | 1.14E-06 | 9 | 401771 |
| Basal metabolic rate | *GPR151* | 1.97E-06 | 6 | 401771 |
| Whole body fat free mass | *STC2* | 5.52E-24 | 4 | 401747 |
| Whole body fat free mass | *NUBP2* | 1.70E-19 | 9 | 401747 |
| Whole body fat free mass | *ZFAT* | 5.30E-19 | 13 | 401747 |
| Whole body fat free mass | *SCMH1* | 2.90E-18 | 8 | 401747 |
| Whole body fat free mass | *SPSB3* | 2.51E-17 | 4 | 401747 |
| Whole body fat free mass | *HTRA1* | 1.32E-12 | 4 | 401747 |
| Whole body fat free mass | *ANO1* | 6.36E-12 | 4 | 401747 |
| Whole body fat free mass | *PHC3* | 7.42E-12 | 7 | 401747 |
| Whole body fat free mass | *C16orf70* | 1.18E-10 | 2 | 401747 |
| Whole body fat free mass | *GRM4* | 6.51E-09 | 2 | 401747 |
| Whole body fat free mass | *ATAD2* | 6.80E-09 | 6 | 401747 |
| Whole body fat free mass | *ESR1* | 7.69E-08 | 9 | 401747 |
| Whole body fat free mass | *ZMYM6* | 2.27E-07 | 11 | 401747 |
| Whole body fat free mass | *TMEM150B* | 5.78E-07 | 3 | 401747 |
| Whole body fat free mass | *FRMD5* | 5.88E-07 | 4 | 401747 |
| Whole body fat free mass | *LCOR* | 1.06E-06 | 11 | 401747 |
| Whole body fat free mass | *ACAN* | 1.31E-06 | 33 | 401747 |
| Whole body fat free mass | *PLEKHJ1* | 1.72E-06 | 5 | 401747 |
| Whole body fat free mass | *GLI3* | 1.94E-06 | 8 | 401747 |
| Whole body fat free mass | *PAM* | 2.01E-06 | 5 | 401747 |

| | | | | |
|---|---|---|---|---|
| Impedance of whole body | *CYR61* | 3.81E-18 | 2 | 401746 |
| Impedance of whole body | *POR* | 2.90E-12 | 7 | 401746 |
| Impedance of whole body | *ADAMTS3* | 7.23E-12 | 8 | 401746 |
| Impedance of whole body | *ANO1* | 4.91E-11 | 4 | 401746 |
| Impedance of whole body | *STC2* | 1.61E-09 | 4 | 401746 |
| Impedance of whole body | *ECM2* | 1.95E-08 | 10 | 401746 |
| Impedance of whole body | *ZNF469* | 3.93E-08 | 53 | 401746 |
| Impedance of whole body | *NUBP2* | 3.02E-07 | 9 | 401746 |
| Impedance of whole body | *FBN2* | 3.59E-07 | 16 | 401746 |
| Impedance of whole body | *FAM198A* | 1.08E-06 | 7 | 401746 |
| Impedance of whole body | *ZMYM6* | 1.18E-06 | 11 | 401746 |
| Impedance of whole body | *SNED1* | 1.99E-06 | 9 | 401746 |
| Body fat percentage | *CYR61* | 4.99E-12 | 2 | 401556 |
| Body fat percentage | *GPR151* | 2.85E-10 | 6 | 401556 |
| Body fat percentage | *SYPL2* | 2.81E-07 | 4 | 401556 |
| Body fat percentage | *C10orf35* | 3.64E-07 | 2 | 401556 |
| Days per week moderate phys activity 10min | *RAD51AP1* | 3.87E-07 | 7 | 389204 |
| Blood pressure diastolic automated mean | *DBH* | 3.08E-14 | 12 | 385365 |
| Blood pressure diastolic automated mean | *SLC9A3R2* | 2.90E-11 | 5 | 385365 |
| Blood pressure diastolic automated mean | *ARID1B* | 1.08E-06 | 7 | 385365 |
| Pulse rate automated mean | *TBX5* | 9.69E-35 | 4 | 385365 |
| Pulse rate automated mean | *MYH6* | 3.61E-15 | 14 | 385365 |
| Pulse rate automated mean | *TTN* | 3.18E-10 | 368 | 385365 |
| Pulse rate automated mean | *KIF1C* | 4.78E-10 | 12 | 385365 |
| Pulse rate automated mean | *ARHGEF40* | 7.02E-08 | 7 | 385365 |
| Pulse rate automated mean | *FNIP1* | 3.58E-07 | 8 | 385365 |
| Pulse rate automated mean | *DBH* | 1.74E-06 | 12 | 385365 |
| Blood pressure systolic automated mean | *SLC9A3R2* | 2.36E-12 | 5 | 385362 |
| Blood pressure systolic automated mean | *ZFAT* | 7.06E-12 | 13 | 385362 |
| Blood pressure systolic automated mean | *DBH* | 2.85E-10 | 12 | 385362 |
| Blood pressure systolic automated mean | *RRAS* | 1.33E-07 | 2 | 385362 |
| Blood pressure systolic automated mean | *NOX4* | 1.91E-07 | 4 | 385362 |
| Blood pressure systolic automated mean | *TBX5* | 2.87E-07 | 4 | 385362 |
| Blood pressure systolic automated mean | *COL21A1* | 8.05E-07 | 14 | 385362 |

**B.**

| Phenotype | Phecode | Gene | P-value | Number of variants | Number of cases | Number of controls |
|---|---|---|---|---|---|---|
| Cholelithiasis and cholecystitis | 574 | ABCG5 | 2.31E-13 | 13 | 16225 | 391307 |
| Cholelithiasis and cholecystitis | 574 | ABCG8 | 4.47E-10 | 12 | 16225 | 391307 |
| Cholelithiasis and cholecystitis | 574 | ABCB4 | 6.86E-07 | 9 | 16225 | 391307 |
| Sebaceous cyst | 706.2 | GORASP1 | 1.26E-18 | 8 | 8876 | 399255 |
| Diseases of hair and hair follicles | 704 | GORASP1 | 4.36E-16 | 8 | 5344 | 402357 |
| Glaucoma | 365 | MYOC | 1.24E-06 | 6 | 4462 | 397761 |
| Pulmonary heart disease | 415 | CREB3L1 | 9.59E-10 | 6 | 4257 | 402375 |
| Ankylosing spondylitis | 715.2 | SLC44A4 | 1.23E-15 | 8 | 620 | 365085 |
| Ankylosing spondylitis | 715.2 | PSMB9 | 5.83E-10 | 3 | 620 | 365085 |
| Ankylosing spondylitis | 715.2 | IER3 | 1.86E-09 | 4 | 620 | 365085 |

**Supplementary Table 5.** Exome-wide significant genes with p-values ≤ $2.5 \times 10^{-6}$ identified by SAIGE-GENE but not identified by SAIGE in the UK Biobank for 53 quantitative traits

.

| | | | | | Most significant variant in the locus (+/- 500kb of the start and end positions of the gene) | | | | | | | |
|---|---|---|---|---|---|---|---|---|---|---|---|---|
| **Phenotype** | **Gene** | **Number of Variants** | **Sample Size** | **P-value (SAKT-O)** | **chr:pos:ref:alt** | **Allele frequency** | **BETA** | **SE** | **P-value** | **In gene-based tests (0=no, 1=yes)** | **function** | **Gene** |
| Waist_circumference | *C16orf70* | 2 | 408227 | 1.94E-06 | 16:66995927:G:A | 0.00717746 | 0.07 | 0.01 | 5.57E-07 | 0 | intronic | *CES3* |
| Hip_circumference | *GPR151* | 6 | 408182 | 6.08E-07 | 5:146212195:C:G | 3.14E-05 | 1.64 | 0.34 | 1.12E-06 | 0 | intronic | *PPP2R2B* |
| Hip_circumference | *C16orf70* | 2 | 408182 | 1.14E-06 | 16:66995927:G:A | 0.00718015 | 0.07 | 0.02 | 7.05E-07 | 0 | intronic | *CES3* |
| Hip_circumference | *TRAPPC4* | 2 | 408182 | 5.83E-07 | 11:118890910:T:C | 0.00017785 | -0.52 | 0.10 | 6.94E-07 | 1 | nonsynonymous | *TRAPPC4* |
| Waist_hip_ratio | *GPR151* | 6 | 408144 | 3.00E-08 | 5:145895394:G:A | 0.00834999 | -0.05 | 0.01 | 8.08E-08 | 1 | stopgain | *GPR151* |
| Height_standing | *PFDN2* | 2 | 408034 | 1.41E-07 | 1:161071861:C:T | 0.00044532 | -0.25 | 0.05 | 8.46E-08 | 1 | nonsynonymous | *PFDN2* |
| Body_mass_index | *FRMD5* | 4 | 407605 | 1.92E-06 | 15:44167588:A:G | 0.04908352 | -0.03 | 0.01 | 1.09E-06 | 0 | intronic | *FRMD5* |
| Body_mass_index | *IQSEC1* | 12 | 407605 | 7.02E-07 | 3:12943013:T:G | 0.00655782 | 0.08 | 0.02 | 6.91E-08 | 1 | nonsynonymous | *IQSEC1* |
| Body_mass_index | *HCRTR2* | 4 | 407605 | 1.50E-06 | 6:54854851:A:C | 0.0145852 | 0.05 | 0.01 | 4.09E-07 | 0 | intergenic | *FAM83B(dist=44954), HCRTR2(dist=116407)* |
| Weight | *FRMD5* | 4 | 401786 | 6.24E-08 | 15:44147215:T:G | 0.00644606 | 0.07 | 0.01 | 1.65E-07 | 0 | intronic | *WDR76* |
| Weight | *HTRA1* | 4 | 401786 | 2.00E-06 | 10:124193181:T:G | 0.47045591 | 0.01 | 0.00 | 1.16E-07 | 0 | intergenic | *PLEKHA1(dist=1310), ARMS2(dist=20998)* |
| Weight | *TRAPPC4* | 2 | 401786 | 1.16E-06 | 11:118602242:C:T | 0.00013775 | -0.63 | 0.12 | 1.60E-07 | 0 | intergenic | *TREH(dist=51861), DDX6(dist=16231)* |
| Whole_body_water_mass | *ZMYM6* | 11 | 401782 | 1.75E-07 | 1:35298496:C:T | 5.12E-05 | 1.18 | 0.23 | 1.59E-07 | 0 | intergenic | *GJA4(dist=37148), SMIM12(dist=17467)* |
| Whole_body_water_mass | *FRMD5* | 4 | 401782 | 2.98E-07 | 15:44564692:A:G | 0.025069 | -0.03 | 0.01 | 6.33E-08 | 0 | intergenic | *FRMD5(dist=77200), CASC4(dist=16217)* |
| Whole_body_water_mass | *GLI3* | 8 | 401782 | 1.09E-06 | 7:41650943:C:T | 0.00041366 | -0.21 | 0.04 | 1.22E-06 | 0 | intergenic | *LINC01449(dist=477844), INHBA(dist=73769)* |
| Basal_metabolic_rate | *GPR151* | 6 | 401771 | 1.97E-06 | 5:146212195:C:G | 3.13E-05 | 1.09 | 0.23 | 1.74E-06 | 0 | intronic | *PPP2R2B* |
| Basal_metabolic_rate | *ZMYM6* | 11 | 401771 | 8.04E-07 | 1:35298496:C:T | 5.12E-05 | 1.20 | 0.24 | 3.81E-07 | 0 | intergenic | *GJA4(dist=37148), SMIM12(dist=17467)* |
| Whole_body_fat_free_mass | *ZMYM6* | 11 | 401747 | 2.27E-07 | 1:35298496:C:T | 5.12E-05 | 1.19 | 0.23 | 1.42E-07 | 0 | intergenic | *GJA4(dist=37148), SMIM12(dist=17467)* |
| Whole_body_fat_free_mass | *FRMD5* | 4 | 401747 | 5.88E-07 | 15:44564692:A:G | 0.02507002 | -0.03 | 0.01 | 7.16E-08 | 0 | intergenic | *FRMD5(dist=77200), CASC4(dist=16217)* |
| Whole_body_fat_free_mass | *GLI3* | 8 | 401747 | 1.94E-06 | 7:41650943:C:T | 0.00041369 | -0.21 | 0.04 | 1.34E-06 | 0 | intergenic | *LINC01449(dist=477844), INHBA(dist=73769)* |
| Body_fat_percentage | *C10orf35* | 2 | 401556 | 3.64E-07 | 10:71391560:A:G | 0.00316523 | 0.10 | 0.02 | 2.26E-07 | 1 | nonsynonymous | *C10orf35* |
| Days_per_week_moderate_phys_activity_10min | *RAD51AP1* | 7 | 389204 | 3.87E-07 | 12:4657293:A:G | 0.00656065 | 0.07 | 0.01 | 4.76E-07 | 1 | nonsynonymous | *RAD51AP1* |
| Blood_pressure_diastolic_automated_mean | *ARID1B* | 7 | 385365 | 1.08E-06 | 6:157525120:A:G | 0.00124304 | -0.19 | 0.04 | 9.01E-07 | 1 | nonsynonymous | *ARID1B* |

| | | | | | | | | | | | | |
|---|---|---|---|---|---|---|---|---|---|---|---|---|
| Pulse_rate_automated_mean | *DBH* | 12 | 385365 | 1.74E-06 | 9:136149399:G:A | 0.18699674 | -0.02 | 0.00 | 3.46E-06 | 0 | intronic | *ABO* |
| Blood_pressure_systolic_ automated_mean | *TBX5* | 4 | 385362 | 2.87E-07 | 12:114837349:C:A | 0.00492783 | -0.09 | 0.02 | 2.91E-07 | 1 | nonsynonymous | *TBX5* |

**Supplementary Table 6.** Exome-wide significant genes with p-values ≤ $2.5x10^{-6}$ identified by SAIGE-GENE and remained significant after conditioning on the most significant variant, given that the most significant variant is a common variant with MAF > 1% or a less frequent non-coding variant that is not included in the gene-based tests for A. 53 quantitative traits B. 6 binary traits.

A.

| | | | | | | **Most significant variant in the locus (+/- 500kb of the start and end positions of the gene)** | | | | | | |
|---|---|---|---|---|---|---|---|---|---|---|---|---|
| **Phenotype** | **Gene** | **Number of Variants** | **Sample Size** | **P-value (SKAT-O)** | **P-value (SKAT-O) conditional** | **chr:pos:ref:alt (GRCh37/hg19)** | **Allele frequency** | **BETA** | **SE** | **P-value** | **function** | **Gene** |
| Hip_circumference | GPR151 | 6 | 408,182 | 6.08E-07 | 6.13E-07 | 5:146212195:C:G | 3.14E-05 | 1.636 | 0.336 | 1.12E-06 | intronic | PPP2R2B |
| Hip_circumference | ANO1 | 4 | 408,182 | 5.98E-07 | 7.03E-07 | 11:69482091:C:A | 0.646 | 0.015 | 0.003 | 1.77E-08 | UTR3 | ORAOV1 (NM_153451:c.*196G>T) |
| Waist_hip_ratio | SLC5A3 | 4 | 408,144 | 1.33E-07 | 6.23E-08 | 21:35593827:G:A | 0.131 | 0.017 | 0.003 | 1.33E-09 | intergenic | LINC00310(dist=31607) ,KCNE2(dist=142496) |
| Height_standing | C11orf57 | 2 | 408,034 | 5.71E-07 | 5.72E-07 | 11:111889209:T:C | 2.71E-05 | -1.410 | 0.226 | 4.42E-10 | intronic | DIXDC1 |
| Height_standing | CADM1 | 2 | 408,034 | 1.35E-07 | 1.36E-07 | 11:115841824:A:C | 6.99E-05 | 0.870 | 0.141 | 6.91E-10 | intergenic | LINC00900(dist=210906) ,LOC101929011(dist=668315) |
| Height_standing | PTH1R | 3 | 408,034 | 1.40E-09 | 7.88E-11 | 3:46933960:C:G | 0.581 | -0.016 | 0.002 | 6.36E-18 | intronic | PTH1R |
| Height_standing | SIX6 | 3 | 408,034 | 1.16E-06 | 3.30E-09 | 14:61072875:T:C | 0.613 | -0.022 | 0.002 | 3.07E-30 | intergenic | SIX6(dist=94350) ,SALRNA1(dist=33059) |
| Height_standing | TBX3 | 2 | 408,034 | 4.59E-09 | 7.75E-08 | 12:115108136:T:C | 0.257 | 0.016 | 0.002 | 1.75E-14 | UTR3 | TBX3 |
| Height_standing | NOX4 | 4 | 408,034 | 4.09E-09 | 2.61E-07 | 11:89216425:T:A | 0.796 | 0.015 | 0.002 | 9.65E-11 | intronic | NOX4 |
| Height_standing | S1PR5 | 5 | 408,034 | 1.44E-10 | 3.18E-10 | 19:10754905:G:T | 0.664 | -0.022 | 0.002 | 8.66E-29 | UTR3 | SLC44A2 |
| Height_standing | MTMR11 | 3 | 408,034 | 1.94E-15 | 1.82E-20 | 1:149906413:T:C | 0.409 | 0.035 | 0.002 | 7.59E-76 | nonsynonymous | MTMR11 |
| Height_standing | SERPINE2 | 3 | 408,034 | 2.80E-07 | 5.64E-07 | 2:225357325:T:C | 0.004 | 0.224 | 0.016 | 5.68E-47 | intronic | CUL3 |
| Height_standing | ST3GAL4 | 5 | 408,034 | 5.37E-11 | 2.15E-10 | 11:125825224:G:T | 0.582 | 0.015 | 0.002 | 1.81E-14 | intergenic | DDX25 |
| Height_standing | ABCB6 | 11 | 408,034 | 4.74E-08 | 1.38E-08 | 2:220004944:T:C | 0.017 | -0.139 | 0.008 | 2.50E-76 | intronic | NHEJ1 |
| Height_standing | TMEM150B | 3 | 408,034 | 2.88E-07 | 1.36E-07 | 19:55993436:G:T | 0.024 | -0.101 | 0.006 | 2.83E-62 | nonsynonymous | ZNF628 |
| Height_standing | ZFAT | 13 | 408,034 | 3.61E-32 | 7.36E-32 | 8:135598132:C:G | 0.368 | -0.023 | 0.002 | 2.53E-31 | intronic | ZFAT |
| Height_standing | ELN | 8 | 408,034 | 6.67E-08 | 2.92E-07 | 7:73474825:G:C | 0.099 | 0.032 | 0.003 | 2.72E-24 | nonsynonymous | ELN |
| Height_standing | AMOTL1 | 6 | 408,034 | 2.85E-07 | 2.87E-07 | 11:94547499:A:G | 3.18E-03 | 0.099 | 0.017 | 3.21E-09 | intronic | AMOTL1 |
| Height_standing | COL11A1 | 8 | 408,034 | 2.91E-07 | 2.90E-07 | 1:103419238:A:G | 0.700 | -0.021 | 0.002 | 9.33E-24 | intronic | COL11A1 |
| Height_standing | PTCH1 | 15 | 408,034 | 2.65E-10 | 1.48E-09 | 9:98368761:T:C | 0.258 | -0.044 | 0.002 | 7.44E-91 | intergenic | PTCH1(dist=89514), LINC00476(dist=199609) |
| Height_standing | GRM4 | 2 | 408,034 | 7.23E-16 | 2.15E-10 | 6:34199092:C:T | 0.911 | -0.078 | 0.003 | 2.35E-125 | intergenic | GRM4(dist=75693), HMGA1(dist=5485) |
| Height_standing | HAPLN3 | 6 | 408,034 | 1.25E-11 | 1.07E-11 | 15:89397827:A:G | 0.030 | -0.097 | 0.006 | 5.72E-69 | intronic | ACAN |
| Height_standing | VASN | 7 | 408,034 | 5.25E-07 | 5.16E-07 | 16:4019350:C:T | 0.821 | 0.021 | 0.002 | 2.35E-18 | intronic | ADCY9 |
| Height_standing | PRSS56 | 3 | 408,034 | 1.68E-07 | 1.50E-07 | 2:232948266:A:G | 0.916 | 0.048 | 0.003 | 1.64E-45 | intronic | DIS3L2 |
| Height_standing | TXLNA | 3 | 408,034 | 5.66E-07 | 5.84E-07 | 1:32356815:C:T | 0.061 | 0.029 | 0.004 | 7.70E-14 | intergenic | SPOCD1(dist=75163), PTP4A2(dist=15207) |
| Height_standing | PHC3 | 7 | 408,034 | 3.24E-07 | 3.54E-07 | 3:169701455:A:G | 3.29E-06 | -7.403 | 0.729 | 3.06E-24 | intronic | SEC62 |
| Height_standing | CLEC3A | 3 | 408,034 | 9.08E-10 | 8.21E-10 | 16:78064682:G:A | 1.87E-04 | 0.558 | 0.078 | 6.46E-13 | nonsynonymous | CLEC3A |
| Height_standing | ESR1 | 9 | 408,034 | 7.28E-10 | 3.49E-07 | 6:152125444:T:C | 0.661 | 0.020 | 0.002 | 5.95E-24 | intronic | ESR1 |
| Height_standing | SAMD4A | 7 | 408,034 | 4.77E-09 | 1.15E-08 | 14:55238871:C:T | 0.321 | 0.016 | 0.002 | 3.64E-16 | intronic | SAMD4A |

| | | | | | | | | | | | | |
|---|---|---|---|---|---|---|---|---|---|---|---|---|
| Height_standing | LRRC8A | 2 | 408,034 | 2.87E-08 | 2.87E-08 | 9:131245464:G:A | 2.42E-03 | -0.165 | 0.021 | 1.68E-15 | ncRNA_intronic | MIR1268A |
| Height_standing | CPPED1 | 7 | 408,034 | 1.40E-07 | 1.32E-07 | 16:12399629:T:A | 1.01E-03 | 0.211 | 0.035 | 1.82E-09 | intronic | SNX29 |
| Height_standing | FER | 5 | 408,034 | 6.68E-12 | 1.12E-10 | 5:108171483:G:A | 0.083 | 0.044 | 0.003 | 2.40E-38 | synonymous | FER |
| Height_standing | LPP | 8 | 408,034 | 2.11E-08 | 8.72E-08 | 3:187443314:G:A | 0.071 | 0.036 | 0.004 | 1.48E-23 | synonymous | BCL6 |
| Height_standing | TRAPPC13 | 2 | 408,034 | 1.04E-10 | 1.15E-10 | 5:64766798:G:A | 2.08E-03 | -0.216 | 0.023 | 1.36E-21 | nonsynonymous | ADAMTS6 |
| Height_standing | CTU2 | 12 | 408,034 | 2.09E-08 | 1.92E-08 | 16:88782050:G:A | 0.010 | -0.086 | 0.010 | 9.03E-19 | nonsynonymous | PIEZO1 |
| Height_standing | PIEZO1 | 55 | 408,034 | 6.63E-08 | 2.21E-08 | 16:88782050:G:A | 0.010 | -0.086 | 0.010 | 9.03E-19 | nonsynonymous | PIEZO1 |
| Height_standing | PROP1 | 4 | 408,034 | 1.63E-07 | 2.89E-07 | 5:176980904:G:A | 0.296 | 0.013 | 0.002 | 8.92E-10 | intronic | FAM193B |
| Height_standing | FNDC3B | 8 | 408,034 | 1.39E-12 | 6.82E-14 | 3:172188000:A:G | 0.493 | -0.025 | 0.002 | 4.54E-40 | intergenic | GHSR(dist=21754), TNFSF10(dist=35298) |
| Height_standing | TARS2 | 3 | 408,034 | 4.22E-07 | 3.80E-09 | 1:149995265:G:A | 0.398 | 0.035 | 0.002 | 7.13E-69 | intergenic | OTUD7B(dist=12579), VPS45(dist=44085) |
| Height_standing | GYG1 | 5 | 408,034 | 1.78E-07 | 1.47E-07 | 3:148856741:A:C | 0.064 | -0.022 | 0.004 | 5.18E-09 | intronic | HPS3 |
| Height_standing | NUBP2 | 9 | 408,034 | 2.01E-22 | 2.51E-23 | 16:2160503:T:G | 0.169 | -0.030 | 0.003 | 9.96E-34 | synonymous | PKD1 |
| Height_standing | SPSB3 | 4 | 408,034 | 1.10E-22 | 1.19E-23 | 16:2160503:T:G | 0.169 | -0.030 | 0.003 | 9.96E-34 | synonymous | PKD1 |
| Height_standing | PKD1 | 43 | 408,034 | 6.40E-12 | 1.96E-08 | 16:2160503:T:G | 0.169 | -0.030 | 0.003 | 9.96E-34 | synonymous | PKD1 |
| Height_standing | C16orf70 | 2 | 408,034 | 7.45E-12 | 4.15E-11 | 16:67329937:T:C | 0.074 | -0.032 | 0.004 | 3.83E-19 | intronic | KCTD19 |
| Height_standing | ATAD5 | 8 | 408,034 | 4.74E-10 | 1.79E-07 | 17:29211667:G:A | 0.270 | -0.042 | 0.002 | 4.20E-87 | intronic | ATAD5 |
| Height_standing | CYR61 | 2 | 408,034 | 4.40E-09 | 7.38E-09 | 1:85997286:G:C | 0.168 | 0.019 | 0.003 | 6.28E-13 | intronic | DDAH1 |
| Height_standing | STC2 | 4 | 408,034 | 1.10E-24 | 4.46E-25 | 5:172993684:C:G | 0.631 | -0.025 | 0.002 | 1.87E-37 | intergenic | MIR8056(dist=219145), LOC285593(dist=12953) |
| Height_standing | PTK7 | 6 | 408,034 | 2.95E-07 | 6.24E-08 | 6:43002721:A:G | 0.139 | -0.018 | 0.003 | 1.20E-11 | intergenic | RRP36(dist=5384), CUL7(dist=2634) |
| Height_standing | ZCCHC6 | 7 | 408,034 | 1.42E-08 | 1.49E-07 | 9:89089476:C:T | 0.503 | 0.016 | 0.002 | 3.03E-18 | intergenic | ZCCHC6(dist=120074), GAS1(dist=469801) |
| Height_standing | NPR3 | 3 | 408,034 | 3.60E-25 | 4.14E-30 | 5:32711633:C:A | 0.191 | -0.041 | 0.002 | 3.15E-61 | UTR5 | NPR3 |
| Body_mass_index | TRAPPC4 | 2 | 407,605 | 2.87E-07 | 2.85E-07 | 11:118396331:A:T | 0.023 | -0.048 | 0.009 | 2.38E-08 | ncRNA_intronic | LOC101929089 |
| Weight | SCMH1 | 8 | 401,786 | 1.87E-07 | 3.69E-07 | 1:41570459:G:A | 0.776 | -0.022 | 0.003 | 1.42E-15 | intronic | SCMH1 |
| Weight | NUBP2 | 9 | 401,786 | 1.70E-07 | 7.64E-08 | 16:2160503:T:G | 0.169 | -0.027 | 0.003 | 4.02E-19 | synonymous | PKD1 |
| Weight | ZFAT | 13 | 401,786 | 1.80E-11 | 3.38E-11 | 8:135612745:A:G | 0.406 | -0.018 | 0.002 | 5.67E-15 | synonymous | ZFAT |
| Whole_body_water_mass | ZMYM6 | 11 | 401,782 | 1.75E-07 | 1.78E-07 | 1:35298496:C:T | 5.12E-05 | 1.184 | 0.226 | 1.59E-07 | intergenic | GJA4(dist=37148), SMIM12(dist=17467) |
| Whole_body_water_mass | FRMD5 | 4 | 401,782 | 2.98E-07 | 5.70E-07 | 15:44564692:A:G | 0.025 | -0.029 | 0.005 | 6.33E-08 | intergenic | FRMD5(dist=77200), CASC4(dist=16217) |
| Whole_body_water_mass | LCOR | 11 | 401,782 | 3.88E-07 | 5.08E-07 | 10:98657257:C:G | 0.014 | -0.045 | 0.007 | 6.72E-11 | intronic | LCOR |
| Whole_body_water_mass | TMEM150B | 3 | 401,782 | 2.96E-07 | 2.83E-07 | 19:55993436:G:T | 0.024 | -0.040 | 0.005 | 2.33E-14 | nonsynonymous | ZNF628 |
| Whole_body_water_mass | ESR1 | 9 | 401,782 | 1.15E-07 | 6.31E-07 | 6:152170247:G:A | 0.455 | 0.014 | 0.002 | 2.55E-17 | intronic | ESR1 |
| Whole_body_water_mass | SCMH1 | 8 | 401,782 | 8.44E-18 | 1.58E-17 | 1:41570459:G:A | 0.776 | -0.022 | 0.002 | 2.47E-29 | intronic | SCMH1 |
| Whole_body_water_mass | NUBP2 | 9 | 401,782 | 1.52E-19 | 2.69E-20 | 16:2160503:T:G | 0.169 | -0.024 | 0.002 | 6.19E-29 | synonymous | PKD1 |
| Whole_body_water_mass | SPSB3 | 4 | 401,782 | 2.40E-17 | 3.72E-18 | 16:2160503:T:G | 0.169 | -0.024 | 0.002 | 6.19E-29 | synonymous | PKD1 |
| Whole_body_water_mass | ZFAT | 13 | 401,782 | 1.83E-18 | 1.96E-17 | 8:135612745:A:G | 0.406 | -0.016 | 0.002 | 4.79E-22 | synonymous | ZFAT |

| | | | | | | | | | | | | |
|---|---|---|---|---|---|---|---|---|---|---|---|---|
| Basal_metabolic_rate | ZMYM6 | 11 | 401,771 | 8.04E-07 | 8.06E-07 | 1:35298496:C:T | 5.12E-05 | 1.198 | 0.236 | 3.81E-07 | intergenic | GJA4(dist=37148), SMIM12(dist=17467) |
| Basal_metabolic_rate | TMEM150B | 3 | 401,771 | 1.10E-06 | 5.76E-07 | 19:55993436:G:T | 0.024 | -0.039 | 0.006 | 1.66E-12 | nonsynonymous | ZNF628 |
| Basal_metabolic_rate | FRMD5 | 4 | 401,771 | 1.35E-07 | 2.81E-07 | 15:44028047:C:T | 0.024 | -0.030 | 0.006 | 4.77E-08 | downstream | CATSPER2P1(dist=99) |
| Basal_metabolic_rate | GRM4 | 2 | 401,771 | 9.53E-10 | 5.94E-07 | 6:34620153:T:A | 0.139 | 0.035 | 0.002 | 1.86E-45 | intronic | C6orf106 |
| Basal_metabolic_rate | SCMH1 | 8 | 401,771 | 3.15E-15 | 5.90E-15 | 1:41570459:G:A | 0.776 | -0.022 | 0.002 | 8.01E-27 | intronic | SCMH1 |
| Basal_metabolic_rate | SPSB3 | 4 | 401,771 | 5.37E-15 | 9.79E-16 | 16:2160503:T:G | 0.169 | -0.025 | 0.002 | 1.21E-27 | synonymous | PKD1 |
| Basal_metabolic_rate | NUBP2 | 9 | 401,771 | 4.62E-17 | 9.62E-18 | 16:2160503:T:G | 0.169 | -0.025 | 0.002 | 1.21E-27 | synonymous | PKD1 |
| Basal_metabolic_rate | ZFAT | 13 | 401,771 | 2.39E-17 | 1.46E-16 | 8:135612745:A:G | 0.406 | -0.016 | 0.002 | 3.07E-21 | synonymous | ZFAT |
| Whole_body_fat_free_mass | ZMYM6 | 11 | 401,747 | 2.27E-07 | 2.04E-07 | 1:35298496:C:T | 5.12E-05 | 1.187 | 0.226 | 1.42E-07 | intergenic | GJA4(dist=37148), SMIM12(dist=17467) |
| Whole_body_fat_free_mass | TMEM150B | 3 | 401,747 | 5.78E-07 | 2.88E-07 | 19:55993436:G:T | 0.024 | -0.041 | 0.005 | 1.40E-14 | nonsynonymous | ZNF628 |
| Whole_body_fat_free_mass | ESR1 | 9 | 401,747 | 7.69E-08 | 5.22E-07 | 6:152170247:G:A | 0.455 | 0.014 | 0.002 | 8.43E-17 | intronic | ESR1 |
| Whole_body_fat_free_mass | SCMH1 | 8 | 401,747 | 2.90E-18 | 5.34E-18 | 1:41570459:G:A | 0.776 | -0.022 | 0.002 | 2.00E-29 | intronic | SCMH1 |
| Whole_body_fat_free_mass | SPSB3 | 4 | 401,747 | 2.51E-17 | 3.82E-18 | 16:2160503:T:G | 0.169 | -0.025 | 0.002 | 1.96E-29 | synonymous | PKD1 |
| Whole_body_fat_free_mass | NUBP2 | 9 | 401,747 | 1.70E-19 | 2.96E-20 | 16:2160503:T:G | 0.169 | -0.025 | 0.002 | 1.96E-29 | synonymous | PKD1 |
| Whole_body_fat_free_mass | ZFAT | 13 | 401,747 | 5.30E-19 | 4.72E-18 | 8:135612745:A:G | 0.406 | -0.016 | 0.002 | 4.24E-22 | synonymous | ZFAT |
| Impedance_of_whole_body | ADAMTS3 | 8 | 401,746 | 7.23E-12 | 2.12E-11 | 4:73519842:C:T | 0.063 | 0.053 | 0.004 | 2.90E-39 | intergenic | ADAMTS3(dist=85326), COX18(dist=400571) |
| Impedance_of_whole_body | SNED1 | 9 | 401,746 | 1.99E-06 | 1.36E-07 | 2:242050740:A:G | 0.329 | 0.014 | 0.002 | 1.29E-11 | intronic | PASK |
| Impedance_of_whole_body | NUBP2 | 9 | 401,746 | 3.02E-07 | 1.59E-07 | 16:2163962:A:G | 0.098 | 0.025 | 0.003 | 1.92E-13 | intronic | PKD1 |
| Impedance_of_whole_body | FBN2 | 16 | 401,746 | 3.59E-07 | 9.05E-07 | 5:127367998:G:C | 0.248 | -0.032 | 0.002 | 1.53E-43 | ncRNA_intronic | LINC01184 |
| Impedance_of_whole_body | ZNF469 | 53 | 401,746 | 3.93E-08 | 1.73E-08 | 16:88321027:C:T | 0.041 | 0.044 | 0.005 | 2.78E-18 | intergenic | LINC02182(dist=92204), ZNF469(dist=172852) |
| Pulse_rate_automated_mean | ARHGEF40 | 7 | 385,365 | 7.02E-08 | 2.57E-10 | 14:21542766:A:G | 0.169 | 0.051 | 0.003 | 3.30E-52 | nonsynonymous | ARHGEF40 |
| Pulse_rate_automated_mean | MYH6 | 14 | 385,365 | 3.61E-15 | 2.56E-13 | 14:23861811:A:G | 0.370 | 0.072 | 0.003 | 1.04E-168 | nonsynonymous | MYH6 |
| Blood_pressure_diastolic_automated_mean | DBH | 12 | 385,365 | 3.08E-14 | 5.20E-15 | 9:136522274:C:T | 0.074 | -0.041 | 0.005 | 4.91E-18 | nonsynonymous | DBH |
| Blood_pressure_systolic_automated_mean | RRAS | 2 | 385,362 | 1.33E-07 | 2.79E-07 | 19:49639399:C:G | 0.155 | -0.020 | 0.003 | 3.68E-10 | intronic | PPFIA3 |
| Blood_pressure_systolic_automated_mean | SLC9A3R2 | 5 | 385,362 | 2.36E-12 | 2.80E-13 | 16:2089006:G:A | 0.159 | -0.024 | 0.003 | 5.15E-14 | UTR3 | SLC9A3R2 |
| Blood_pressure_systolic_automated_mean | DBH | 12 | 385,362 | 2.85E-10 | 1.05E-10 | 9:136522274:C:T | 0.074 | -0.030 | 0.005 | 1.47E-11 | nonsynonymous | DBH |

B.

| | | | | | | | | Most significant variant in the locus (+/- 500kb of the start and end positions of the gene) | | | | | | |
|---|---|---|---|---|---|---|---|---|---|---|---|---|---|---|
| **Phenotype** | **Phecode** | **Gene** | **Number of Variants** | **Number of cases** | **Number of controls** | **P-value (SKAT-O)** | **P-value (SKAT-O) conditional** | **chr:pos:ref:alt (GRCh37/hg19)** | **Allele frequency** | **Beta** | **SE** | **P-value** | **Function** | **Gene** |
| Cholelithiasis and cholecystitis | 574 | ABCB4 | 9 | 16225 | 391307 | 6.86E-07 | 1.46E-10 | 7:87105795:T:C | 0.139 | -0.148 | 0.017 | 3.29E-18 | upstream | ABCB4(dist=776) |
| Cholelithiasis and cholecystitis | 574 | ABCG5 | 13 | 16225 | 391307 | 2.31E-13 | 4.58E-11 | 2:44069772:G:A | 0.065 | 0.767 | 0.025 | 4.16E-201 | intronic | ABCG8 |
| Cholelithiasis and cholecystitis | 574 | ABCG8 | 12 | 16225 | 391307 | 4.47E-10 | 7.42E-11 | 2:44069772:G:A | 0.065 | 0.767 | 0.025 | 4.16E-201 | intronic | ABCG8 |
| Diseases of hair and hair follicles | 704 | GORASP1 | 8 | 5344 | 402357 | 4.36E-16 | 2.41E-11 | 3:38659248:G:C | 0.113 | 0.408 | 0.033 | 1.50E-35 | intronic | SCN5A |
| Ankylosing spondylitis | 715.2 | IER3 | 4 | 620 | 365085 | 1.86E-09 | 2.34E-08 | 6:31210279:C:T | 0.036 | 3.73 | 0.209 | 2.21E-71 | intergenic | HCG27(dist=38534), HLA-C(dist=26247) |
| Ankylosing spondylitis | 715.2 | PSMB9 | 3 | 620 | 365085 | 5.83E-10 | 1.36E-07 | 6:32582577:A:C | 0.340 | 0.61 | 0.062 | 1.10E-22 | intergenic | HLA-DRB1(dist=24964), HLA-DQA1(dist=22606) |

**Supplementary Table 7.** Empirical type I error rates for SAIGE-GENE, SAIGE-GENE-GCadj (GC adjusted SAIGE-GENE), EmmaX-SKAT[9,22] and SMMAT[21]. $h^2$: heritability.

| | | **500 families and 5000 independent samples** | | | | | | **1000 families and no independent samples** | | | | | |
|---|---|---|---|---|---|---|---|---|---|---|---|---|---|
| | | $h^2$**=0.2** | | | $h^2$**=0.4** | | | $h^2$**=0.2** | | | $h^2$**=0.4** | | |
| | **alpha** | **burden** | **skat** | **skato** | **burden** | **skat** | **skato** | **burden** | **skat** | **skato** | **burden** | **skat** | **skato** |
| **SAIGE-GENE** | **0.05** | 5.25E-02 | 5.62E-02 | 5.79E-02 | 5.49E-02 | 6.26E-02 | 6.32E-02 | 5.26E-02 | 5.62E-02 | 5.79E-02 | 5.48E-02 | 6.24E-02 | 6.30E-02 |
| | **0.0001** | 1.10E-04 | 1.30E-04 | 1.30E-04 | 1.40E-04 | 1.70E-04 | 1.80E-04 | 1.20E-04 | 1.30E-04 | 1.40E-04 | 1.40E-04 | 1.70E-04 | 1.80E-04 |
| | **2.50E-06** | 2.80E-06 | 4.20E-06 | 2.80E-06 | 3.50E-06 | 5.20E-06 | 3.30E-06 | 3.71E-06 | 5.22E-06 | 5.82E-06 | 3.81E-06 | 6.51E-06 | 5.61E-06 |
| **SAIGE-GENE -Gcadj** | **0.05** | 5.00E-02 | 5.00E-02 | 5.45E-02 | 5.01E-02 | 5.01E-02 | 5.10E-02 | 4.99E-02 | 4.99E-02 | 5.47E-02 | 4.99E-02 | 4.99E-02 | 5.10E-02 |
| | **0.0001** | 9.00E-05 | 9.00E-05 | 1.10E-04 | 1.00E-04 | 1.00E-04 | 8.00E-05 | 1.00E-04 | 1.00E-04 | 1.20E-04 | 1.00E-04 | 1.00E-04 | 8.00E-05 |
| | **2.50E-06** | 2.20E-06 | 2.20E-06 | 2.20E-06 | 1.90E-06 | 1.80E-06 | 1.30E-06 | 2.91E-06 | 2.91E-06 | 4.41E-06 | 2.51E-06 | 2.61E-06 | 2.51E-06 |
| **EmmaX-SKAT** | **0.05** | 5.16E-02 | 5.36E-02 | 5.58E-02 | 5.33E-02 | 5.75E-02 | 5.91E-02 | 5.18E-02 | 5.39E-02 | 5.60E-02 | 5.32E-02 | 5.76E-02 | 5.91E-02 |
| | **0.0001** | 1.10E-04 | 1.10E-04 | 1.20E-04 | 1.20E-04 | 1.30E-04 | 1.50E-04 | 1.10E-04 | 1.20E-04 | 1.30E-04 | 1.30E-04 | 1.40E-04 | 1.50E-04 |
| | **2.50E-06** | 2.36E-06 | 3.37E-06 | 2.13E-06 | 2.58E-06 | 3.03E-06 | 2.25E-06 | 3.71E-06 | 4.61E-06 | 5.40E-06 | 3.49E-06 | 5.06E-06 | 4.95E-06 |
| **SMMAT** | **0.05** | 5.17E-02 | 5.36E-02 | 5.41E-02 | 5.33E-02 | 5.74E-02 | 5.73E-02 | 5.18E-02 | 5.39E-02 | 5.43E-02 | 5.32E-02 | 5.76E-02 | 5.74E-02 |
| | **0.0001** | 1.10E-04 | 1.10E-04 | 1.40E-04 | 1.20E-04 | 1.30E-04 | 1.70E-04 | 1.10E-04 | 1.20E-04 | 1.50E-04 | 1.30E-04 | 1.40E-04 | 1.70E-04 |
| | **2.50E-06** | 2.50E-06 | 3.80E-06 | 2.90E-06 | 2.80E-06 | 3.40E-06 | 3.30E-06 | 3.47E-06 | 4.42E-06 | 6.00E-06 | 3.26E-06 | 4.63E-06 | 5.47E-06 |

**Supplementary Table 8.** Empirical type I error rates for SAIGE-GENE with the larger sample size of 1,000 families and 10,000 independent samples (total sample size N = 20,000). The heritability $h^2 = 0.2$.

| | alpha | Burden | SKAT | SKAT-O |
|---|---|---|---|---|
| SAIGE-GENE | **0.05** | 5.28E-02 | 5.68E-02 | 5.84E-02 |
| | **0.0001** | 1.00E-04 | 1.00E-04 | 1.00E-04 |
| | **2.50E-06** | 3.44E-06 | 4.33E-06 | 3.00E-06 |

**Supplementary Table 9.** Empirical type I error rates for SAIGE-GENE, EmmaX-SKAT[9,22] and SMMAT[21] for skewed distributed phenotypes with and without inverse normal transformation for 500 families and 5,000 independent samples (total sample size N = 10,000). The heritability $h^2 = 0.2$.

| | | with inverse normal transformation | | | without inverse normal transformation | | |
|---|---|---|---|---|---|---|---|
| | **alpha** | **Burden** | **SKAT** | **SKAT-O** | **Burden** | **SKAT** | **SKAT-O** |
| **SAIGE-GENE** | **0.05** | 5.19E-02 | 5.61E-02 | 5.75E-02 | 5.13E-02 | 5.85E-02 | 5.87E-02 |
| | **0.0001** | 1.25E-04 | 1.75E-04 | 1.81E-04 | 1.52E-04 | 3.50E-04 | 3.27E-04 |
| | **2.50E-06** | 4.44E-06 | 8.27E-06 | 6.65E-06 | 5.68E-06 | 2.86E-05 | 2.19E-05 |
| **EmmaX-SKAT** | **0.05** | 5.10E-02 | 5.37E-02 | 5.55E-02 | 5.09E-02 | 5.75E-02 | 5.79E-02 |
| | **0.0001** | 1.18E-04 | 1.55E-04 | 1.61E-04 | 1.45E-04 | 3.48E-04 | 3.30E-04 |
| | **2.50E-06** | 4.82E-06 | 6.55E-06 | 6.55E-06 | 6.92E-06 | 2.42E-05 | 2.29E-05 |
| **SMMAT** | **0.05** | 5.14E-02 | 5.41E-02 | 5.45E-02 | 5.09E-02 | 5.72E-02 | 5.59E-02 |
| | **0.0001** | 1.19E-04 | 1.62E-04 | 1.83E-04 | 1.42E-04 | 3.40E-04 | 3.40E-04 |
| | **2.50E-06** | 5.42E-06 | 8.34E-06 | 9.17E-06 | 6.79E-06 | 2.59E-05 | 2.59E-05 |

**Supplementary Table 10.** Empirical type I error rates for SAIGE-GENE and SMMAT[21] for skewed distributed phenotypes with the three-step phenotype transformation procedure for 500 families and 5,000 independent samples (total sample size N = 10,000). The heritability $h^2 = 0.2$.

| | alpha | Burden | SKAT | SKAT-O |
|---|---|---|---|---|
| **SAIGE-GENE** | **0.05** | 4.71E-02 | 4.42E-02 | 4.77E-02 |
| | **0.0001** | 8.53E-05 | 7.94E-05 | 9.32E-05 |
| | **2.50E-06** | 1.79E-06 | 2.53E-06 | 2.74E-06 |
| **SMMAT** | **0.05** | 5.03E-02 | 5.06E-02 | 5.16E-02 |
| | **0.0001** | 1.03E-04 | 1.05E-04 | 1.29E-04 |
| | **2.50E-06** | 1.71E-06 | 2.32E-06 | 3.54E-06 |

**Supplementary Table 11.** Empirical type I error rates for SAIGE-GENE in the presence of population stratification. Phenotypes were simulated based on real genotypes from the UK Biobank for randomly selected 5,000 samples with white British ancestry and 5,000 samples with European ancestry but not white British. The heritability $h^2 = 0.2$.

| alpha | Burden | SKAT | SKAT-O |
|---|---|---|---|
| **0.05** | 5.05E-02 | 5.09E-02 | 5.33E-02 |
| **0.0001** | 1.02E-04 | 1.06E-04 | 1.33E-04 |
| **2.50E-06** | 2.42E-06 | 2.42E-06 | 3.85E-06 |

**Supplementary Table 12.** Empirical type I error rates for SAIGE-GENE in the presence of non-negligible cryptic relatedness. Phenotypes were simulated based on real genotypes from the UK Biobank for randomly selected 10,000 sample with White British ancestry (5,000 are related with up to 3rd degree and 5,000 are unrelated). The heritability $h^2 = 0.2$.

| alpha | Burden | SKAT | SKAT-O |
|---|---|---|---|
| **0.05** | 5.07E-02 | 5.11E-02 | 5.35E-02 |
| **0.0001** | 1.06E-04 | 1.07E-04 | 1.45E-04 |
| **2.50E-06** | 3.04E-06 | 2.88E-06 | 3.80E-06 |

**Supplementary Table 13.** Empirical Type I error rates of SAIGE-GENE for binary traits with five different prevalence. Phenotypes were simulated for 500 families and 5,000 independent samples (total sample size N = 10,000). The liability scale heritability $h^2_{latent}$ =0.23. A. Unadjusted SAIGE-GENE; B. SAIGE-GENE with the robust adjustment to account for unbalanced case-control ratios. SKAT-O-GCadj and SKAT-GCadj are GC-adjusted SKAT-O and SKAT.

A. Unadjusted SAIGE-GENE without applying the robust adjustment.

| | Alpha | Prev=0.01 | Prev=0.05 | Prev=0.1 | Prev=0.2 | Prev=0.5 |
|---|---|---|---|---|---|---|
| **SKAT-O** | **0.05** | 7.27E-02 | 5.55E-02 | 5.38E-02 | 5.31E-02 | 5.32E-02 |
| | **0.0001** | 2.77E-03 | 4.57E-04 | 2.32E-04 | 1.43E-04 | 1.04E-04 |
| | **2.50E-06** | 5.38E-04 | 4.04E-05 | 1.49E-05 | 5.76E-06 | 1.70E-06 |
| **SKAT** | **0.05** | 8.67E-02 | 5.59E-02 | 5.21E-02 | 5.05E-02 | 5.03E-02 |
| | **0.0001** | 3.10E-03 | 4.74E-04 | 2.33E-04 | 1.30E-04 | 8.96E-05 |
| | **2.50E-06** | 6.34E-04 | 4.64E-05 | 1.58E-05 | 6.69E-06 | 1.70E-06 |
| **Burden** | **0.05** | 4.57E-02 | 4.94E-02 | 4.99E-02 | 5.03E-02 | 5.06E-02 |
| | **0.0001** | 5.69E-04 | 1.85E-04 | 1.27E-04 | 1.08E-04 | 1.01E-04 |
| | **2.50E-06** | 7.09E-05 | 1.16E-05 | 4.80E-06 | 3.60E-06 | 2.90E-06 |

B. SIAGE-GENE with the robust adjustment.

| | Alpha | Prev=0.01 | Prev=0.05 | Prev=0.1 | Prev=0.2 | Prev=0.5 |
|---|---|---|---|---|---|---|
| **SKAT-O** | **0.05** | 5.52E-02 | 5.06E-02 | 4.77E-02 | 4.39E-02 | 4.12E-02 |
| | **0.0001** | 3.14E-04 | 1.58E-04 | 1.16E-04 | 8.10E-05 | 5.51E-05 |
| | **2.50E-06** | 7.30E-06 | 4.51E-06 | 3.80E-06 | 1.54E-06 | 1.10E-06 |
| **SKAT-O-GCadj** | **0.05** | 5.00E-02 | 5.00E-02 | 4.77E-02 | 4.39E-02 | 4.12E-02 |
| | **0.0001** | 2.08E-04 | 1.51E-04 | 1.16E-04 | 8.10E-05 | 5.51E-05 |
| | **2.50E-06** | 3.90E-06 | 4.51E-06 | 3.80E-06 | 1.54E-06 | 1.10E-06 |
| **SKAT** | **0.05** | 6.59E-02 | 5.01E-02 | 4.49E-02 | 3.97E-02 | 3.63E-02 |
| | **0.0001** | 3.40E-04 | 1.53E-04 | 1.13E-04 | 6.76E-05 | 3.94E-05 |
| | **2.50E-06** | 8.20E-06 | 4.11E-06 | 2.90E-06 | 1.65E-06 | 5.01E-07 |
| **SKAT-GCadj** | **0.05** | 5.00E-02 | 5.00E-02 | 4.49E-02 | 3.97E-02 | 3.63E-02 |
| | **0.0001** | 1.01E-04 | 1.53E-04 | 1.13E-04 | 6.76E-05 | 3.94E-05 |
| | **2.50E-06** | 9.00E-07 | 4.11E-06 | 2.90E-06 | 1.65E-06 | 5.01E-07 |
| **Burden** | **0.05** | 3.95E-02 | 4.67E-02 | 4.66E-02 | 4.54E-02 | 4.42E-02 |
| | **0.0001** | 7.82E-05 | 8.58E-05 | 7.65E-05 | 7.19E-05 | 6.41E-05 |
| | **2.50E-06** | 1.80E-06 | 2.91E-06 | 2.00E-06 | 1.54E-06 | 2.10E-06 |

**Supplementary Table 14.** Empirical type I error rates for SAIGE-GENE in the simulation study with case-control sampling from an underlying large cohort. The liability scale heritability $h^2_{latent}$ =0.23.

| | | Case:Control=1:1 | | | Case:Control=1:9 | | |
|---|---|---|---|---|---|---|---|
| | alpha | Burden | SKAT | SKAT-O | Burden | SKAT | SKAT-O |
| SAIGE-GENE | **0.05** | 4.97E-02 | 4.86E-02 | 5.17E-02 | 4.87E-02 | 4.96E-02 | 5.16E-02 |
| | **0.0001** | 9.21E-05 | 9.05E-05 | 1.01E-04 | 9.07E-05 | 1.18E-04 | 1.27E-04 |
| | **2.50E-06** | 2.68E-06 | 1.56E-06 | 2.23E-06 | 2.37E-06 | 3.85E-06 | 2.07E-06 |

**Supplementary Table 15.** Empirical power for SAIGE-GENE and EmmaX-SKAT with two different percentages of causal variants (top vs bottom panels) and two different ratios of positive and negative effect directions (left vs right). $\beta$: effect size. $h^2$: heritability.

| $h^2$=0.4 | Proportion of causal variants= 0.4 | | | | | |
|---|---|---|---|---|---|---|
| | $\beta$ −/+=0.8/0.2 | | | $\beta$ −/+=1/0 | | |
| | Burden | SKAT | SKAT-O | Burden | SKAT | SKAT-O |
| EmmaX-SKAT | 54.80% | 80.20% | 84.20% | 89.10% | 82.1% | 92.00% |
| SAIGE-GENE | 54.60% | 81.30% | 84.00% | 89.00% | 83.20% | 91.70% |
| | Proportion of causal variants= 0.1 | | | | | |
| | $\beta$ −/+=0.8/0.2 | | | $\beta$ −/+=1/0 | | |
| | Burden | SKAT | SKAT-O | Burden | SKAT | SKAT-O |
| EmmaX-SKAT | 35.3% | 66.30% | 66.60% | 51.70% | 65.6% | 66.10% |
| SAIGE-GENE | 35.20% | 67.60% | 66.50% | 52.00% | 67% | 66.40% |

**Supplementary Table 16.** Empirical power for the SKAT-O test in SAIGE-GENE for binary phenotypes in cohort studies and case-control sampling studies. 40% of variants were simulated to be causal. 80% of causal variants were risk-increasing variants, while 20% were risk-decreasing variants. The liability scale heritability $h^2_{latent}$ =0.23. Note that the effect sizes in cohort study and case-control sampling were different, so the power in these two designs is not directly comparable.

| | Cohort study | | Case-control sampling | |
|---|---|---|---|---|
| case:control | Unadjusted SKAT-O | Robust SKAT-O | Unadjusted SKAT-O | Robust SKAT-O |
| 1:1 | 56.00% | 58.00% | 60.00% | 61.00% |
| 1:9 | 44.00% | 56.00% | 21.00% | 23.00% |
| 1:19 | 27.00% | 37.00% | 8.00% | 9.00% |
| 1:99 | 5.00% | 8.00% | | |
